# Combinatorial chromatin dynamics foster accurate cardiopharyngeal fate choices

**DOI:** 10.1101/546945

**Authors:** Claudia Racioppi, Keira A Wiechecki, Lionel Christiaen

## Abstract

During embryogenesis, chromatin accessibility profiles control lineage-specific gene expression by modulating transcription, thus impacting multipotent and fate-restricted states. Cardiac and pharyngeal/head muscles share a common origin in the cardiopharyngeal mesoderm, but the chromatin landscapes that govern mutlipotency and fate choices remain elusive. Here, we leveraged the simple chordate model *Ciona* to profile chromatin accessibility through stereotyped transitions from naive *Mesp*+ mesoderm to distinct fate-restricted heart and pharyngeal muscle precursors. An FGF-Foxf pathway acts in multipotent progenitors to establish lineage-specific patterns of accessibility, demonstrating extensive spatiotemporal decoupling between early enhancer accessibility and late cell-type-specific activity. We found that multiple *cis*-regulatory elements, with distinct accessibility profiles and motif compositions, are required to activate *Ebf* and *Tbx1/10*, two key determinants of cardiopharyngeal fate choices. We propose that these “combined enhancers” foster spatially and temporally accurate fate choices, by increasing the repertoire of regulatory inputs that control gene expression, through either accessibility and/or activity.

## Introduction

How a species’ genome encodes its diverse and specific biological features has fascinated generations of biologists, and answers regarding the genetic control of body plan, organ, tissue and cell type formation have emerged from steady progress in developmental biology. Cell types arise as cells divide and the progeny of pluripotent embryonic stem cells progress through multipotent and fate-restricted states. The ontogeny of diverse terminal cell identities involves differential expression of hundreds to thousands of genes. Their dynamic activities are orchestrated by complex gene regulatory networks, whereby DNA-binding proteins and co-factors act upon specific *cis*-regulatory elements to control gene expression (Davidson, 2010). Technical and conceptual revolutions in genome biology have extensively characterized the chromatin dynamics that govern the function of *cis*-regulatory elements (Klemm et al., 2019). Specifically, as the nuclear genome is packaged in nucleosomes, DNA-binding transcription factors compete with histones to interact with *cis*-regulatory elements and control gene expression. Thus, identifying changing landscapes of accessible chromatin governing the transition from multipotent to fate restricted progenitors offers privileged insights into the genomic code for progressive cell type specification.

Dynamic chromatin states underlying cardiomyocyte differentiation have been extensively profiled (Paige et al., 2012; Wamstad et al., 2012), and chromatin state regulation is essential for heart development (He et al., 2014; Rosa-Garrido et al., 2013; Zaidi et al., 2013). However, different parts of the heart originate from separate first and second fields of progenitor cells, including those referred to as cardiopharyngeal, which can also produce branchiomeric head muscles (Diogo et al., 2015; Lescroart et al., 2010). Bulk and single cell transcription profiling have begun to illuminate gene expression changes underlying cardiopharyngeal fate choices (Lescroart et al., 2018, 2014), but the corresponding chromatin dynamics remains largely elusive.

The tunicate *Ciona* emerged as a powerful chordate model to study early cardiopharyngeal development with high spatio-temporal resolution (Diogo et al., 2015; Kaplan et al., 2015). In *Ciona*, the cardiopharyngeal lineages arise from naive *Mesp+* mesodermal progenitors that emerge at the onset of gastrulation, and divide into two multipotent cardiopharyngeal progenitors (aka trunk ventral cells, TVCs) and two anterior tail muscles (ATMs), on either side of the embryo (Figure 1A). Following induction by FGF-MAPK signaling, cardiopharyngeal progenitors migrate collectively, before dividing asymmetrically and medio-laterally to produce small median first heart precursors (FHPs), and large lateral second trunk ventral cells (STVCs) (Davidson, 2005; Stolfi et al., 2010; Wang et al., 2013). The latter are also multipotent cardiopharyngeal progenitors, which upregulate *Tbx1/10* and then divide again to produce small median second heart precursors (SHPs), and large lateral atrial siphon muscle founder cells (ASMFs). ASMFs activate *Ebf,* which is necessary and sufficient to induce pharyngeal muscle specification (Razy-Krajka et al., 2014; Stolfi et al., 2014, 2010; Tolkin and Christiaen, 2016). Importantly, spatially and temporally accurate activation of *Tbx1/10* and *Ebf* in the STVC and ASMF, respectively, is essential to permit the emergence of all cardiopharyngeal cell lineages, as their ectopic expression would inhibit proper heart fate specification (Figure 1A).

**Figure 1.**
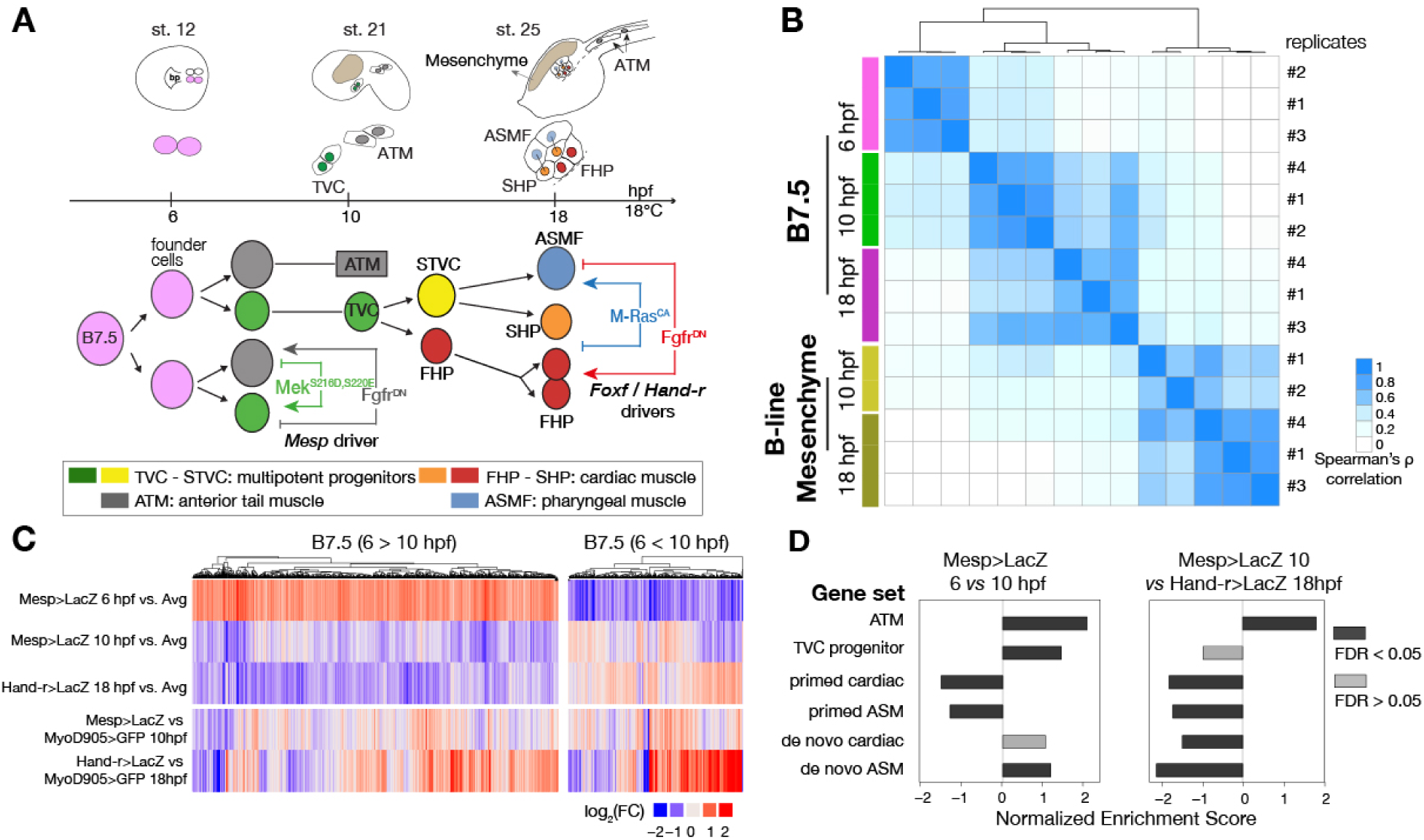
Profiling chromatin accessibility dynamics during early cardiopharyngeal cell development. (**A**) Embryos, larvae and lineage diagram showing B7.5 blastomeres, their cardiopharyngeal progeny, and the main stages sampled for ATAC-seq. Anterior tail muscle (ATM, gray), trunk ventral cell (TVC, green), secondary TVC (STVC, green), first heart precursor (FHP, red), second heart precursor (SHP, orange), atrial siphon precursors cell (ASMF, blue). Stages (St.) according to (Hotta et al., 2007) with hours post fertilization (hpf). (**B**) Spearman correlation of RPKM (reads per kb per million mapped reads) values in 14,178 regions changing accessibility over time or between B7.5 and B-line mesenchyme lineages. (**C**) Temporal changes in chromatin accessibility for 5,450 regions. “B7.5 6>10”: 3,691 regions more accessible at Mesp>LacZ 6 than Mesp>LacZ 10 hpf. “B7.5 6<10”: 1,759 regions more accessible at Mesp>LacZ 10 than Mesp>LacZ 6 hpf. The accessibility of these regions is shown for Mesp>LacZ 6, Mesp>LacZ 10, and Hand-r>LacZ 18 hpf vs. the average (avg) accessibility in the control cells. Cell-type specific chromatin accessibility is shown in the comparison of Mesp>LacZ and MyoD905>GFP at 10 and Hand-r>LacZ and MyoD905>GFP18 hpf. (**D**) Gene Set Enrichment Analysis (GSEA) normalized enrichment score of defined gene sets in regions ranked by difference in accessibility between time points as indicated (see Supplementary Text).

Building on previous extensive transcription profiles (Christiaen et al., 2008; Razy-Krajka et al., 2014), including single cell RNA-seq (scRNA-seq) from multipotent cardiopharyngeal progenitors to first and second heart lineages and pharyngeal muscle precursors (Wang et al., 2019), here we characterized the genome-wide chromatin accessibility dynamics underlying cardiopharyngeal fate specification. We identified regulatory inputs that govern *cis*-regulatory element accessibility and activity, as well as cell-type-specific enhancers for key cardiopharyngeal determinants. We found that, in multipotent progenitors, an FGF-Foxf pathway controls cardiopharyngeal-specific patterns of accessibility, which govern later heart vs. pharyngeal muscle-specific expression profiles. We further characterized temporal patterns of chromatin accessibility during cardiopharyngeal development. In particular, activation of fate determinants *Tbx1/10* and *Ebf* specifically in the STVCs and ASMF, respectively, require multiple *cis*-regulatory elements with distinct spatio-temporal patterns of accessibility, which precede gene expression. We propose that these elements function as “combined enhancers”, which mediate distinct inputs, including from determinants of chromatin accessibility, to regulate gene activation. The observation that *cis*-regulatory inputs from multiple elements control expression of a single gene is consistent with the “shadow-” and “super-enhancer” paradigms (Barolo, 2012; Hnisz et al., 2013; Hong et al., 2008; Kvon et al., 2014; Perry et al., 2011, 2010; Pott and Lieb, 2015; Whyte et al., 2013). However, while shadow enhancer promotes robust transcription through the actions of multiple elements mediating similar regulatory inputs (Frankel, 2012; Frankel et al., 2010; Lam et al., 2015; Zeitlinger et al., 2007), we propose that combined enhancers promote spatially and temporally accurate fate choices, by augmenting the repertoire of *trans*-acting inputs controlling gene activation through enhancer activity and/or chromatin accessibility.

## RESULTS

### A reference accessome for cardiopharyngeal development

To characterize the chromatin landscape underlying early cardiopharyngeal development, we used the assay for transposon-accessible chromatin (ATAC-seq; (Buenrostro et al., 2013)) on lineage-specific samples isolated at successive time points, and following defined perturbations (Figure 1A; Table S1; (Razy-Krajka et al., 2018a; Wang et al., 2019)). Using the B7.5 lineage-specific *Mesp>tagRFP* reporter (Wang et al., 2018), we used FACS to collect ~4,000 cells per biological replicate from embryos dissociated at five time points encompassing key transitions in cardiopharyngeal development (Figure 1A): naive *Mesp*+ mesoderm (aka founder cells; (Cooley et al., 2011), ATMs, TVCs, STVCs as well as fate-restricted first and second heart precursors (FHPs and SHPs), and pharyngeal muscle precursors (aka atrial siphon muscle founder cells - ASMF- and their progeny, the ASM precursors- ASMP; (Razy-Krajka et al., 2014)). The latter fate-restricted progenitors were obtained from larvae dissociated at three time points (15, 18 and 20 hours post-fertilization, hpf; Figure 1A). From the same embryonic cell populations, we used a co-transfected *MyoD905>GFP* reporter to isolate B-line mesenchymal cells (Christiaen et al., 2008). In the present study, we predominantly focused our analysis on the cardiopharyngeal progenitors at 6, 10 and 18 hpf (Figure 1A).

We obtained ~500 million unique ATAC-seq reads, with fragment-size distributions showing the characteristic ~150 bp periodicity and patterns of mono-, di- and tri-nucleosomal fragments (Buenrostro et al., 2013), which were absent in the genomic DNA control (Figure 1—figure supplement 1A). We identified ATAC-seq peaks using MACS2 (Zhang et al., 2008), and generated a combined atlas of 56,090 unique and non-overlapping accessible regions covering 9.25% of the *C. robusta* genome, which we used as our reference “accessome” (Figure 1—figure supplement 1B,E). General metrics including peak numbers, size, GC content and genomic distribution were comparable to consensus peaksets reported in other studies of chromatin accessibility in developmental contexts (Materials & Methods; Figure 1—figure supplement 1; (Daugherty et al., 2017; Hockman et al., 2018; Janes et al., 2018; Li et al., 2007; Madgwick et al., 2018)).

Next, we annotated the reference accessome by associating accessible regions with other genomic features, especially gene models. In *Ciona*, the transcripts of approximately half of the protein-coding genes undergo spliced-leader (SL) *trans*-splicing, causing the 5’ end of mRNAs to differ from the transcription start site (TSS) (Vandenberghe et al., 2001). Using annotated TSSs (Satou et al., 2006; Vandenberghe et al., 2001; Yokomori et al., 2016), RNA-seq datasets (Wang et al., 2019), and our ATAC-seq data (Figure 1—figure supplement 2C), we determined that promoter regions and 5’ untranslated regions (5’UTR) were over-represented in the accessome (*p*<0.001, binomial test; Figure 1—figure supplement 1C), and we detected nucleosome footprints immediately upstream of TSSs, consistent with a tendency for constitutive accessibility (Figure 1—figure supplement 2A,B; (T. N. Mavrich et al., 2008; Travis N. Mavrich et al., 2008). By contrast, intronic and intergenic regions were significantly under-represented in our reference accessome, compared to the whole genome, although they were the most abundant elements (32.8% and 20.8%, respectively; Figure 1—figure supplement 1B). This suggests that most of these elements are accessible in specific contexts, as expected for tissue-specific *cis*-regulatory elements (Long et al., 2016).

We associated annotated genes with ATAC-seq peaks located within 10 kb of the TSS or transcription termination site (TTS) (Figure 1—figure supplement 3A) (Brozovic et al., 2018), thus assigning median values of 11 peaks per gene, and 3 genes per peak, owing to the compact *Ciona* genome (Figure 1—figure supplement 3B,C). Notably, active regulatory genes encoding transcription factors and signaling molecules were associated with significantly more peaks than other expressed genes (*p* < 0.001, binomial test; Figure 1—figure supplement 3F). This high peak density surrounding regulatory genes is reminiscent of previously described super-enhancers (Whyte et al., 2013) and Clusters of Open *Cis*-Regulatory Elements (COREs) surrounding developmental regulators (Gaulton et al., 2010; Khan et al., 2018; Pott and Lieb, 2015).

### Cardiopharyngeal accessibility profiles are established in multipotent progenitors

Using this reference accessome, we investigated lineage-specific and dynamic patterns of chromatin accessibility during fate decisions. We observed the greatest contrast in accessibility between the B7.5 and B-line mesenchyme lineages, with biological replicates correlating most highly (Spearman’s ρ > 0.93), indicating reproducible detection of extensive lineage-specific accessibility (Figure 1B). Within the B7.5 lineage, correlation analysis suggested that most changes occur between 6 and 10 hpf, during the transition from naive *Mesp*+ mesoderm to multipotent cardiopharyngeal progenitors (TVCs). Higher correlation between multipotent progenitors and mixed heart and pharyngeal muscle precursors, obtained from 18 hpf larvae, suggested more stable accessibility profiles during and immediately following early cardiopharyngeal fate choices (Figure 1B). Consistent with correlation analyses, most significant temporal changes in accessibility occurred during the transition from naive *Mesp+* mesoderm to multipotent progenitors (5,450 regions, FDR < 0.05; Figure 1C). Specifically, about two thirds (64.7%, 3,525/5,450) of these regions showed reduced accessibility at 10 hpf, in multipotent progenitors, compared to 6 hpf naive *Mesp*+ mesoderm (Figure 1C). Conversely, 1,252 regions become accessible between 6 and 10 hpf or later, and 38.8% (486/1,252) of these regions were more accessible in the B7.5-lineage compared to the mesenchyme (Figure 1C). Moreover, the subset of regions opening between 6 and 10 hpf or later was enriched in genomic elements associated with cardiopharyngeal markers, including primed pan-cardiac and pharyngeal muscle markers, while elements flanking tail muscle markers (ATMs) or multipotent progenitor-specific genes were predominantly closing between 6 and 10 to 18 hpf (Figure 1D; see Supplementary text for definition of gene sets). Taken together, these observations suggest that cardiopharyngeal accessibility profiles are established specifically in the B7.5 lineage, upon induction of multipotent progenitors, and persist in fate-restricted cells.

To further analyze changes in accessibility associated with multipotent progenitor induction, we performed ATAC-seq on B7.5 lineage cells isolated at 10 hpf following defined perturbations of FGF-MAPK signaling: a constitutively active form of Mek (Mek^S216D,S220E^), which converts all B7.5 lineage cells into multipotent cardiopharyngeal progenitors or a dominant negative form of the Fgf receptor (Fgfr^DN^), which blocks induction and transforms all B7.5-derived cells into ATMs (Figure 1A; (Davidson et al., 2006; Razy-Krajka et al., 2018a)). We used DESeq2 (Love et al., 2014) to compute differential accessibility of the elements in the reference accessome, and identified 2,728 and 2,491 differentially accessible regions following either inhibition or activation of FGF-MAPK signaling, respectively (Figure 2A,B; Figure 2—figure supplement 1A). Using peak-to-gene annotations (Figure 1—figure supplement 3A), we cross-referenced ATAC-seq with expression microarray data obtained from B7.5 lineage cells expressing the same Fgfr^DN^ (Christiaen et al., 2008), and observed a positive correlation between changes in differential accessibility and differential gene expression at 10 hpf (Spearman’s ρ = 0.47; Figure 2A; Figure 2—figure supplement 2A). Specifically, 48% of FGF-MAPK-regulated genes were associated with at least one element showing consistent differential accessibility, including 260 candidate FGF-MAPK-activated TVC markers associated with 557 regions predicted to open specifically in multipotent cardiopharyngeal progenitors at 10 hpf (Table S2, Figure 2—figure supplement 2B). Conversely, the majority (603 regions) of differentially accessible ATAC-seq peaks associated with 263 FGF-MAPK-inhibited tail muscles markers, and were also more accessible upon inhibition of FGF signaling (Table S2; Figure 2—figure supplement 1A). Taken together, these observations indicate that cardiopharyngeal accessibility profiles are established in the multipotent progenitors by opening regions associated with genes upregulated upon induction by FGF-MAPK signaling.

**Figure 2.**
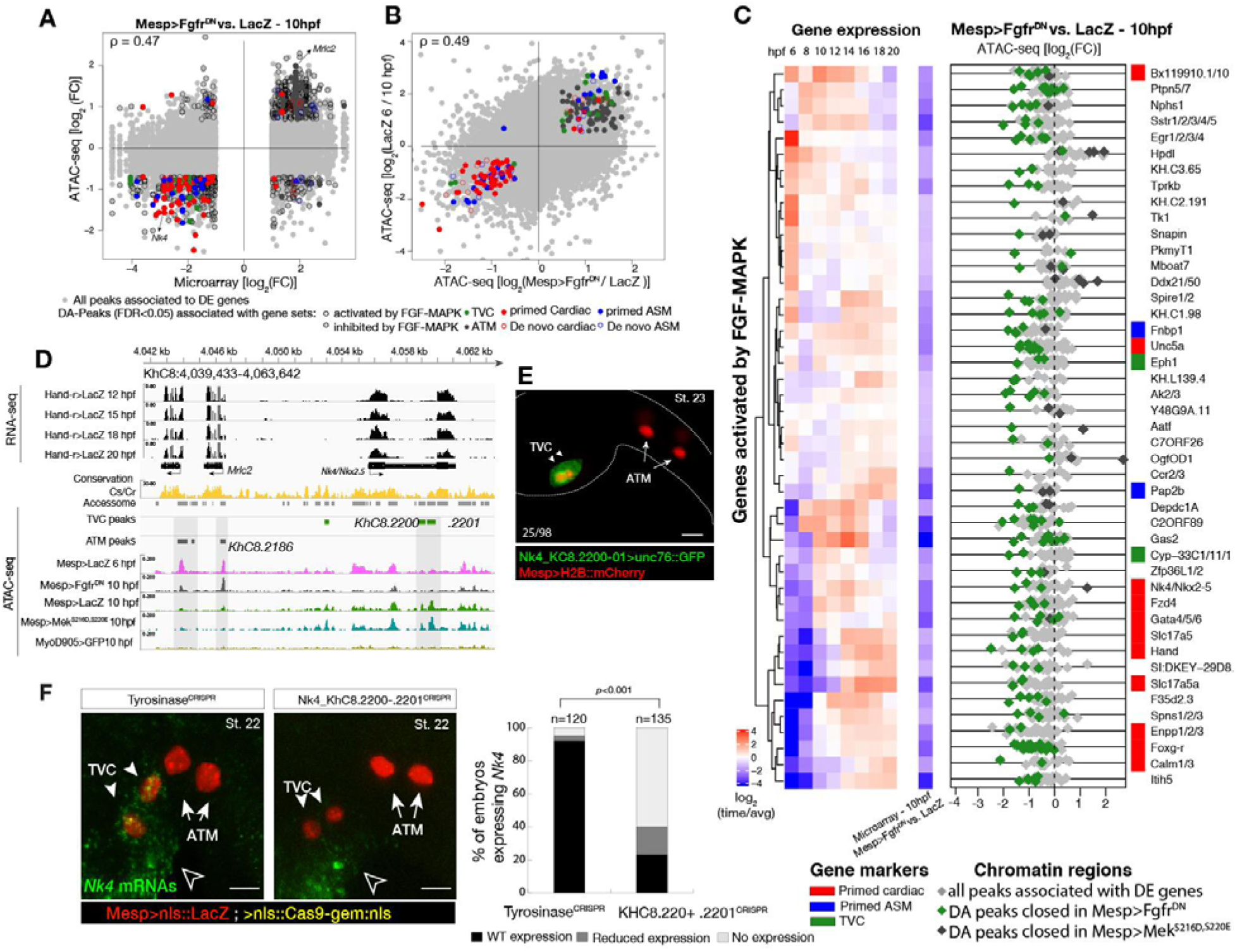
Cardiopharyngeal accessibility profiles are established in multipotent progenitors. (**A-B**) Correlations between differential gene expression (DE) and differential chromatin accessibility (DA) in response to FGF-MAPK perturbation in the multipotent progenitors (A) and between chromatin accessibility in response to FGF-MAPK perturbation and in multipotent progenitors (10hpf) versus founder cells (6hpf). (B). Colored dots are DA peaks associated to cell type-specific DE genes. D is the Spearman correlation of expression and accessibility for DA regions associated to DE genes (A) or of region response to MAPK perturbation with accessibility in founder cells versus multipotent progenitors (B). (**C**) Relationship between expression and accessibility of DE genes associated to DA regions for genes in the bottom 0.75% quantile of fold change between expression in Fgfr^DN^ and control (log_2_FC < −1.32). Microarray log_2_ (fold change (FC)) values are shown on the left. The fold change for all time points is versus the average. (**D**) A 24 kb region on chromosome 8 displaying expression (RNA-seq) and chromatin accessibility (ATAC-seq; normalized by total sequencing depth). Gray shaded boxes show validated ATM-specific promoters and a newly identified TVC-specific enhancer in *Nk4/Nkx2-5* intron. (**E**) Enhancer driven *in vivo* reporter expression (green) of tested ATAC-seq regions (KhC8.2200 and .2201). TVCs marked with Mesp>H2B::mCherry (red). Numbers indicate observed/total of half-embryos scored. (**F**) Endogenous expression of *Nk4/Nkx2-5* visualized by *in situ* (green) in Tyrosinase^CRISPR^ and upon CRISPR/Cas9-induced deletions of TVC-specific region. Nuclei of B7.5 lineage cells are labelled by *Mesp>nls::LacZ* and revealed with an anti beta-galactosidase antibody (red). *Nk4/Nkx2-5* expression was not affected in the epidermis (open arrowhead). Experiment performed in biological replicates. Scale bar, 20 μm. Fisher exact test, total numbers of individual halves scored per condition are shown in ’n=’. Gene expression data for 6 hpf and “FGF-MAPK perturbation 10 hpf” (Christiaen et al. 2008) and 8 to 20 hpf (Razy-Krajka et al. 2014) were previously published.

Consistent with the hypothesis that FGF-MAPK-dependent, cardiopharyngeal-specific elements act as tissue-specific enhancers, they were predominantly found in intronic or intergenic regions (48% and 37%, respectively, FDR < 0.05, Hypergeometric test). Conversely, tissue-specific peaks associated with tail muscle markers were enriched in promoters, TSS and 5’UTR (Hypergeometric test, FDR < 0.05, 57%, 23% and 15%, respectively; Figure 2—figure supplement 2C). Previously characterized enhancers for TVC-specific genes *Lgr4/5/6*, *Rhodf*, *Foxf*, *Unc5*, *Rgs21, Ddr, Asb2* and *Gata4/5/6* showed ATAC-seq patterns consistent with cardiopharyngeal-specific accessibility (Table S3; (Beh et al., 2007; Bernadskaya et al., 2019; Christiaen et al., 2008; Woznica et al., 2012)). We thus leveraged differential accessibility profiles to identify novel enhancers of cardiopharyngeal gene expression. We focused on a locus containing the conserved cardiac determinant and TVC marker, *Nk4/Nkx2-5 (Wang et al., 2013)*, and two tail-muscle specific *Myosin regulatory light chain* (*Mrlc*) genes (Kusakabe et al., 2004; Satou et al., 2001b; Sierro et al., 2006), with associated elements showing the predicted TVC- and ATM-specific accessibility patterns, respectively (Figure 2A,D). Reporter gene expression assays showed that a DNA fragment containing differentially accessible elements located in the *Nk4/Nkx2-5* intron (*KhC8.2200* and *.2201*) was sufficient to drive *GFP* expression specifically in cardiopharyngeal multipotent progenitors (Figure 2E). B7.5 lineage-specific CRISPR/Cas9-induced deletions of these elements reduced or eliminated *Nk4/Nkx2-5* expression specifically in TVCs, thus demonstrating its role as a *bona fide* cardiopharyngeal enhancer (Figure 2F; Figure2—figure supplement 5D). Extending these analyses to other loci, including *Fgf4*, *Fzd4*, *Foxg-r*, *Fbln*, *Eph1, Ncaph, Hand* and *Smurf1/2*, we identified 8 out of 15 candidate cardiopharyngeal enhancers that drove reporter expression in the multipotent progenitors (Figure 2—figure supplement 3; Table S4), and B7.5-lineage-specific CRISPR/Cas9-mediated mutagenesis targeting differentially accessible elements reduced TVC-specific expression of the neighbouring genes *Fgf4*, *Smurf1/2* and *Fbln* (Figure 2—figure supplement 4; Figure 2—figure supplement 6; Table S5). Conversely, candidate ATM-specific elements activated reporter gene expression in the tail muscles, including ATM cells, but not in the cardiopharyngeal progenitors, and were located near tail muscle markers (Figure 2—figure supplement 1B,C; Table S6). Collectively, these findings indicate that genomic elements that open specifically in multipotent progenitors act as transcriptional enhancers of cardiopharyngeal gene expression and their accessibility is controlled by FGF-MAPK induction.

### A Foxf-dependent code for cardiopharyngeal accessibility

Next, we harnessed chromatin accessibility patterns predictive of cardiopharyngeal enhancer activity to identify enriched sequence motifs, and thus candidate regulators of chromatin accessibility and gene expression. For this, we performed a one-tailed hypergeometric test for enrichment of known motifs. We complemented this analysis by calculating differential accessibility of motifs using chromVAR (Schep et al., 2017), which was developed to analyze sequence motifs associated with cell-type-specific accessibility (Figure 3A). Naive *Mesp+* mesoderm-specific elements, which closed between 6 and 10 hpf, were enriched in motifs for Homeodomain, T-box and Ets families of transcription factors (TF), consistent with documented roles for Lhx3/4, Tbx6 and Ets homologs in B7.5 blastomeres (Figure 3A; (Davidson et al., 2006, 2005; Satou et al., 2001a). Candidate tail muscle-specific elements, which opened upon Fgfr^DN^ misexpression, were similar to naive *Mesp*+ mesoderm, and enriched in motifs for the basic helix-loop-helix (bHLH) family of TFs, which includes Mesp and Mrf/MyoD, a conserved muscle-specific transcription regulator that promotes tail muscle differentiation (Christiaen et al., 2008; Meedel et al., 2007; Razy-Krajka et al., 2014; Tolkin and Christiaen, 2016). By contrast, motifs for Zinc Finger, Fox/Forkhead and nuclear receptor families of TFs were enriched among candidate cardiopharyngeal-specific elements, revealing a typical mesendodermal signature for early cardiopharyngeal progenitors (Cusanovich et al., 2018).

**Figure 3.**
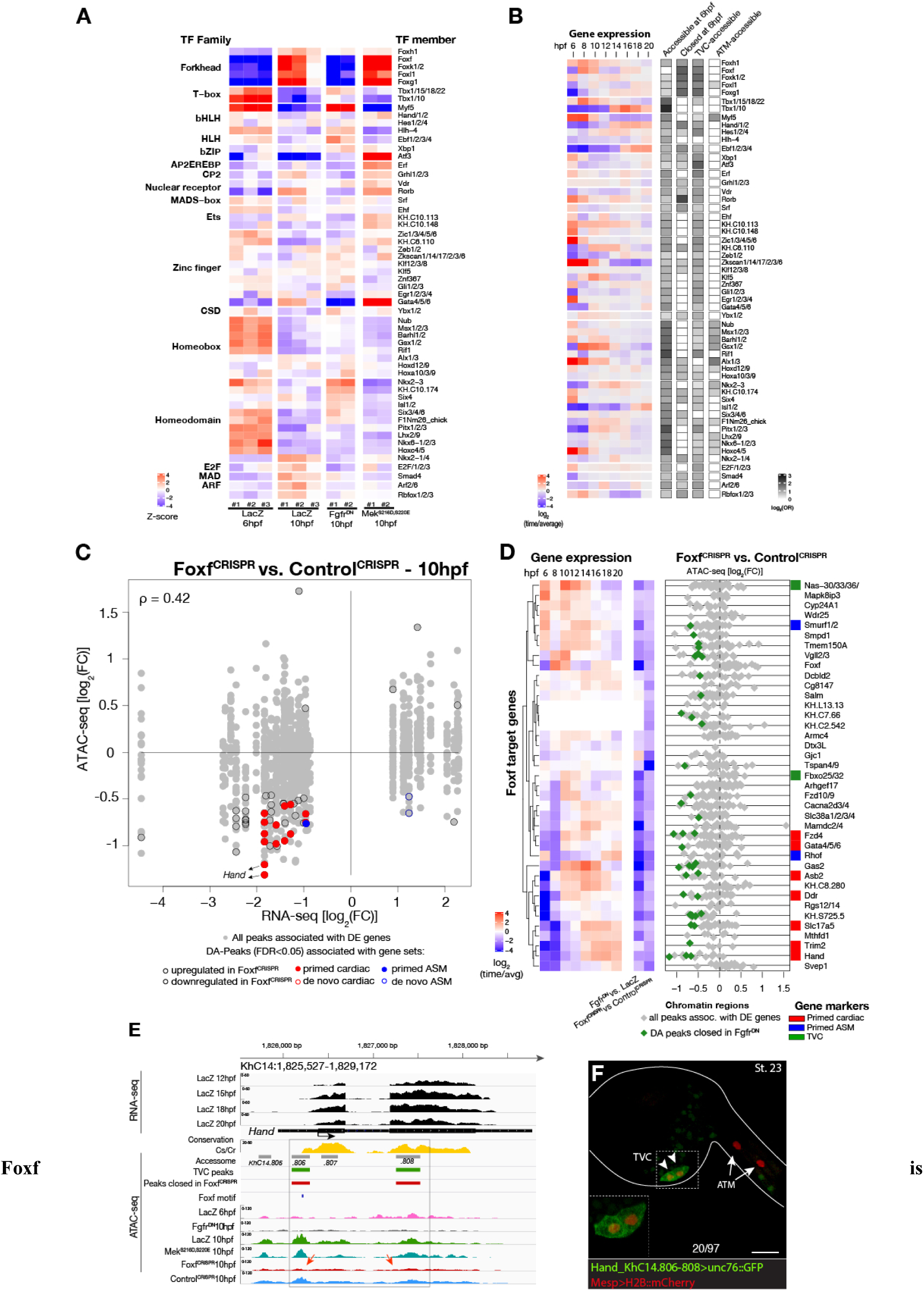
required for cardiopharyngeal-specific chromatin accessibility. (**A**) Motif accessibility between libraries from ChromVAR (Schep et al., 2017). Motifs were obtained and associated to *Ciona* transcription factors (TFs) as described in the Supplementary Text. Deviations were computed for FGF signaling-dependent regions at 10 hpf and B7.5 replicates at 6 and 10 hpf. We calculated the differential accessibility of all motifs between conditions and time points. Only the most significant motif is shown for each TF. (**B**) Expression of transcription factors over time compared to enrichment of corresponding TF motifs in condition specific peak sets. log_2_ odds ratio values (see Materials and Methods) are shown for motifs that are significantly enriched in a peak set (one-tailed hypergeometric test, FDR < 0.05). Only TFs expressed in the B7.5 lineage are shown. (**C**) Differential expression of Foxf target genes (DE) vs. differential chromatin accessibility (DA) in _Foxf_CRISPR. is the Spearman correlation of expression and accessibility for DA regions associated to DE genes. (**D**) Association between expression of Foxf target genes and accessibility of proximal regions which were both TVC-specific and closed in Foxf^CRISPR^ as in Figure 2C. (**E**) A 3.6 kb region on chromosome 14 displaying expression profiles of RNA-seq and chromatin accessibility profiles of ATAC-seq normalized tag count. Foxf core binding site (GTAAACA) is displayed as blue line. The boxed region indicates a newly identified TVC-specific enhancer in *Hand* locus. Red arrow indicates a TVC-specific enhancer showing closed chromatin in Foxf^CRISPR^ ATAC-seq. (**F**) Enhancer driven *in vivo* reporter expression (green) of tested ATAC-seq peaks. TVCs marked with Mesp>H2B::mCherry (red). Numbers indicate observed/total of half-embryos scored. Experiment performed in biological replicates. Scale bar, 30 μm. Gene expression data for 6hpf and “FGF-MAPK perturbation 10hpf” (Christiaen et al., 2008), and from 8 to 20hpf (Razy-Krajka et al., 2014) were previously published.

Combined with motif enrichment analyses, temporal gene expression profiles (Razy-Krajka et al., 2014) identified candidate *trans*-acting regulators of cardiopharyngeal-specific accessibility and/or activity (Figure 3B). For example, regions specifically accessible in tail muscle- or naive *Mesp+* mesoderm were enriched in homeobox, Ets and T-box motifs, consistent with early expression of *Mrf/MyoD*, *Lhx3/4*, *Ets1/2*, and *Tbx6*, respectively. Similarly, the increased accessibility of motifs for K50 Paired homeodomain proteins in naive *Mesp+* mesoderm indicated a possible role for *Otx*, which is expressed early in B7.5 blastomeres (Figure 3A,B; (Hudson, 2003)). Cardiopharyngeal-specific enrichment for Fox/Forkhead and Zinc Finger motifs pointed to several known factors, including *Foxf*, one of the first genes activated in multipotent progenitors upon induction by FGF-MAPK, prior to *Gata4/5/6 (Beh et al., 2007; Christiaen et al., 2008; Ragkousi et al., 2011)*. Moreover, GATA and Forkhead proteins are founding members of a group of TFs known as pioneers, which can bind their target sites in closed chromatin and promote accessibility (Cirillo et al., 2002; Zaret and Carroll, 2011). Protein sequence alignments indicated the presence of key residues, conserved between the DNA binding domains of Foxf and the classic pioneer FOXA, which mimic linker histone H1 in its ability to displace DNA-bound nucleosomes (Figure 3—figure supplement 1B,C; (Clark et al., 1993)). Finally, the *Foxf* enhancer was accessible in naive *Mesp*+ founder cells, suggesting that it is poised for activation, unlike the intronic *Gata4/5/6* enhancer (Figure 3—figure supplement 2A). Consistent with a role for Fox and GATA proteins in opening and activating the *Nk4/Nkx2-5* enhancer, we found putative cognate binding sites in the newly identified element conserved with the closely related species, *C. savignyi* (Figure 3—figure supplement 3B,C). Taken together, these analyses identified a putative code for cardiopharyngeal-specific accessibility and enhancer activity, which comprise motifs for candidate DNA binding factors of the Forkhead and GATA and identified *Foxf* as candidate determinant of cardiopharyngeal accessibility.

To test if Foxf contributes to establishing cardiopharyngeal accessibility and gene expression profiles, we used reagents for B7.5 lineage-specific loss-of-function by CRISPR/Cas9-mediated mutagenesis (Foxf^CRISPR^; (Gandhi et al., 2017)), and performed ATAC- and RNA-seq on FACS-purified cells isolated from tailbud embryos at 10 hpf. RNA-seq confirmed that CRISPR/Cas9 mutagenesis inhibited *Foxf* itself and other TVC-expressed genes, including effectors of collective cell migration such as *Ddr*, consistent with previous microarray data (Bernadskaya et al., 2019; Christiaen et al., 2008) (Figure 3D; Figure3—figure supplement 2B). Out of 52 differentially expressed genes (Figure 3—figure supplement 2B; Table S7), seven down-regulated genes were previously annotated as primed pan-cardiac markers, including *Hand*, *Gata4/5/6* and *Fzd4 (Wang et al., 2019)*. Down-regulated genes also included primed pharyngeal muscle markers, such as *Rhod/f* (Figure 3D; Figure 3—figure supplement 2B; (Christiaen et al., 2008; Razy-Krajka et al., 2014)), suggesting that Foxf promotes the onset of both the cardiac and pharyngeal muscle programs in multipotent progenitors, a feature known as multilineage transcriptional priming (Razy-Krajka et al., 2014; Wang et al., 2019).

Consistent with the effects of Foxf mutagenesis on gene expression, regions closed in *Foxf^CRISPR^* samples included known cardiopharyngeal enhancers for *Gata4/5/6* and *Ddr* (Figure 3D; Figure3—figure supplement 4; (Bernadskaya et al., 2019; Christiaen et al., 2008; Woznica et al., 2012)), newly identified enhancers for *Eph1, Smurf1/2* and *Fzd4*, and a novel enhancer of *Hand* expression (*Hand*_*KhC14.805* - *.807*; Figure 3C-F, Table S8). These differentially accessible elements contain several, evolutionary conserved, putative Fox binding sites (Figure 3—figure supplement 5; Figure 3—figure supplement 6). We identified two conserved putative Forkhead binding sites in the minimal STVC-specific enhancer from the *Tbx1/10* locus (termed T12, (Razy-Krajka et al., 2018a), which were necessary for reporter gene expression (Figure 5 — figure supplement 2 D,E). Moreover, loss-of-function of Foxf (Foxf^CRISPR^) drastically reduced T12 enhancer activity (Figure 5 —figure supplement 2). These results are consistent with the hypothesis that Foxf acts directly on the minimal *Tbx1/10* enhancer to promote its activity in the second multipotent cardiopharyngeal progenitors.

Notably, 98% (40/41) of the regions with diminished accessibility following Foxf inhibition, and located near a candidate Foxf target gene, were also accessible in multipotent progenitor cells (Figure 3D; Figure 3—figure supplement 7A). Moreover, 22% (600/2,728) of the predicted multipotent progenitor-specific elements were closed upon Foxf inhibition, and gene set enrichment analysis indicated that *Foxf* loss-of-function generally decreased the accessibility of cardiopharyngeal-specific elements (Figure 3—figure supplement 2C; Figure3—figure supplement 7B). Finally, 18 of 41 (44%) of the Foxf-dependent elements associated with candidate Foxf targets were closed in 6 hpf founder cells and appear to open specifically in the cardiopharyngeal progenitors by 10 hpf (Figure 3—figure supplement 2D; Figure 3—figure supplement 7 B,C; Table S9). This dynamic is consistent with a requirement for Foxf activity following its activation in the TVCs, immediately after division of the naive *Mesp*+ progenitors. Taken together, these results indicate that, in newborn multipotent progenitors, FGF-MAPK signaling upregulates *Foxf* (Beh et al., 2007; Christiaen et al., 2008), which is in turn required to open a substantial fraction of cardiopharyngeal-specific elements for gene expression in multipotent progenitors, including for such essential determinants as *Gata4/5/6* and *Hand* (Figure 4H).

**Figure 4.**
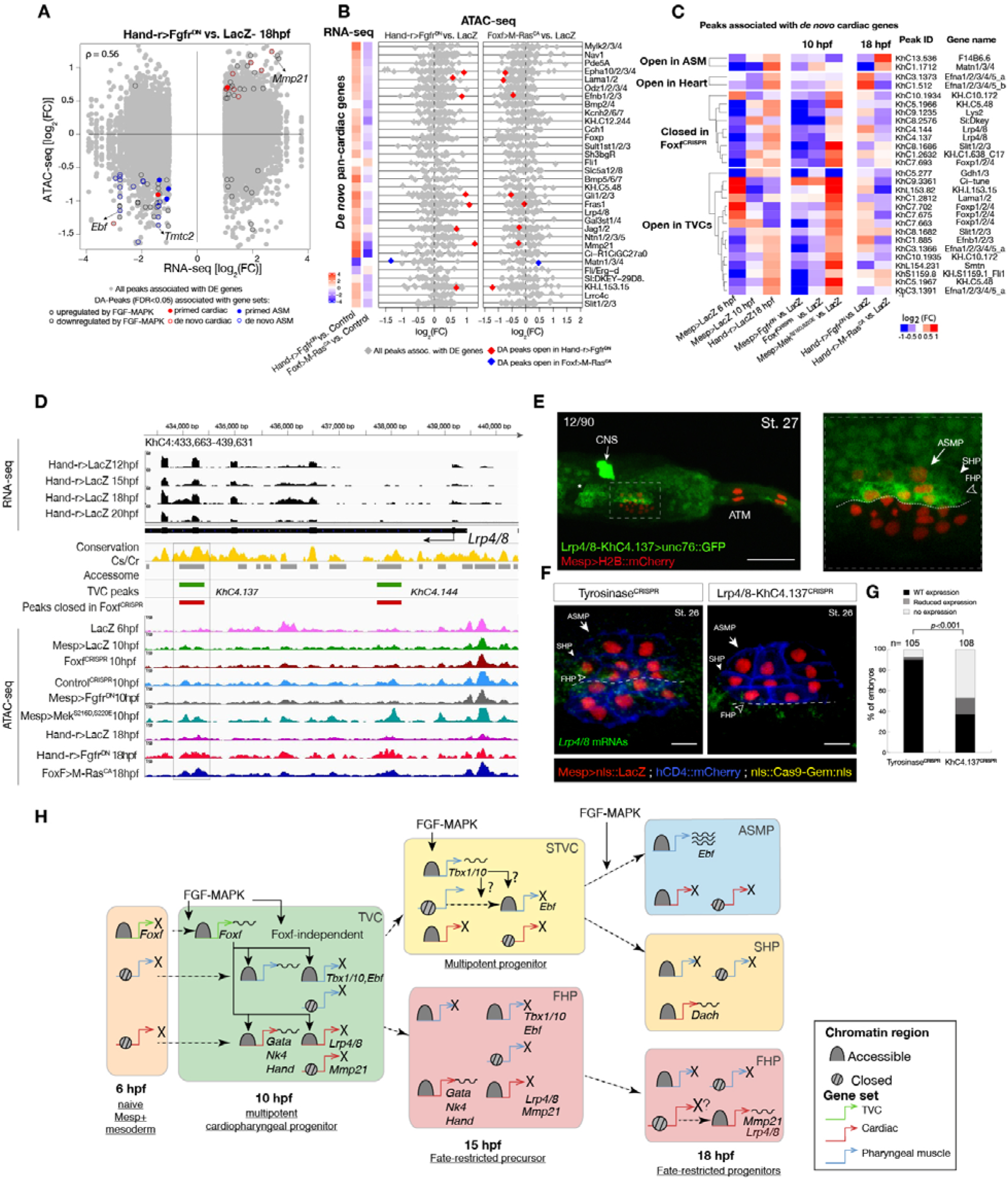
Cardiopharyngeal lineage-specific accessibility profiles and decoupling between enhancer accessibility and activity for *de novo* expressed genes. (**A**) Differentially expressed (DE) genes vs differentially accessible (DA) peaks in response to FGF-MAPK perturbation in the fate-restricted cells. □ is the Spearman correlation of expression and accessibility for DA peaks associated to DE genes. (**B**) Relationship between accessibility and expression of *de novo* pan-cardiac genes as in Figure 2C. DE genes in either condition are shown on the left. (**C**) Time-dependent ATAC-seq peaks associated to *de novo* expressed pan-cardiac genes. The accessibility of these peaks is shown for 6, 10 and 18 hpf vs. the average accessibility in the controls (LacZ) and upon FGF-MAPK perturbations at either 10 or 18hpf. Peaks were classified as “Open in ASM” (less accessible in Fgfr^DN^ vs. M-Ras^CA^ or LacZ at 18hpf), “Open in Heart” (less accessible in M-Ras^CA^ vs. Fgfr^DN^ or LacZ at 18hpf), “Closed in Foxf^CRISPR^” (less accessible in Foxf^CRISPR^ vs. Control^CRISPR^), or “Open in TVC” (less accessible in Fgfr^DN^ vs. Mek^S216D,S220E^ or LacZ at 10 hpf). Only regions changing accessibility between 6 and 10 hpf, or 10 and 18 hpf are shown. (**D**) A 6 kb region on chromosome 4 displaying expression profiles of RNA-seq and chromatin accessibility profiles of ATAC-seq normalized tag count. Peak ID refers to elements tested for reporter assay *in vivo*. The newly identified enhancer in *Lrp4/8* locus is in the boxed region. (**E**) Enhancer driven *in vivo* reporter expression (green) of tested “*KhC4.137*” peak. TVCs marked with *Mesp>H2B::mCherry* (red). Numbers indicate observed/total of half-embryo scored. Zoom on cardiopharyngeal cell lineage (panel on the right). (**F**) Endogenous expression of *Lrp4/8* visualized by *in situ* (green) in Tyrosinase^CRISPR^ and upon CRISPR/Cas9-induced deletion of ATAC-seq peaks. Nuclei of B7.5 lineage cells are labelled by *Mesp>nls::LacZ* and revealed with an anti beta-galactosidase antibody (red). Mesp driven hCD4::mCherry accumulates at the cell membrane as revealed by anti mCherry antibody (Blue). Experiment performed in biological replicates. Scale bar=10 μm. (**G**) Fisher exact test; n is the total number of individual embryo halves scored per condition. (**H**) Summary model: patterns of chromatin accessibility dynamics and gene expression during early cardiopharyngeal fate specification.

### Chromatin accessibility in late heart vs. pharyngeal muscle precursors

Besides controlling coherent chromatin opening, enhancer activity and gene expression in multipotent cardiopharyngeal progenitors, FGF-Foxf inputs also appeared to open regions associated with later *de novo*-expressed heart and pharyngeal muscle markers (Figure 2A,B; Table S10; (Razy-Krajka et al., 2018a, 2014; Wang et al., 2019, 2013)). Accessibility patterns were also better correlated between 10 and 18 hpf (Figure 1B), suggesting a decoupling between early accessibility and late heart-vs. pharyngeal muscle-specific expression in late fate-restricted precursors. To identify accessibility patterns underlying the heart vs. pharyngeal muscle fate choices, we compared bulk RNA-seq (Wang et al., 2019) and ATAC-seq datasets obtained from cardiopharyngeal lineage cells isolated from 18 hpf larvae, following the same defined perturbations of FGF-MAPK signaling (Figure 1A; Figure 4A; Figure 4—figure supplements 1A-D; 2A,B; (Davidson et al., 2006; Razy-Krajka et al., 2018a)). Among cardiac and pharyngeal muscle markers, we identified 35 FGF-MAPK-regulated genes associated with one or more elements showing consistent differential accessibility (Figure 4A,B; Figure 4—figure supplement 2A,B; Table S11). This indicated that, at least for a subset of cardiopharyngeal marker genes, FGF-MAPK-dependent changes in gene expression follow corresponding changes in chromatin accessibility in early heart and pharyngeal muscle precursors.

Gene-level inspection of differential accessibility associated with either inhibition or activation of gene expression revealed that only a fraction of associated elements was either closing or opening upon perturbation of FGF-MAPK signaling (Figure 4B; Figure 4—figure supplement 2B). For example, the first heart lineage marker *<u>M</u>atrix <u>m</u>etallo<u>p</u>roteinase 21*/*Mmp21* (Wang et al., 2019) was associated with multiple upstream and intronic elements, but only some of these elements were differentially accessible following either gain or loss of FGF-MAPK function (Figure 4—figure supplement 2A-C), and a 3kb fragment containing the upstream differentially accessible element sufficed to drive reporter gene expression throughout the cardiopharyngeal lineage, but not specifically in the first heart precursors (Figure 4—figure supplement 2C-E). Similarly, reporter gene expression assays showed that DNA fragments containing differentially accessible elements located ~0.5 kb upstream of the coding region of the *de novo*-expressed gene *Tmtc2* (*KhC2.3468*), and upstream of *KH.C1.1093_ZAN* (*KhC3.47, KhC3.46*), were sufficient to drive *GFP* expression in both cardiac and pharyngeal muscle progenitors, consistent with the notion that electroporated plasmids are not “chromatinized” and thus constitutively accessible (Figure 4—figure supplement 2 F-G; Figure 4—figure supplement 3 C-D). This suggested that, for genes like *Mmp21, Tmtc2* and *Zan*, cell-type-specific accessibility determines cardiac vs. pharyngeal muscle-specific gene expression.

Remarkably, the vast majority (91%, 356 genes out of 391, Table S12) of differentially expressed genes were not associated with differentially accessible elements (Figure 4A,B; Figure 4—figure supplement 2A,B). Specifically, out of 30 *de novo*-expressed pan-cardiac genes that were also differentially expressed upon FGF-MAPK perturbation at 18 hpf, only 8 (27%±8%, SE) were associated with one differentially accessible element following perturbation of FGF-MAPK signaling (Figure 4B). Similarly, out of 23 *de novo*-expressed pharyngeal muscle genes, which were also differentially expressed upon FGF-MAPK perturbation at 18 hpf, 11 were associated with one differentially accessible element following perturbation of FGF-MAPK signaling (Figure 4—figure supplement 3A). This suggested that most differential gene expression in early heart and pharyngeal muscle precursors arise from differential *cis*-regulatory activity of elements that are otherwise accessible throughout the cardiopharyngeal mesoderm. In keeping with this hypothesis, accessible regions associated with *de novo*-expressed pan-cardiac and pharyngeal muscle markers tended to open between 6 and 10 hpf, in a pattern consistent with FGF- and Foxf-dependent cardiopharyngeal-specific accessibility (Figure 4C; Figure 4—figure supplement 3B). These observations suggest that *cis*-regulatory elements controlling cell-type-specific *de novo* gene expression open in multipotent progenitors, prior to becoming active in fate-restricted precursors. Such decoupling between enhancer accessibility and activity has been observed in other developmental contexts, including early cardiogenesis in mammals (Paige et al., 2012; Wamstad et al., 2012).

As a proof of principle, we analyzed the *Lrp4/8* locus, which harbors two intronic elements (*KhC4.137* and *KhC4.144*) that opened upon TVC induction in an FGF- and Foxf-dependent manner, prior to *Lrp4/8* upregulation in cardiac progenitors (Wang et al., 2019), and were not differentially accessible at 18 hpf (Figure 4C,D). Of the two regions, only *KhC4.137* was sufficient to drive *GFP* expression in heart precursors indicating enhancer activity, and illustrating the decoupling between early and broad accessibility and late, cell-type-specific, activity (Figure 4E). Reporter gene expression and CRISPR/Cas9-mediated mutagenesis assays followed by FISH indicated that *KhC4.137* is both necessary and sufficient to activate gene expression in heart precursors (Figure 4E,F), showing that it acts as a *bona fide* enhancer, and demonstrating a specific case of decoupling between early and broad accessibility and late, cell-type-specific, activity.

To identify candidate regulators of late accessibility and/or activity, we parsed accessible elements associated with *de novo*-expressed heart and pharyngeal muscle markers into pre-accessible/primed or *de novo*-accessible elements and discovered sequence motifs enriched in each category (Figure 4—figure supplement 4A; Table S13). Putative binding sites for SMAD and homeodomain proteins such as Smad4 and Pitx respectively were enriched among pre-accessible elements associated with cardiac markers, and found in the primed elements regulating *Lrp4/8* upregulation (Figure 4—figure supplement 4A,B), suggesting a specific role in transcriptional activation, consistent with conserved roles for Pitx2 and BMP-SMAD signaling during heart development (Figure 4—figure supplement 4C,D; (Nowotschin et al., 2006; Schultheiss et al., 1997)). Motifs for known regulators of cardiac development, including Meis (Desjardins and Naya, 2016; Paige et al., 2012), were over-represented among *de novo*-accessible elements associated with cardiac markers, suggesting roles in establishing accessibility and/or regulating enhancer activity (Figure 4—figure supplement 4A). Notably, GATA motifs were enriched in primed accessible elements associated with cardiac markers, consistent with conserved roles for GATA factors as pioneer factors, and during cardiac development (Pikkarainen et al., 2004). Among motifs enriched in accessible elements associated with *de novo*-expressed pharyngeal muscle markers, the presence of ETS-, bHLH, and EBF-family motifs is consistent with established roles for FGF-MAPK, Hand-r, Mrf and Ebf in pharyngeal muscle specification (Razy-Krajka et al., 2018a, 2014; Stolfi et al., 2014, 2010; Wang et al., 2013). Notably, the enrichment of EBF motifs among *de novo*-accessible elements associated with *de novo*-expressed pharyngeal muscle markers is reminiscent of the ability of EBF-family factors to interact with nucleosome-bound cognate sites, suggestive of a pioneering activity in committed pharyngeal muscle precursors (Boller et al., 2016) (Buenrostro et al., 2013). In summary, this analysis identified distinct combinations of established and putative *trans*-acting factors differentially controlling the accessibility and/or activity of *cis*-regulatory elements that govern heart-vs. pharyngeal-muscle-specific gene expression (Figure 4H).

### Combinatorial *cis*-regulatory control of cardiopharyngeal determinants

The above analyses focused on one-to-one associations between accessible elements and neighboring genes to uncover candidate *trans*-acting inputs controlling gene expression through defined elements. However, most genes are associated with multiple accessible regions, especially developmental regulators (Figure 1—figure supplement 3D-F), presumably exposing diverse motifs for transcription factor binding. Moreover, distinct elements associated with the same neighboring gene often exhibited different accessibility dynamics. For instance, the loci of several *de novo*-expressed heart and pharyngeal muscle markers contained both primed-accessible and *de novo*-accessible elements (Figure 4C; Figure 4—figure supplement 3B; Figure 4—figure supplement 4A,B). This suggested that individual genes respond to a variety of regulatory inputs mediated through separate *cis*-regulatory elements.

To explore this possibility, we focused on *Tbx1/10* and *Ebf*, two established determinants of cardiopharyngeal fates (Razy-Krajka et al., 2018a, 2014; Stolfi et al., 2014, 2010; Tolkin and Christiaen, 2016; Wang et al., 2013). Both loci contained multiple accessible regions, including elements open already in the naive *Mesp*+ mesoderm (e.g. *Ebf*_*KhL24.35*/*36*), cardiopharyngeal-lineage-specific elements that open prior to gene activation but after induction of multipotent progenitors (e.g. *Ebf*_*KhL24.34*), and elements that open *de novo* in fate-restricted pharyngeal muscle precursors, where the gene is activated (e.g. *Ebf*_*KhL24.37*) (Figure 5A, B). Previous reporter gene expression assays identified the latter element, *Ebf*_*KhL24.37,* as a weak minimal enhancer with pharyngeal muscle-specific activity (Wang et al., 2013). CRISPR/Cas9-mediated mutagenesis assays followed by FISH indicated that each one of these elements is necessary for proper activation of *Ebf* in pharyngeal muscle progenitors (Figure 5C-E; Figure 5—figure supplement 1B,C). Consistently, we found that targeted deletions of individual accessible elements upstream of *Ebf* induced pharyngeal muscle precursor migration defects in 37%±5% (SE) of 28 hpf larvae when targeting *Ebf*_*KhL24.37* (n=101, Figure 5 - figure supplement 3). These observations indicate that each accessible *cis*-regulatory element upstream of *Ebf* is necessary for proper expression and subsequent pharyngeal muscle morphogenesis.

**Figure 5.**
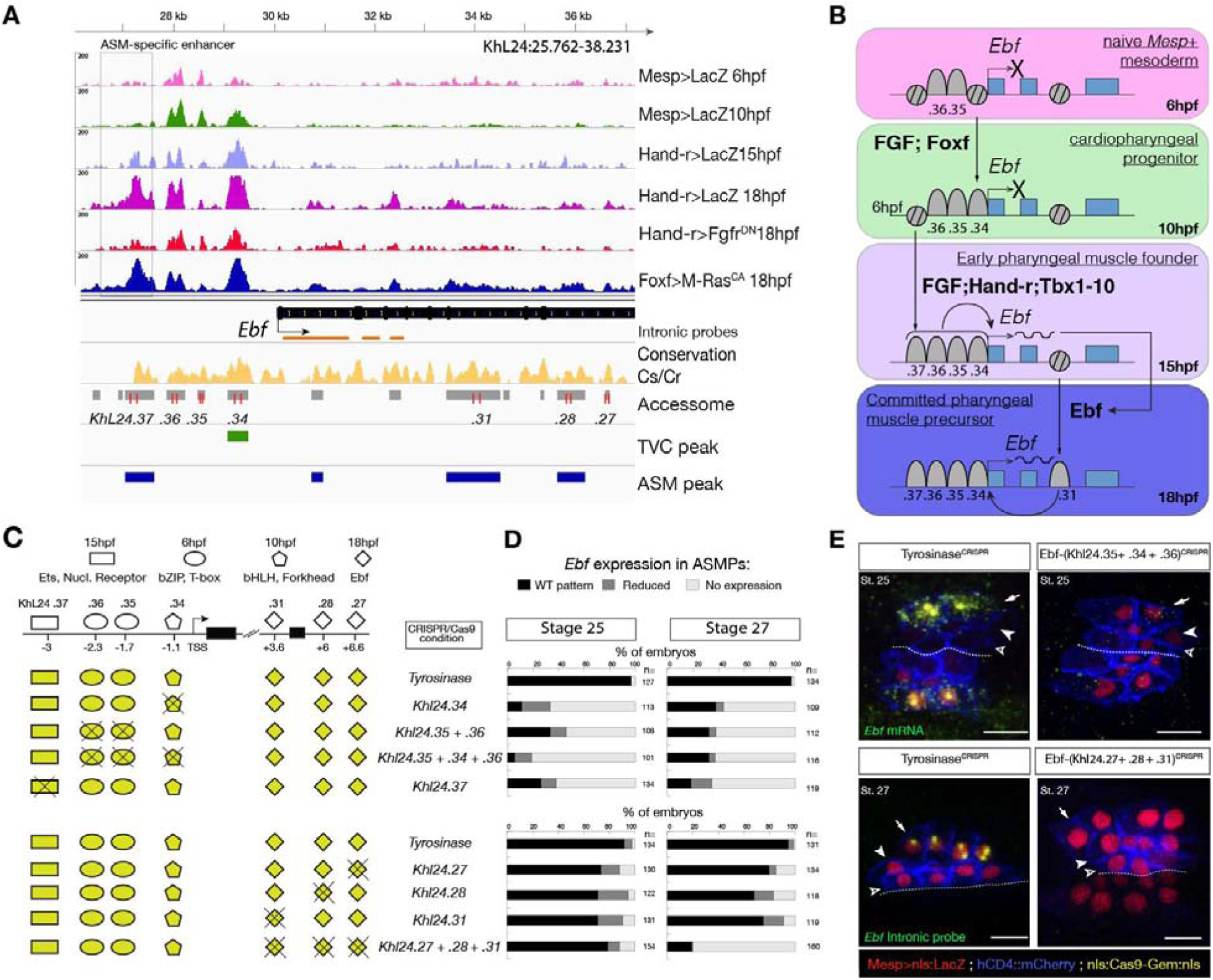
>Combinations of *cis*-regulatory elements with distinct chromatin accessibility profiles are required for *Ebf* transcription in pharyngeal-muscle precursors. (**A**) A 12 kb region of the scaffold L24 displaying expression profiles of RNA-seq and chromatin accessibility profiles of ATAC-seq (normalized tag count) in the *Ebf* locus. sgRNAs used to target ATAC-seq peaks are shown in red; intronic antisense riboprobes are shown in orange (**B**) Schematic representation showing sequential opening of *cis*-regulatory elements required for *Ebf* activation in pharyngeal muscle founder cells, and maintenance by auto-regulation in committed precursor. (**C**) Schematic representation of *Ebf cis*-regulatory elements targeted for CRISPR/Cas9-mediated deletions. Shapes represent binding sites located in the regulatory elements and differentially accessible over time. (**D**) Proportions of larva halves showing the indicated *Ebf* transcription patterns, in indicated experimental conditions; all the treatments were significant versus *Tyrosinase* (Fisher exact test, *p* < 0.001). (**E**) Endogenous expression of *Ebf* visualized by *in situ* (green) in Tyrosinase^CRISPR^ and upon CRISPR/Cas9-induced deletion of ATAC-seq peaks as indicated, at stage 25 (E) and 27 (F) based on (Hotta et al., 2007). For stage 25, an anti-sense riboprobe for the full length cDNA was used, whereas for stage 27 an intronic anti-sense riboprobe targeting the first three introns of *Ebf* transcript (orange lines) as previously used in (Wang et al. 2013). Nuclei of B7.5 lineage cells are labelled by *Mesp>nls::LacZ* and revealed with an anti beta-galactosidase antibody (red). Mesp driven hCD4::mCherry accumulates at the cell membrane as revealed by anti mCherry antibody (Blue). Scale bar = 10 µm.

Consistent with the established roles of Hand-r, Tbx1/10 and Ets-mediated FGF-MAPK signaling in activating *Ebf*, the primed cardiopharyngeal-specific element (*KhL24.34*) contained Fox and bHLH motifs, and the more distal *de novo*-accessible minimal enhancer (*KhL24.37*) also contained putative Ets and RORγ binding sites, whereas the constitutively accessible elements (*KhC24.35* and *.36*) contained primarily CREB and T-box binding sites (Figure 5C; Figure 5— figure supplement 1A). *Tbx1/10* showed a similar logic, whereby a constitutively accessible upstream element (*KhC7.909*) acts as an enhancer of cardiopharyngeal expression (Razy-Krajka et al., 2018a), and whose activity also requires a primed cardiopharyngeal-specific intronic element (*KhC7.914*) (Figure 5—figure supplement 2A-F). As a complement, and to more directly test the importance of enhancer accessibility, we targeted the intronic and distal elements in the *Tbx1/10* locus using dCas9::KRAB (Klann et al., 2017), which recruits deacetylases and presumably close chromatin (Sripathy et al., 2006); (Groner et al., 2010); (Schultz et al., 2002); (Reynolds et al., 2012; Thakore et al., 2015), and showed loss of *Tbx1/10* expression and function, as evaluated by expression of its target *Ebf* (Wang et al., 2013) (Figure 5 - figure supplement 4).

Of note, *Ebf* expression is maintained by auto-regulation (Razy-Krajka et al., 2018a), which requires separate intronic elements that harbor putative Ebf binding sites and open later (Figure 5B, Figure 5 - figure supplement 1A). Together with Ebf potent myogenic and anti-cardiogenic effects (Razy-Krajka et al., 2014; Stolfi et al., 2014, 2010; Tolkin and Christiaen, 2016), this auto-regulatory logic canalizes the pharyngeal muscle fate, stressing the importance of spatially and temporally accurate onset of expression to avoid ectopic ASM specification at the expense of cardiac identities, especially in the second heart lineage. These observations suggest that pharyngeal muscle fate specification relies on “combined enhancers”, characterized by a combination of *trans*-acting inputs mediated by distinct elements with variable dynamics of accessibility, to control the onset of *Ebf* expression in the cardiopharyngeal mesoderm (Figure 5F).

To test whether “combined enhancers” drive spatially and temporally accurate expression in pharyngeal muscle progenitors, we built a reporter containing multiple copies of the minimal, but weak, *Ebf* enhancer (*KhL24.37*) *(Wang et al., 2013)*. Two and three copies of the *KhL24.37* element (2x and 3x*KhL24.37*) significantly increased reporter gene expression in pharyngeal muscle precursors (43%±3% SE for 2x*KhL24.37* ; 58%±2% SE for 3x*KhL24.37*), compared to a single copy construct) (14%±3% SE for 1x *KhL24.37*) (Figure 6A-C), restoring reporter gene expression to levels similar to “full length” upstream element encompassing all combined enhancers, with endogenous genomic spacing (Ebf-full length −3348/−178) (Wang et al., 2013) (80%±1%, SE). Remarkably, unlike the “full length” combined enhancers, the 3x*KhL24.37* construct induced precocious reporter gene expression in the STVCs (89%±3%, SE) (n=95, Figure 6D-F) causing an ectopic GFP expression in the second heart lineage (13%±2%, SE) (n=218, Figure 6B,C). To test whether spacing between accessible elements could affect transcriptional output, we built a concatemer of KhL24.37, .36, .35, and .34 elements without endogenous spacer sequences. This construct increased the proportion of embryos with ASM cells expressing the reporter to 92%±2% (SE, n=130; Figure 6C), but it did not induce ectopic expression in the second heart lineage, supporting the notion that combined enhancers drive high but spatially and temporally accurate expression.

**Figure 6.**
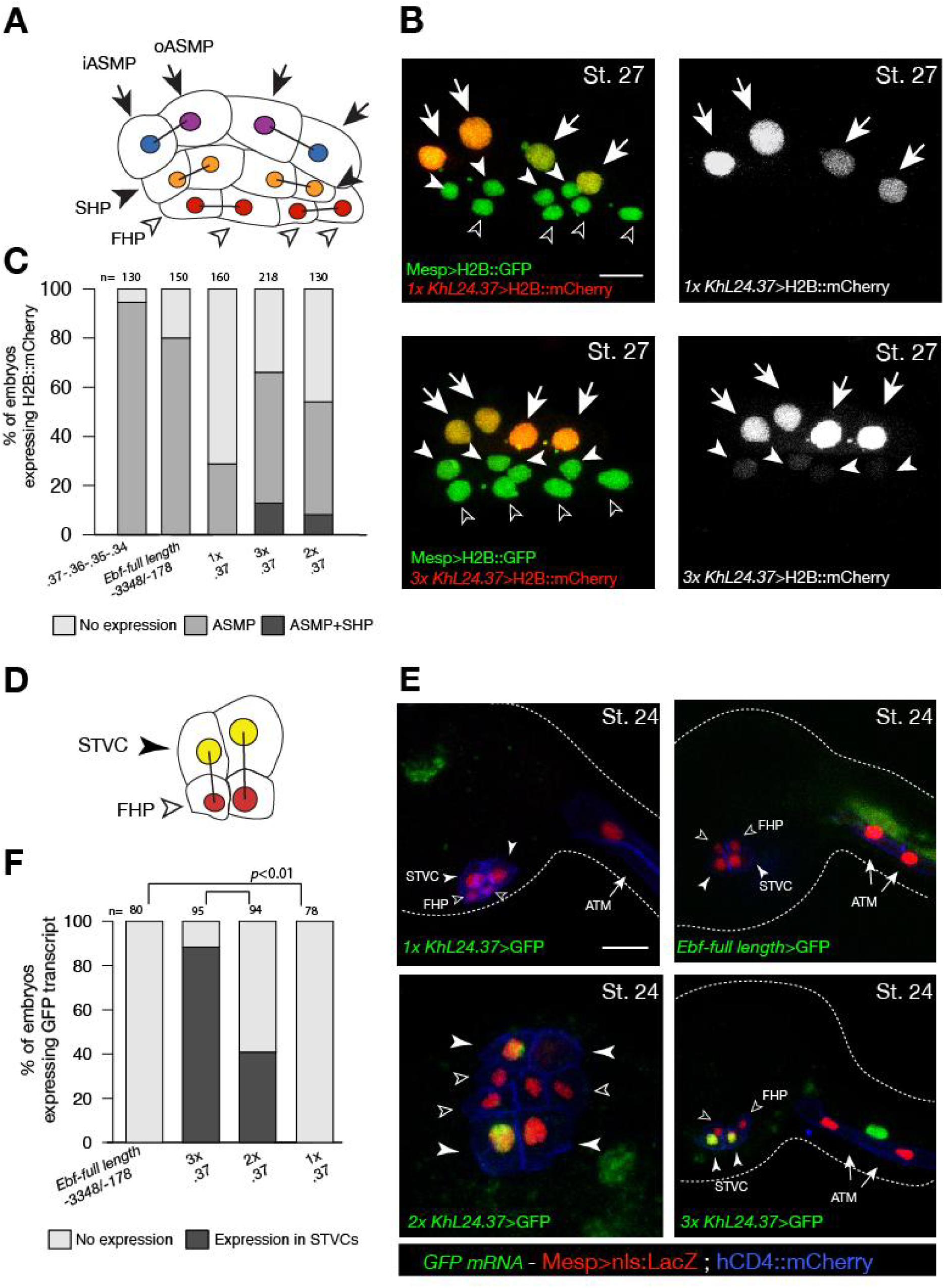
Multiple copies of a weak *Ebf* enhancer drive ectopic reporter gene expression. (**A**) Schematic representation of cardiopharyngeal lineage cells at Stage 27 (Hotta et al., 2007); First heart precursors (FHPs, red and open arrowheads), second heart precursors (SHPs, orange and arrows), inner ASM precursors and derivatives (iASMPs, violet and solid arrowhead), outer ASM precursors and derivatives (oASMPs, dark blue and solid arrowhead) (see (Razy-Krajka et al., 2014); black bars link sister cells. (**B**) Lineage tracing in individual larvae expressing single (1x) and multiple copy (3x) of *Ebf cis*-regulatory element, *KhL24.37*, driving H2B:mCherry (red) in cardiopharynegal progenitors at stage 27; B7.5 lineage is marked with *Mesp>H2B::GFP* (green). The single copy of *KhL24.37* element drives H2B::mCherry reporter expression specifically in the ASMPs (upper left panel, in white); three copies of *KhL24.37* (3xL24.37) drives expression in ASMPs and induces ectopic expression SHPs (lower left panel, in white). Experiment performed in biological replicate. Scale bar = 30 µm. (**C**) Proportions of embryos expressing H2B:mCherry in indicated cell-type progenitors by the indicated *cis*-regulaory elements. The “full length” upstream region encompassing all combined enhancers with endogenous spacing (Ebf-full length - 3348/−178)(Wang et al., 2013) as well as the concatemer of KhL24.37, .36, .35, and .34 elements, lacking endogenous spacer sequences were used as controls. Statistical analysis using a Fisher exact test showed all comparisons with either control to be significant (*p* < 0.01); “n” is the total number of individual halves scored per condition. (**D**) Schematic representation of cardiopharyngeal lineage cells at Stage 24 (Hotta et al., 2007); First heart precursors (FHPs, red and open arrowheads), secondary TVC (arrows). (**E**) Expression of GFP visualized by *in situ* hybridization on embryos at Stage 24 electroporated with single, multi copies and full-length of *Ebf cis*-regulatory element. Nuclei of B7.5 lineage cells are labelled by *Mesp>nls::LacZ* and revealed with an anti beta-galactosidase antibody (red). Mesp driven hCD4::mCherry accumulates at the cell membrane as revealed by anti mCherry antibody (Blue). Scale bar = 10 µm. (**F**) Proportions of embryos showing GFP driven by the indicated *Ebf cis*-regulatory element (Fisher exact test, *p* < 0.01).

We reasoned that high but precocious activation of *Ebf* should suffice to trigger the autoregulatory loop, and cause ectopic pharyngeal muscle specification in the cells that normally form the second heart lineage, as observed previously (Razy-Krajka et al., 2014; Stolfi et al., 2010). To directly test this possibility, we used different combinations of the *cis*-regulatory elements upstream of *Ebf* to drive expression of a functional *Ebf* cDNA, and assayed the effects endogenous *Ebf* expression and cardiopharyngeal fates (Figure 7). As expected, one copy of the *KhL24.37* element (1x*KhL24.37*) failed to cause ectopic activation of the endogenous locus, as evaluated using intronic probes, whereas using the *KhL24.37* trimer (*3xKhL24.37*) to drive expression of the *Ebf* cDNA sufficed to activate the endogenous locus ectopically in ~40% of the embryos, as shown by the presence of nascent transcripts in 4 out of 6 nuclei per side, instead of 2 (Figure 7A, B). This observation is consistent with our model that the upstream elements mediate high activating inputs for the onset of *Ebf* expression, whereas maintenance upon commitment to a pharyngeal muscle identity relies on autoregulation (Figure 5F; (Razy-Krajka et al., 2018a).

**Figure 7.**
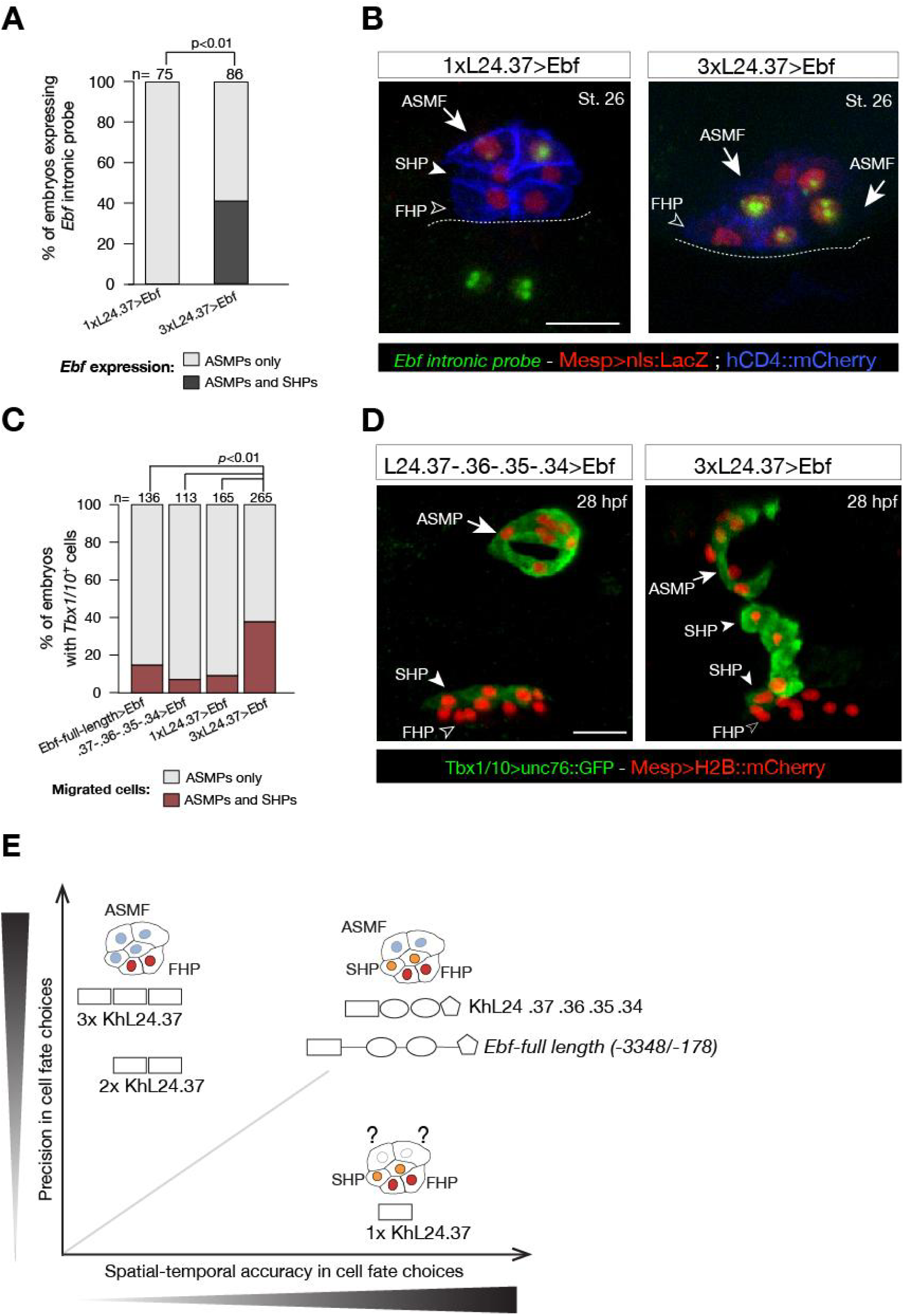
Multimer of one weak *Ebf* enhancer drives ectopic pharyngeal muscle fate specification. **(A)** Targeted expression of an *Ebf* cDNA by three copies of *KhL24.37* element induces expression of endogenous *Ebf* in four cells compared to the control where *Ebf* is detected only in the two ASMPs. Proportions of embryos expressing *Ebf* in ASMPs only or in ASMPs and SHPs in the indicated conditions. The single *KhL24.37 cis*-regulatory was used as control. **(B)** Expression of *Ebf* visualized by *in situ* hybridization on embryos at Stage 26 electroporated with single or three copies *KhL24.37* element driving *Ebf* cDNA. Intron-specific probes show nascent *Ebf* transcripts in ASMPs. Nuclei of B7.5 lineage cells are labelled by Mesp>nls::LacZ and revealed with an anti beta-galactosidase antibody (Red). *Mesp* driven hCD4::mCherry accumulates at the cell membrane as revealed by anti mCherry antibody (Blue). Scale bar = 10 µm. **(C)** Proportions of embryos showing GFP driven STVC-specific enhancer of *Tbx1/10* (Fisher exact test, *p* < 0.01). The “full length” upstream element encompassing all combined enhancers with endogenous spacing (Ebf-full length −3348/−178) as well as the “full length” without endogenous spacing (KhL24.37, .36, .35, .34) and the single copy *KhL24.37* element driving *Ebf* were used as a positive controls. **(D)** Example of an embryo at 28hpf showing GFP expression only in the ASM (solid arrowhead) and SHP (arrow) but not in the FHP (open arrowheads), where *Tbx1/10* enhancer is not active. Targeted expression of *Ebf* cDNA by three copies of *KhL24.37* element induces cells that normally form the second heart lineage to migrate alongside the pharyngeal muscle. Nuclei of B7.5 lineage cells are labelled by Mesp>H2B::mCherry. Scale bar = 15 µm. (**E**) Proposed role of combinatorial logics in fostering both precision and spatial and temporal accuracy of cell fate choices. The *Ebf*-full length, with or without endogenous spacers, fosters spatial / temporal accurate and precise cell fate choice, whereas the single copy of *KhL24.37* (1xL24.37) gives spatiotemporally accurate reporter expression but is likely insufficient to induce a precise ASM fate. Multiple copies of *KhL24.37 cis*-regulatory element rescue a high reporter activity that reflects precise pharyngeal fate at the expense of spatial and temporal accuracy. The shapes of the distinct *cis*-regulatory elements are as in Figure 5C. Statistical analysis using a Fisher exact test (*p*<0.01); “n” is the total number of individual halves scored per condition.

These results suggested that inaccurate activation and maintenance of *Ebf* expression would cause ectopic pharyngeal muscle specification at the expense of the second heart lineage. We tested this possibility by analyzing pharyngeal muscle morphogenesis in stage 30 larvae, which is characterized by collective migration away from heart progenitors and formation of a ring of atrial siphon muscle precursors (Figure 7). We used the STVC-specific *Tbx1/10* enhancer to visualize both the pharyngeal muscle precursors, which migrate and form a ring, and the second heart precursors, which remain associated with the *Tbx1/10*-negative first heart lineage (Figure 7D; (Razy-Krajka et al., 2018a). Remarkably, *Ebf* misexpression using three copies of the *KhL24.37* element induced cells that normally form the second heart lineage to migrate alongside the pharyngeal muscles in 38% of larvae at 28hpf (38%±3%, SE) (n=265, Figure 7 C-D). Importantly, neither one copy of *KhL24.37* (1x*KhL24.37*), nor the full combined enhancers, with or without endogenous spacers, sufficed to cause substantial fate transformation of *Tbx1/10*+ second heart lineage into migratory pharyngeal muscle precursors (Figure 7C-D). These results indicate that driving expression of an *Ebf* cDNA by multimerizing a weak *Ebf* enhancer sufficed to cause ectopic activation of the endogenous locus and transformation of second heart lineage cells into migratory pharyngeal muscle precursors.

Taken together, these results provide evidence for a decoupling between two essential aspects of transcriptional activation, whereby multiple copies of a single regulatory element enable robust gene expression, compatible with precise fate specification, albeit at the expense of spatial and temporal accuracy; whereas individual regulatory elements appear to integrate distinct *trans*-acting inputs controlling enhancer accessibility and/or activity, thus increasing the repertoire of regulatory inputs controlling developmental gene expression (Figure 7E). In other words, while multiple elements are required for proper activation of cell fate determinants, such as *Tbx1/10* and *Ebf*, in a manner reminiscent of super- and shadow enhancers (Lagha et al., 2012), we propose that combined enhancers foster spatially and temporally accurate cell fate decisions.

## DISCUSSION

We characterized the accessible genome of the tunicate *Ciona*, with a special focus on the cardiopharyngeal lineage that produces heart and pharyngeal muscles. As seen in other systems, less than 10% of the *Ciona* genome is accessible, and distributed across thousands of short regions, most of which are stably accessible across time and lineages, especially promoter regions. By contrast, developmentally regulated regions either closed upon induction of multipotent progenitors or opened specifically in the cardiopharyngeal lineage in response to FGF-MAPK signaling and Foxf activity. The latter elements were predominantly found in intergenic and intronic regions, and near cardiopharyngeal markers, consistent with their function as transcriptional enhancers. Similarly to other Forkhead factors (Zaret and Carroll, 2011), *Ciona* Foxf is required to open cardiopharyngeal elements for either immediate or later activation in multipotent or fate-restricted progenitors, respectively. Notably Foxf homologs play deeply conserved roles in visceral muscles specification (Jakobsen et al., 2007; Scimone et al., 2018; Zaffran et al., 2001), including during heart development in mammals (Hoffmann et al., 2014). GATA motifs are also over-represented among cardiopharyngeal-specific elements, consistent with a conserved role for GATA homologs in heart development (Holtzinger and Evans, 2007; Molkentin et al., 1997; Qian and Bodmer, 2009; Reiter et al., 1999; Sorrentino et al., 2005; Zhao et al., 2008). As combinations of FOX and GATA inputs play well-established roles in early endoderm specification (Cirillo et al., 2002), we speculate that cardiopharyngeal regulatory programs were built upon an ancestral endomesodermal chromatin landscape during Olfactores evolution.

The majority of cell-type-specific markers expressed *de novo* are associated with “primed accessible” elements, as observed in numerous systems including cardiac differentiation of embryonic stem cells (Paige et al., 2012; Wamstad et al., 2012), and consistent with the role of pioneer factors in establishing competence for subsequent activation (Zaret and Carroll, 2011). In the case of *Tbx1/10* and *Ebf*, spatially and temporally accurate activation is essential to permit the emergence of first and second cardiac, and pharyngeal muscle lineages (Razy-Krajka et al., 2018a, 2014; Wang et al., 2019, 2013). We found that several elements, exhibiting distinct accessibility dynamics, are required for proper activation of both *Tbx1/10* and *Ebf*. Specifically, a minimal distal enhancer proved necessary, but not sufficient for *Ebf* activation in newborn pharyngeal muscle precursors. By contrast, multiple copies of the same element sufficed to restore high transcriptional activity, but caused precocious activation in the *Tbx1/10*+ multipotent progenitors (aka STVCs, Figure 1A and Figure 6E), ectopic GFP expression in the second heart lineage (Figure 6), and eventually heart-to-pharyngeal muscle fate transformation when used to express *Ebf* itself (Figure 7). We propose that, whereas the activity of multiple elements with similar spatio-temporal transcriptional outputs permits precise and robust gene activation (Bentovim et al., 2017; Lagha et al., 2012), the modular organization of combined enhancers increases the repertoire of regulatory inputs, acting through both accessibility and activity, to control gene activation. Together with canalizing mechanisms, including positive auto-regulatory feedbacks, such multi-level combinatorial inputs achieve exquisite spatio-temporal control while permitting strong activation, thus ensuring both precise and accurate developmental fate choices (Figure 7E).

## MATERIALS AND METHODS

### Animals and electroporations

Gravid wild *Ciona intestinalis* type A, now called *Ciona robusta (Pennati et al., 2015)*, were obtained from M-REP (Carlsbad, CA, USA), and kept under constant light to avoid spawning. Gametes from several animals were collected separately for *in vitro* cross-fertilization followed by dechorionation and electroporation as previously described (Lionel Christiaen et al., 2009), and cultured in filtered artificial seawater (FASW) in agarose-coated plastic Petri dishes at 18°C. Different quantities of plasmids were electroporated depending on the constructs: the amount of fluorescent reporter DNA (*Mesp>nls::lacZ*, *Mesp>hCD4::mCherry*, *Mesp>tagRFP*, *MyoD905>eGFP* and *Hand-r>tagBFP*) and NLS::lacZ was typically 50 μg, but only 15 µg for *Mesp-1>H2B::mCherry*. For perturbation constructs (*Mesp>Fgfr^DN^, Mesp>Mek^S216D,S220E^, Foxf>M-Ras^CA^, Hand-r>Fgfr^DN^*), 70 µg were usually electroporated, except for *Mesp>nls::Cas9-Gem::nls* (30 µg) and pairs of *U6 >sgRNA* plasmids (25 µg each).

### Molecular cloning

Putative enhancers were amplified from *Ciona robusta* genome using primers containing specific sequence tails (Table S4) for subcloning into a vector upstream of a basal promoter from *Zfpm* (aka *Friend of GATA*/*Fog (Rothbächer et al., 2007)*) driving expression of green fluorescent protein (GFP) fused to cytoplasmic unc76 (Stolfi et al., 2010). The “full length” *Ebf* enhancer with non-endogenous spacing in the genome (KhL24.37, .36, .35, .34) was generated by synthesizing DNA fragment (Twist Bioscience) and subcloning into the full-length reporter plasmids.

*nls::dCas9-KRAB::nls* was derived from “pLV-dCas9-KRAB-PGK-HygR” (Klann et al., 2017) (Addgene plasmid: #83890) and inserted downstream of the promoter *Mesp* (Davidson, 2005).

### CRISPR/Cas9-mediated mutagenesis of ATAC-seq peaks

Two to four single guide RNAs (sgRNA) with Doench scores (http://crispor.tefor.net (Haeussler et al., 2016)) higher than 60 were designed to induce deletions in selected accessible elements using CRISPR/Cas9 in the B7.5 lineage as described (Gandhi et al., 2017). sgRNAs targeting non-overlapping sequences per gene are listed in Table S5. The efficiency of sgRNAs were evaluated using the peakshift method as described (Gandhi et al., 2017) (Figure 2—figure supplements 5B-C; Figure 2—figure supplement 6C). CRISPR/Cas9-mediated deletions were also evaluated by PCR-amplification directly from embryo lysates following electroporated with *Eef1a1>nls::Cas9-Gem::nls* (Figure 2—figure supplement 6C). sgRNAs were expressed using the *Ciona robusta U6* promoter (Stolfi et al., 2014) (Figure 2—figure supplement 5D; Figure 2— figure supplement 6C-D). For each peak, three or four guide RNAs were used in combination with 25 μg of each expression plasmid. 25 μg of *Mesp>nls::Cas9-Gem::nls* and *Mesp>nls::dCas9-KRAB::nls* plasmids were co-electroporated with guide RNA expression plasmids for B7.5 lineage-specific CRISPR/Cas9-mediated mutagenesis. Two guide RNAs were used to mutagenize Tyrosinase, which is not expressed in the cardiopharyngeal lineage and thus used to control the specificity of the CRISPR/Cas9 system (Wang et al., 2019).

### Fluorescent *In Situ* Hybridization-Immunohistochemistry (FISH-IHC) in *Ciona* embryos

FISH-IHC were performed as previously described (L. Christiaen et al., 2009; Razy-Krajka et al., 2018a). Embryos were harvested and fixed at desired developmental stages for 2 hours in 4% MEM-PFA and stored in 75% ethanol at −20°C. Antisense RNA probes were synthesized as described (Racioppi et al., 2014). For Ebf FISH-IHC, an anti-sense riboprobe targeting the full length cDNA was used at Stage 24 (Figure 5—figure supplement 1B) and 25 (Figure 5E, upper panel) where an anti-sense riboprobe targeting the first three intronic elements in the *Ebf* transcript as previously used in (Wang et al., 2013) was used for stage 27 (Figure 5E, lower panel; Figure 6E). The template for *Fbln* anti-sense riboprobe was PCR-amplified with the following oligos: Fbln_pb_fw (5’ TTGCGCTAAGTCATGACAGC 3’), Fbln_pb_rev (5’CATTTGCCGATTCAGCTATGT3’). *In vitro* antisense RNA synthesis was performed using T7 RNA Polymerase (Roche, Cat. No. 10881767001) and DIG RNA Labeling Mix (Roche, Cat. No. 11277073910). Anti-Digoxigenin-POD Fab fragment (Roche, IN) was first used to detect the hybridized probes, then the signal was revealed using Tyramide Signal Amplification (TSA) with Fluorescein TSA Plus Evaluation Kits (Perkin Elmer, MA). Anti–β-galactosidase monoclonal mouse antibody (Promega) was co-incubated with anti-mCherry polyclonal rabbit antibody (Bio Vision, Cat. No. 5993-100) for immunodetection of *Mesp>nls::lacZ* and *Mesp>hCD4::mCherry* products respectively. Goat anti-mouse secondary antibodies coupled with AlexaFluor-555 and AlexaFluor-633 were used to detect β galactosidase-bound mouse antibodies and mCherry-bound rabbit antibodies after the TSA reaction. FISH samples were mounted in ProLong® Gold Antifade Mountant (ThermoFisher Scientific, Waltham, MA. Catalog number P36930).

### Imaging

Images were acquired with an inverted Leica TCS SP8 X confocal microscope, using HC PL APO 63×/1.30 objective. Z-stacks were acquired with 1 μm z-steps. Maximum projections were processed with maximum projection tools from the LEICA software LAS-AF.

### Cell dissociation and FACS

Sample dissociation and FACS were performed as previously described (Christiaen et al., 2016; Wang et al., 2018). Embryos and larvae were harvested at 6, 10, 15, 18 and 20 hpf in 5 ml borosilicate glass tubes (Fisher Scientific, Waltham, MA. Cat.No. 14-961-26) and washed with 2 ml calcium- and magnesium-free artificial seawater (CMF-ASW: 449 mM NaCl, 33 mM Na_2_S O_4_, 9 mM KCl, 2.15 mM NaHCO_3_, 10 mM Tris-Cl pH 8.2, 2.5 mM EGTA). Embryos and larvae were dissociated in 2 ml 0.2 % trypsin (w/v, Sigma, T-4799) in CMF-ASW by pipetting with glass Pasteur pipettes. The dissociation was stopped by adding 2 ml filtered ice cold 0.05% BSA CMF-ASW. Dissociated cells were passed through 40 μm cell-strainer and collected in 5 ml polystyrene round-bottom tube (Corning Life Sciences, Oneonta, New York. REF 352235). Cells were collected by centrifugation at 800 g for 3 min at 4°C, followed by two washes with ice cold 0.05 % BSA CMF-ASW. Cell suspensions were filtered again through a 40 μm cell-strainer and stored on ice. Cell suspensions were used for sorting within 1 hour. B7.5 lineage cells were labeled by *Mesp>tagRFP* reporter. The B-line mesenchyme cells were counter-selected using *MyoD905>eGFP* as described (Christiaen et al., 2016; Wang et al., 2018). The TVC-specific *Hand-r>tagBFP* reporter was used in a 3-color FACS scheme for positive co-selection of TVC-derived cells (Wang et al., 2018), in order to minimize the effects of mosaicism. Dissociated cells were loaded in a BD FACS AriaTM cell sorter. 488 nm laser, FITC filter was used for EGFP; 407 nm laser, 561 nm laser, DsRed filter was used for tagRFP and Pacific BlueTM filter was used for tagBFP. The nozzle size was 100 μm. eGFP^+^ mesenchyme cells were collected for downstream ATAC-seq analysis, whereas tagRFP^+^, tagBFP^+^ and eGFP^−^cardiopharyngeal lineage cells were collected for both ATAC- and RNA-seq analyses.

### Preparation and sequencing of ATAC-seq library

ATAC-seq was performed with minor modifications to the published protocol (Buenrostro et al., 2015). 4000 cells obtained by FACS were centrifuged at 800 x *g* for 4 min at 4 °C for each sample. Cells were resuspended in 5 µL of cold lysis buffer (10 mM Tris-HCl [pH 7.4], 10 nM NaCl, 3 nM MgCl_2_, 0,1% v/v Igepal CA-360 [Sigma-Aldrich] and incubated on ice for 5 min. After centrifugation of the cells at 500 x *g* for 10 min at 4 °C, the supernatant was discarded. The tagmentation reaction was performed at 37 °C for 30 min using Nextera Sample Preparation kit (Illumina) with the addition of a tagmentation-stop step by the addition of EDTA to a final concentration of 50 nM and incubation at 50°C for 30 minutes (Hockman et al., 2018). After tagmentation, transposed DNA fragments were amplified using the following PCR condition: 1 cycle of 72°C for 5 min and 98°C for 30 s, followed by 12 to 14 cycles of 98°C for 10 s, 63°C for 30 s and 72°C for 1 min. Amplified libraries were purified using PCR purification MinElute kit (Qiagen), and the quality of of the purified library was assessed by using 2100 Bioanalyzer (Agilent Technologies) and a High Sensitivity DNA analysis Kit (Agilent Technologies) to confirm a period pattern in the size of the amplified DNA. After size selection by using Agencourt Ampure XP beads (Beckman Coulter, Cat#A633880) with a bead-to-sample ratio of 1.8:1, the libraries were sequenced as paired-end 50 bp by using the HiSeq 2500 platform (Illumina) according to the manufacturer’s instructions. An input control was also generated by using 10 ng of *C. robusta* genomic DNA.

### ATAC-seq data analysis

#### Alignment of ATAC-seq reads

Raw reads were preprocessed by FastQC (version 0.11.2, http://www.bioinformatics.babraham.ac.uk/projects/fastqc) and adaptors were trimmed using Trim Galore (version 0.4.4, http://www.bioinformatics.babraham.ac.uk/projects/trim_galore) before being aligned to *Ciona robusta* genome (joined scaffold (KH), http://ghost.zool.kyoto-u.ac.jp/datas/JoinedScaffold.zip) using Bowtie2 (version 2.3.2, (Langmead and Salzberg, 2012) with the parameters “--no-discordant -p 20 -X 1000”. Reads with mapping quality score > 30 were kept for downstream analysis using SAMtools (version 1.2, (Li et al., 2009). Mitochondrial reads were removed using NGSutils 0.5.9 (Breese and Liu, 2013). At least three replicates for each sample were merged for peak calling by MACS2 (version 2.7.9) (Zhang et al., 2008) (-- nomodel --nolambda --gsize 1.2e8). We defined the accessome by concatenating all narrow peaks from MACS2 using bedtools merge (2.26.0) (Quinlan and Hall, 2010). Peaks were filtered for length > 50 bp, to obtain a final number of 56,090 peaks.

#### Accessome annotation

We annotated each peak to all transcripts from the *C. robusta* genome (http://ghost.zool.kyoto-u.ac.jp/datas/KH.KHGene.2013.gff3.zip) within 10 kb using GenomicRanges (version 1.28.6, (Lawrence et al., 2013). We also identified whether each peak overlapped with the genomic feature categories “TSS, 5’ UTR, intron, exon, 3’ UTR, TTS, or intergenic region” (Figure 1— figure supplement 1C). We defined a window of 107 bp upstream and downstream of the start of the 5’ UTR of any transcript as the TSS. We defined a window of 1 kb upstream the TSS to be the promoter, which we further divided into two 500 bp windows. We defined the TTS to be a window 200 bp upstream and downstream from the end of the 3’ UTR of any transcript. We defined regions not covered by any feature to be intergenic.

#### Differential accessibility analysis

We evaluated the significant change in chromatin accessibility between different samples using DEseq2 (Love et al., 2014) with the parameters (method = “LRT”, alpha = 0.05, cooksCutoff = FALSE) for differential accessibility analysis. Libraries with fewer than 500,000 reads were removed. We performed a likelihood ratio test for four models:

1. the 10 hpf model, tested for difference in accessibility between any of the conditions at 10 hpf (*Mesp>Fgfr^DN^, Mesp>Mek^S216D,S220E^, U6 >Foxf.1, U6>Foxf.4*,listed in Table S5) and controls (*Mesp>nls::LacZ* and *U6>Ngn.p1, U6>Ngn.p2* used for Foxf^CRISPR^, listed in Table S5).
2. the 15 - 20 hpf model, tested for difference in accessibility between *Hand-r>Fgfr^DN^* and control *Hand-r>LacZ* at any of the later time points (15, 18 or 20 hpf), *Foxf>M-Ras^CA^* and *Foxf>nls::LacZ* for the 18 hpf samples, versus a null model where accessibility is only dependent on time.
3. the time model, tested for any change in accessibility between control B7.5 samples (*Mesp>nls::LacZ* and *Hand-r>nls::LacZ*) at 6, 10, 15, 18 and 20 hpf versus a null model dependent where accessibility is not changing.
4. the control vs. mesenchyme model, tested for any difference in accessibility between B7.5 control (*Mesp>nls::LacZ* and *Hand-r>nls::LacZ*) and mesenchymal (*MyoD905>eGFP*) samples versus a null model dependent at 10, 15, 18 and 20 hpf.

#### Cell-type-specific accessibility

We defined cell-type specific accessibility based on the peak sets:

- Elements opened by FGF-MAPK at 10 hpf: the union of peaks closed in *Mesp>Fgfr* vs. *Mesp>nls::LacZ* (log_2_FC < −0.5 and FDR < 0.05) and peaks open in *Mesp>Mek^S216D,S220E^* vs. *Mesp>Fgfr^DN^* (log_2_FC > 0.5 and FDR < 0.05) in model 1.
- Elements closed in FGF-MAPK at 10 hpf: the union of peaks closed in *Mesp>Mek^S216D,S220E^* vs. *Mesp>nls::LacZ* and peaks closed in *Mesp>Mek^S216D,S220E^* vs. *Mesp>Fgfr^DN^* (log_2_FC < −0.5 and FDR < 0.05) in model 1.
- Elements opened in FGF-MAPK at 18 hpf: the union of peaks closed in *Hand-r>Fgfr^DN^* vs. *Hand-r>nls::LacZ* (log_2_FC < −0.5 and FDR < 0.05 in the 15-20hpf model), open in *Foxf>M-Ras^CA^* vs. *Foxf>nls::LacZ*, and open in *Foxf>M-Ras^CA^* vs. *Hand-r>Fgfr^DN^* (log_2_FC > 0.5 and FDR < 0.05) in model 2.
- Elements closed in FGF-MAPK at 18hpf: the union of peaks open in *Hand-r>Fgfr^DN^* vs. *Hand-r>nls::LacZ* (log_2_FC > 0.5 and FDR < 0.05 in the 15-20 hpf model), closed in *Foxf>M-Ras^CA^* vs. *Foxf>nls::LacZ*, and closed in *Foxf>M-Ras^CA^* vs. *Hand-r>Fgfr^DN^* (log_2_FC < −0.5 and FDR < 0.05) in model 2.
- *De novo* elements: peaks closed in control 10 vs. 18 hpf (log_2_FC < −0.5 and FDR < 0.05 in model 3 that are not opened by FGF-MAPK at 10 hpf.
- Primed elements: peaks opened in FGF-MAPK at 10 hpf that are not closed in *Mesp>LacZ* 10 vs. 18 hpf.

We tested for enrichment of peaks overlapping with each category of genomic feature in each of these peak sets (Figure 2—figure supplement 2C).

#### Definition of gene sets

We used cardiopharyngeal lineage cell-type-specific gene sets (cardiopharyngeal markers), including primed and *de novo*-expressed Cardiac and ASM/pharyngeal muscle as well as secondary TVC (STVC), first heart precursor (FHP), second heart precursor (SHP) markers defined by scRNA-seq (Wang et al., 2019). Anterior tail muscles (ATM) marker genes were derived from publicly available expression data (Brozovic et al., 2018) (Table S6). To these we added the gene sets either activated or inhibited by MAPK at 10 or 18 hpf as defined by microarray (Christiaen et al., 2008) and Bulk RNA-seq analysis (Wang et al., 2019).

#### Gene Set Enrichment Analysis (GSEA)

We performed GSEA on elements associated to these gene sets using fgsea 1.2.1 (Sergushichev, 2016) with parameters: minSize=5, maxSize=Inf, nperm=10000. Because the gene sets used for GSEA need not be exclusive, we can use ATAC-seq peaks as “genes” and define a gene set as all peaks annotated to genes in the set. We measured enrichment for peaks associated to the gene sets in open peaks (indicated by a positive enrichment score, ES) or closed peaks (indicated by a negative ES) in each comparison.

#### Motif analyses of ATAC-seq

Motifs were selected and associated to *Ciona* TFs as described in the Supplementary Text. We compiled the *C. robusta* genome (http://ghost.zool.kyoto-u.ac.jp/datas/JoinedScaffold.zip) as a BSGenome (version 1.44.2) package using “forgeBSgenomeDataPkg”. We searched for all motifs present in accessible elements using motifmatchr (version 0.99.8) and TFBSTools (version 1.14.2, (Tan and Lenhard, 2016). Background nucleotide frequencies were computed for the whole accessome. Matches were returned if they passed a cutoff of *p* < 5*10^−5^. This cutoff was used to set a score threshold using the standard method of comparing the probability of observing the sequence given the PWM versus the probability of observing the sequence given the background nucleotide sequence of the accessible elements (Staden, 1989). We tested for motif enrichment in a peak set versus the accessome using a one-tailed hypergeometric test. The odds ratio represents the probability that an outcome (a peak containing the indicated motif) will occur if the peak is contained in the indicated peak set, compared to the probability of the outcome occurring in an element randomly selected from the accessome. For Figure 4—figure supplement 4S18A, the test indicates the probability that an element will contain a motif given when the element is accessible. To ensure that the elements’ accessibility is not reflecting expression of primed genes, elements were removed if they were also associated to a gene expressed earlier in the cardiopharyngeal lineage. Elements associated to *de novo* ASM genes were removed if they were also associated to a cardiac or FGF-MAPK inhibited at 18hpf gene. Elements associated to *de novo* cardiac genes were removed if they were also associated to an ASM or FGF-MAPK activated at 18hpf gene.

Accessibility of motif sites in differentially accessible (FDR < 0.05) peaks in the 10 hpf model was calculated using chromVAR (version 0.99.3, (Schep et al., 2017). Peaks were resized to 200 bp. Deviations were computed between all 10 hpf conditions as well as the 6 hpf controls. Replicates were grouped by both conditions and time. We tested for significantly differentially accessible motifs (FDR < 0.01) using the “differentialDeviations” function from chromVAR. For the enrichment of motif families, primed or *de novo* accessible elements associated to *de novo* cardiac expressed genes were considered cardiac accessible if they had log_2_(FC) < 0 in *Foxf>M-Ras^CA^* vs. *Foxf>nls::LacZ* or log_2_(FC) > 0 in *Hand-r>Fgfr^DN^* vs. *Hand-r>nls::LacZ*. Primed or *de novo* elements associated to *de novo* pharyngeal muscle (ASM) genes were considered ASM accessible if they had log_2_(FC) > 0 in *Foxf>M-Ras^CA^* vs. *Foxf>nls::LacZ* or log_2_(FC) < 0 in *Hand-r>Fgfr^DN^* vs. *Hand-r>nls::LacZ*. Motifs and TF families were taken from CIS-BP. To reduce redundancy, only SELEX and HocoMoco motifs were selected. The EBF motif was added from HOMER (Heinz et al., 2010). A one-tailed hypergeometric test was performed for enrichment of peaks containing any motif in a family in a peak set versus the accessome. The peak sets tested were primed cardiac accessible peaks associated to *de novo* cardiac expressed genes, *de novo* cardiac accessible peaks associated to *de novo* cardiac expressed genes, primed ASM accessible peaks associated to *de novo* ASM expressed genes, and *de novo* ASM accessible peaks associated to *de novo* ASM expressed genes. We also tested for enrichment of individual motifs from CIS-BP in these peak sets as described above.

### Bulk RNA-seq Library Preparation, Sequencing and Analyses

We used bulk RNA-seq performed following defined perturbations of FGF-MAPK signaling and FACS (Wang et al., 2019). To profile transcriptomes of FACS-purified cells from Foxf^CRISPR^ and control samples, 1000 cells were directly sorted in 100 μl lysis buffer from the RNAqueous®-Micro Total RNA Isolation Kit (Ambion). For each condition, samples were obtained in two biological replicates. The total RNA extraction was performed following the manufacturer’s instruction. The quality and quantity of total RNA was checked using Agilent RNA Screen Tape (Agilent) using 4200 TapeStation system. RNA samples with RNA Integrity Number (RIN) > 9 were kept for downstream cDNA synthesis. 250-2000 pg of total RNA was loaded as a template for cDNA synthesis using the SMART-Seq v4 Ultra Low Input RNA Kit (Clontech) with template switching technology. RNA-Seq Libraries were prepared and barcoded using Ovation® Ultralow System V2 1–16 (NuGen). Up to 6 barcoded samples were pooled in one lane of the flow cell and sequenced by Illumina NextSeq 500 (MidOutput run). Paired-end 75 bp length reads were obtained from all the bulk RNA-seq libraries. Bulk RNA-seq libraries were aligned using STAR 2.5.3a (Zhang et al., 2018) with the parameters “--runThreadN 20 -- bamRemoveDuplicatesType UniqueIdentical”. Counts were obtained using htseq-count (0.6.1p1) (Anders et al., 2015). Differential expression was calculated using edgeR for the 10hpf conditions and DESeq2 for the conditions at 18hpf.

#### Cell-type-specific expression gene sets

We tested for differential expression in the bulk RNA-seq data for the pairwise comparisons Foxf^CRISPR^ vs control^CRISPR^, *Foxf>M-Ras^CA^* vs. control (*Foxf>nls::LacZ*), *Hand-r>Fgfr^DN^* vs. control (*Hand-r>nls::LacZ*), and *Foxf>M-Ras^CA^* vs. *Hand-r>Fgfr^DN^* using DESeq2 1.16.1(Love et al., 2014). We defined genes downregulated in Foxf^CRISPR^ vs. control^CRISPR^ (log_2_FC < −0.75 and FDR < 0.05) as Foxf targets. We defined “MAPK-inhibited genes at 18 hpf” as the intersection of genes downregulated in *Foxf>M-Ras^CA^* vs. control and genes downregulated in *Foxf>M-Ras^CA^* vs. *Hand-r>Fgfr^DN^* (log_2_FC < −1 and FDR < 0.05) as. We defined the intersect of genes downregulated in *Hand-r>Fgfr^DN^* vs. control (log_2_FC < −1 and FDR < 0.05) and genes upregulated in *Foxf>M-Ras^CA^* vs. *Hand-r>Fgfr^DN^* (log_2_FC > 1 and FDR < 0.05) as “MAPK activated genes” (Figure 4—figure supplement 1A-D). For Figure 4B and Figure 4—figure supplement 3A, we defined the union of differentially expressed genes in either *Foxf>M-Ras^CA^* vs. control or *Hand-r>Fgfr^DN^* vs. control as “differentially expressed upon FGF-MAPK perturbation at 18 hpf.”

We integrated microarray data to obtain gene sets for MAPK perturbation at 10 hpf (Christiaen et al., 2008). We defined genes downregulated in *Mesp>Fgfr^DN^* vs. control (log_2_FC < −1 and p < 0.05) as “MAPK activated genes at 10 hpf.” We defined genes upregulated in *Mesp>Fgfr^DN^* vs. control (*Mesp>nls::LacZ*) (log_2_FC >1 and p < 0.05) as “MAPK inhibited genes at 10 hpf.” From this same data set we obtained genes downregulated in control 6hpf vs. 10hpf (log_2_FC < −1 and p < 0.05) and upregulated in LacZ 6hpf vs. 10hpf (log_2_FC > 1 and p < 0.05).

### Statistics

We used R (version 3.6.1) (Rizzo, 2016) to perform all statistical analysis. An alpha level of 0.05 was adapted for statistical significance throughout the analyses. We corrected for multiple hypothesis testing using false discovery rate (FDR) where indicated. All other *p*-values are unadjusted. For statistics presented in FISH panels, a Fisher’s exact test was run for each condition vs. the control. Embryos were classified as wild type, reduced expression, or no expression. A *p*-value less than 0.01 was considered significant (Figure 2F; Figure 2—figure supplement 4; Figure 3F; Figure 4G; Figure 5—figure supplement 2B,E). The binomial tests in Figure 1—figure supplements 1C, 2D, and 3G assume accessibility to not be feature-specific and dependent only on the fraction of the genome in base pairs covered by a feature. If promoter sites comprise 10% of the genome, we expect 10% of accessible elements will overlap with promoter sites by chance. For Figure 3—figure supplement 7A and B, we performed a binomial test on the joint probability of each intersection against the null hypothesis that the probability of each set is independent. The null probability was calculated as the product of the probabilities of all constituent sets of an intersection. This is similar to the method used to calculate the deviation score in the original UpSet plot (Lex et al., 2014)

### Software

All computations were performed on an x86_64-centos-linux-gnu platform. In addition to software specified elsewhere in the Methods section, we created a SQLite database using RSQLite 2.1.1 (Müller et al., 2018), ComplexHeatmap (2.0.0) (Gu et al., 2016), circlize (0.4.6) (Gu et al., 2014), latticeExtra (0.6-28) (Sarkar and Andrews, 2016) and Integrative Genomics Viewer (Robinson et al., 2011).

### Code availability

All analyses were done with publicly available software. All scripts for data analysis are available at https://github.com/ChristiaenLab/ATAC-seq

### Data availability

All sequencing data were deposited on GEO with accession GSE126691.

## ACKNOWLEDGEMENTS

We are grateful to Karen Lam, Sana Badri, Emily Miraldi and Richard Bonneau for help processing ATAC-seq data in the early phase of this study. We thank Dayanne M. Castro for help and discussion on ATAC-seq data analysis.. We thank Emily Huang for TVC-specific enhancers validation, Tina Jiang, Elizabeth Zelid and Carly Vaccaro for help with reporter assays.We thank Alberto Stolfi for assistance on cloning and constructive input and discussions. We are grateful to Pui Leng Ip for support with the FACS, Wei Wang for sharing single cell and bulk RNA-seq data as well as Florian Razy-Krajka for sharing *Mek^S216D,S220E^* prior to publication.. We thank Esteban Mazzoni and members of the Christiaen lab for discussions, in particular Yelena Bernadskaya and Keaton Schuster for comments on the manuscript and figures. We thank Tatjana Sauka-Spengler for advice with ATAC-seq protocol. C.R. has been supported by a long-term fellowship ALTF 1608-2014 from EMBO. This work was funded by awards R01 HL108643 from NIH/NHLBI, R01 HD096770 from NIH/NICHD, and 15CVD01 from the Leducq Foundation to L.C.

## AUTHORS CONTRIBUTIONS

C.R. and L.C. obtained funding. C.R., K.W. and L.C. designed the experiments and analyses, C.R. performed the experiments, C.R. and K.W. performed the computational analyses, C.R., K.W. and L.C. wrote the paper.

## SUPPLEMENTARY TEXT

### General characterization of the cardiopharyngeal accessome

We prepared 39 ATAC-seq libraries from FACS-purified B7.5/cardiopharyngeal lineage and B-line mesenchyme cells, whole *Ciona* embryos, and genomic DNA controls (Table S1). To generate a reference peak set (aka “accessome”), we first identified ATAC-seq peaks in all 39 libraries using MACS2 (Zhang et al., 2008). To correct for nonspecific sequencing biases, we subtracted sequenced gDNA from these libraries as described (Toenhake et al., 2018). Then, we combined and merged peaks to generate an atlas of 56,090 unique and non-overlapping accessible regions (Figure 1—figure supplement 1F). The accessome was more similar in GC content to the coding regions of the genome (43%) than the non-coding regions (35%), as observed in other studies (Li et al., 2007; Madgwick et al., 2018) (Figure 1—figure supplement 1E). Moreover, we did not observe any difference in GC content by genomic features (Figure 1—figure supplement 1D). Peaks in the accessome had a mean length of 190 bp and covered 9.25% of the *C. robusta* genome (Figure 1—figure supplement 1B,F,G).

Approximately half of *C. robusta* protein-coding genes exhibit spliced-leader (SL) *trans*-splicing editing, which replaces the original 5’ end sequence of pre-mRNAs (the outron) by a short non-coding RNA (Ganot et al., 2004; Hastings, 2005; Satou et al., 2006; Vandenberghe et al., 2001). For these genes, the actual transcription start site (TSS) is thus not represented by the 5’ end of the mature mRNA. To circumvent this difficulty and identify TSSs, we used a TSS-seq dataset that produced high-density regions of TSSs, or TSS clusters (TSCs) (Yokomori et al., 2016) for *Ciona* larval stage as guidelines for core promoter length. We used the mean TSC length plus two standard deviations to define windows of 107 bp upstream and downstream of the start of each transcript as the putative TSS. We extended these regions 1 kb upstream to obtain putative promoters, and accessible regions overlapping with these putative promoters were considered as ATAC-seq promoter peaks. Out of 20,266 promoter proximal regions, 9.7% (1,973 peaks) overlapped with TSCs obtained from TSS-seq (Yokomori et al., 2016) (Figure 1—figure supplement 2C). Moreover, we categorized the ATAC-seq promoter peaks in relation to the distance to the TSS. We found that 13,559 peaks overlapped with a window of 500 bp upstream of the TSS (ATAC-seq promoter 0.5 kb), whereas 12,250 peaks overlapped with a window of 500 bp to 1 kb upstream of the TSS (ATAC-seq promoter 1 kb).

As a complement, we characterized putative differentially expressed (DE) promoters by bulk RNA-seq reads aligned to the putative promoters, as defined above. 3,326 of these DE promoters (16.4%) overlapped with ATAC-seq promoter peaks (Figure 1—figure supplement 2C). A two-tailed binomial test showed that our ATAC-seq peaks associated to promoters (0.5 and 1kb from TSS), TSS sites an 5’UTR significantly overlapped with promoter regions identified by TSS-seq, while intergenic regions were significantly depleted in TSCs (*p* < 0.001; Figure 1—figure supplement 2D). Promoter regions were enriched for known core promoter motifs, such as TATA-box, Inr, BRE, DCE and DPE (Haberle and Stark, 2018) (Figure 1—figure supplement 2E). Enriched motifs were consistent between promoter peaks, TSS-seq as well as DE promoters. Enriched motifs also correlated between non-coding regions within and adjacent to the putative TSS, as well as 5’ and 3’ UTR (Figure 1—figure supplement 2F).

We associated an ATAC-seq peak to an annotated gene if it overlapped a window encompassing 10 kb upstream of the transcription start site (TSS), the gene body with introns, and 10 kb downstream of the transcription termination site (TTS) (Figure 1—figure supplement 3A) (Brozovic et al., 2018). Using this tentative proximity-based peak-to-gene assignments, we associated genes with a median of 11 peaks, whereas peaks associated with a median of 3 genes (Figure 1—figure supplement 3B,C). As expected, longer genes tended to associate with more peaks (Figure 1—figure supplement 3D). In the specific case of regulatory genes encoding transcription factors (TF) and signaling molecules, we observed more associated peaks (median 18) of similar median length to other peaks (162 and 154 bp for transcription factors -TFs- and signaling molecules, respectively; Figure 1—figure supplement 2E-G), reminiscent of ‘shadow enhancer’(Hong et al., 2008) and ‘super enhancer’(Whyte et al., 2013). A binomial test showed that the accessome contains more peaks associated to TFs and signaling molecules than would be expected by chance, considering the fraction of the genome covered by these genes (6,081 TF peaks, 2,473 signaling molecule peaks, median = 18, *P*<0.001, Figure 1—figure supplement 3G).

We next examined the distribution of the peaks within the accessome at specific elements in the *Ciona* genome, including core promoter (500 bp and 1 kb upstream of TSS), coding exons, introns, 3’ untranslated region (UTR) and 5’ UTR, transcription termination sites (TTS) and proximal peaks (Figure 1—figure supplement 3A). Approximately 13% of accessible regions localize to TSS, 11.9% within 500 bp of TSS and 9.6% between 0.5 and 1kb. These proportions are significantly higher than expected by chance considering the fraction of the genome covered by each class of elements (p<0.01, two-tailed binomial test; Figure 1—figure supplement 2B), thus indicating that promoter proximal regions tend to be accessible as observed in other systems (Mayran et al., 2018). The genomic regions most represented were introns (covering 32.8% of the accessome), intergenic regions (covering 20.8% of the accessome), and 500 bp upstream of the TSS (covering 20.2% of the accessome). However, whereas 20.9% of regions 500 bp upstream of the TSS in the genome are covered by the accessome, only 11.7% of intergenic regions in the genome and 6.4% of intronic regions in the genome are covered by the accessome. The remaining peaks are positioned more distally and are roughly evenly divided between intronic (20.1%) and intergenic peaks (21%) where *cis*-regulatory elements, such as transcriptional enhancers, are typically encountered(Long et al., 2016) (Figure 1—figure supplement 1E). Of note, introns were significantly under-represented in our lineage-specific accessome, possibly revealing that intronic regulatory elements tend to become accessible in a more context-dependent manner, compared to promoter proximal regions.

Together, these results indicate the reproducibility of high-resolution chromatin accessibility obtained from FACSorted cardiopharyngeal- and mesenchymal lineage cells.

### Ciona robusta motif inference

We obtained inferred *C. intestinalis* (former name of *C. robusta*) transcription factor (TF) binding motifs from CIS-BP (Weirauch et al., 2014). The TF binding motifs were inferred based on similarity of DNA-binding domain sequences between *C. robusta* TFs and those of TFs with known binding motifs. A strong correlation has been observed between conserved DNA-binding domains and bound motifs (Weirauch et al., 2014).

### Motif selection

To the inferred motifs we added SELEX-seq motifs for *C. robusta* from (Nitta et al., 2019). We then searched the *C. robusta* genome for orthologs to known motifs from the HOMER database. All motifs with orthologous *C. robusta* TFs were added to the set of candidate TF motifs. This set of candidate TF motifs consisted of 1745 motifs associated to 294 *C. robusta* TFs. Of these, 1228 motifs could be unambiguously associated to one of 237 TFs. To obtain motifs for the remaining 57 TFs, to minimize redundancy, we remove from consideration all associations to TFs with an assigned motif, then assign all unambiguous motifs to a TF. This iteration continues until no more unambiguous motifs can be assigned. Motifs for the remaining TFs are assigned by repeating the process for motifs associated to only two TFs, then three TFs, until each TF has an assigned motif. Our final set of motifs had 1487 members, with the median TF associated to two motifs.

For Figure 1—figure supplement 2E, we used previously published eukaryotic core promoter motifs (Haberle and Stark, 2018). For Figure 1—figure supplement 2F, correlations were performed on the inferred C. robusta DNA-binding motifs.

### Gene Set Enrichment Analysis (GSEA)

For GSEA of the ATAC-seq data, peaks were ranked by log_2_(FC) for a specific pairwise comparison, and all peaks annotated to a gene in a given gene set (as described above) were considered members of that gene set. The normalized enrichment score (NES) is a measure of whether members of the gene set are enriched at the top or bottom of a ranked list (Figure 1D). NES is calculated by keeping a running tally of whether or not the peak at each position of the list is a member of the gene set and comparing the maximum value to that of a null distribution calculated from a random permutation of peaks. For example, In Figure 1D, a positive NES indicates that peaks associated to a gene set are more open at 6 hpf than at 10 hpf, while a negative NES indicates that peaks associated to a gene set are more accessible at 10 hpf than at 6 hpf.

### chromVAR algorithm

The accessibility of a motif is calculated by summing the fragment counts for all accessible elements containing the motif. An expected accessibility of the motif is calculated from the total counts for a motif from all replicates normalized to the library size of each replicate. For each biological replicate, the raw accessibility deviation of each motif is defined as the difference of the observed counts and the expected counts divided by the expected counts. A normally distributed background accessibility deviation for each motif is calculated by iteratively sampling accessible elements from the same replicate. The deviation Z-score for each motif is given by the difference of the raw accessibility deviation and the mean of the background accessibility deviation divided by the standard deviation of the background accessibility deviation. For a more thorough description of the algorithm underlying chromVAR, see (Schep et al., 2017).

## SUPPLEMENTARY FIGURES

**Figure 1—figure supplement 1.**
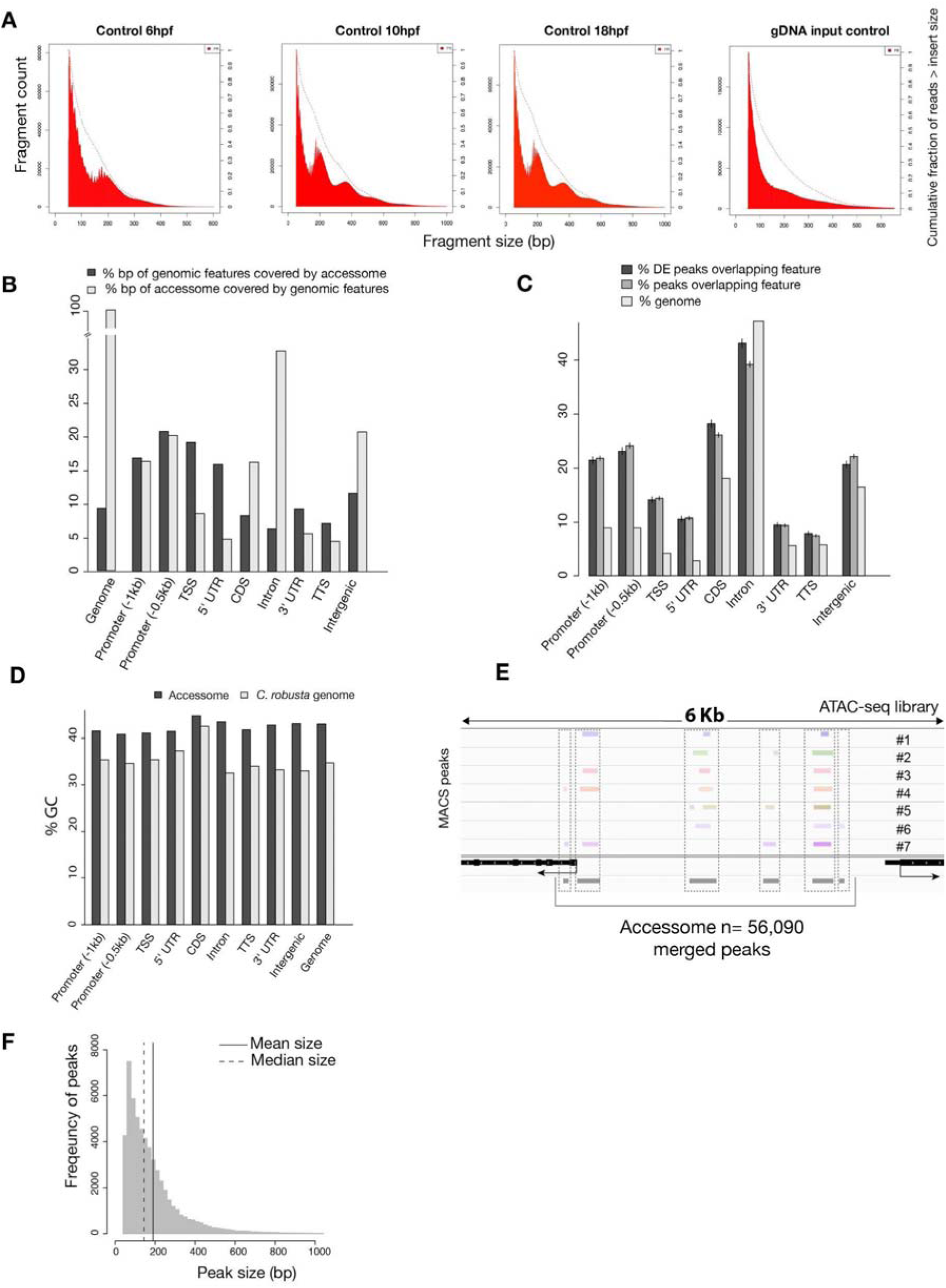
General characterization of the accessome. (**A**) Fragment size for B7.5 control samples at developmental time indicated (first three panels from the left) showing consistent approximately 147 bp periodicity, likely corresponding to nucleosome-protected fragments. This periodicity is not present in the purified genomic DNA (gDNA) input control (far right panel). Dotted lines show the cumulative fraction of reads larger than a given size. (**B**) Proportion of genomic features covered by the accessome and proportion of the accessome covered by genomic features (in bp). (**C**) Two-tailed binomial test for enrichment of accessible elements overlapping a genomic feature. Accessible elements associated to any DE gene from any scRNA-seq or bulk RNA-seq experiment were considered DE peaks. Bars show the predicted (based on bp coverage of the genome by a feature) and observed probabilities that an accessible element will overlap a genomic feature. Error bars show the 99% confidence interval. (**D**) Comparison of GC content of genomic features and accessible regions overlapping these features. (**E**) A 6 kb region displaying MACS2-called peaks in seven ATAC-seq libraries (upper panel), gene model (black) and accessome (gray). (**F**) Accessible element size distribution.

**Figure 1—figure supplement 2.**
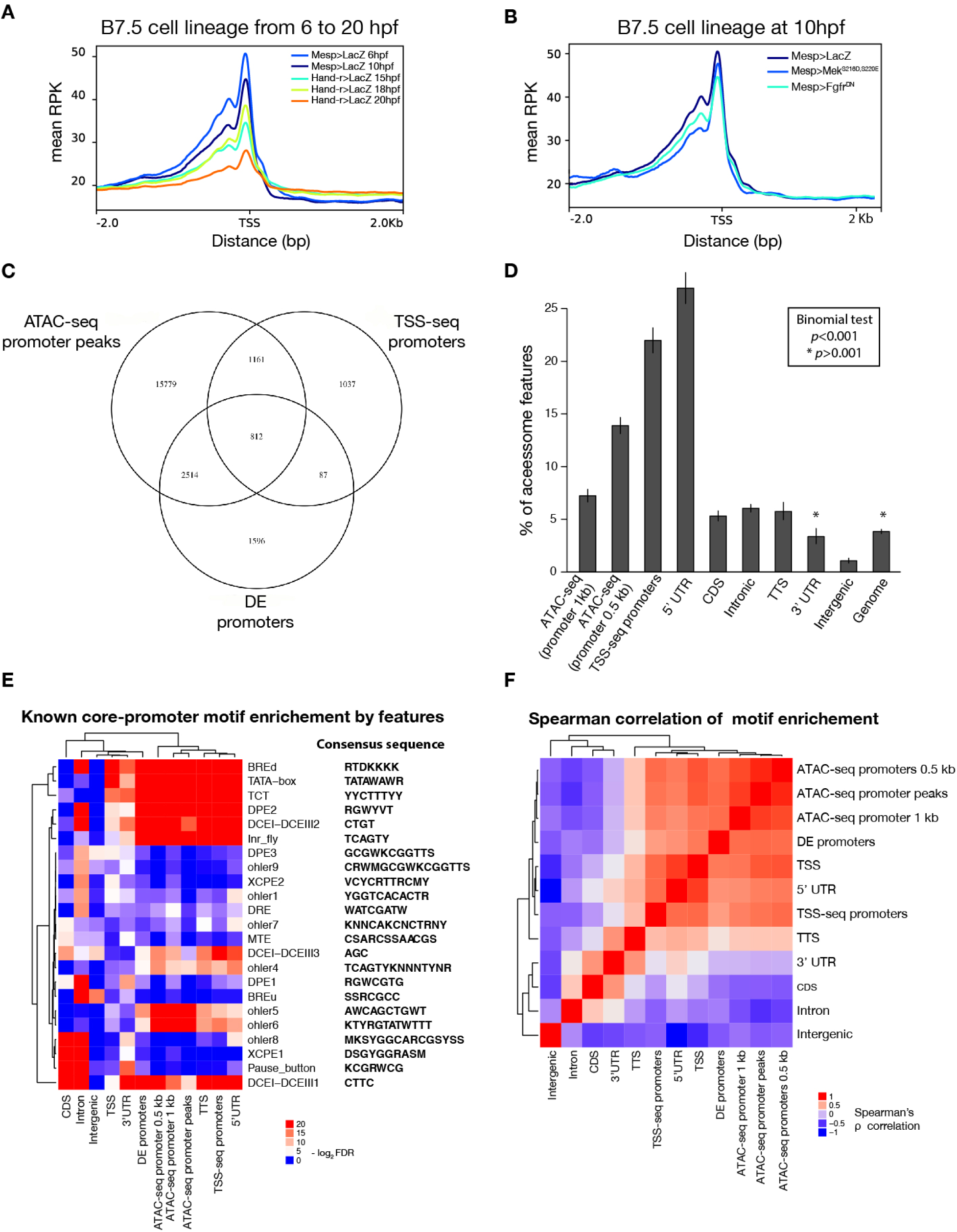
Characterization of promoter regions. (**A**) Read density within 2 kb upstream or downstream of TSS shows global closing of TSS over time. (**B**) Accessibility of the TSS is maintained in response to Fgfr^DN^ perturbation at 10 hpf. (**C**) Intersections of overlapping promoter elements. “ATAC-seq promoter peaks” are accessible elements overlapping the putative promoter (−1,107 to +107 bp from the TSS). “DE promoters” are differentially expressed putative promoters in any bulk RNA-seq condition (see Supplemental Methods). (**D**) Two-tailed binomial test for enrichment of accessible elements overlapping TSS-seq sites by genomic feature. Bars show the observed probability that an accessible element overlapping a genomic feature will also overlap a TSS-seq element. Error bars show the 99% confidence interval. The expected probability was calculated from the percent of the accessome overlapping TSS-seq elements (3.8%). (**E**) One-tailed hypergeometric test for enrichment of promoter motifs in genomic features. Eukaryotic core promoter motifs were taken from (Haberle and Stark, 2018). (**F**) Spearman correlation of inferred motif enrichment (see Supplementary Text). Motifs in each class of element were ranked based on a one-tailed hypergeometric test.

**Figure 1—figure supplement 3.**
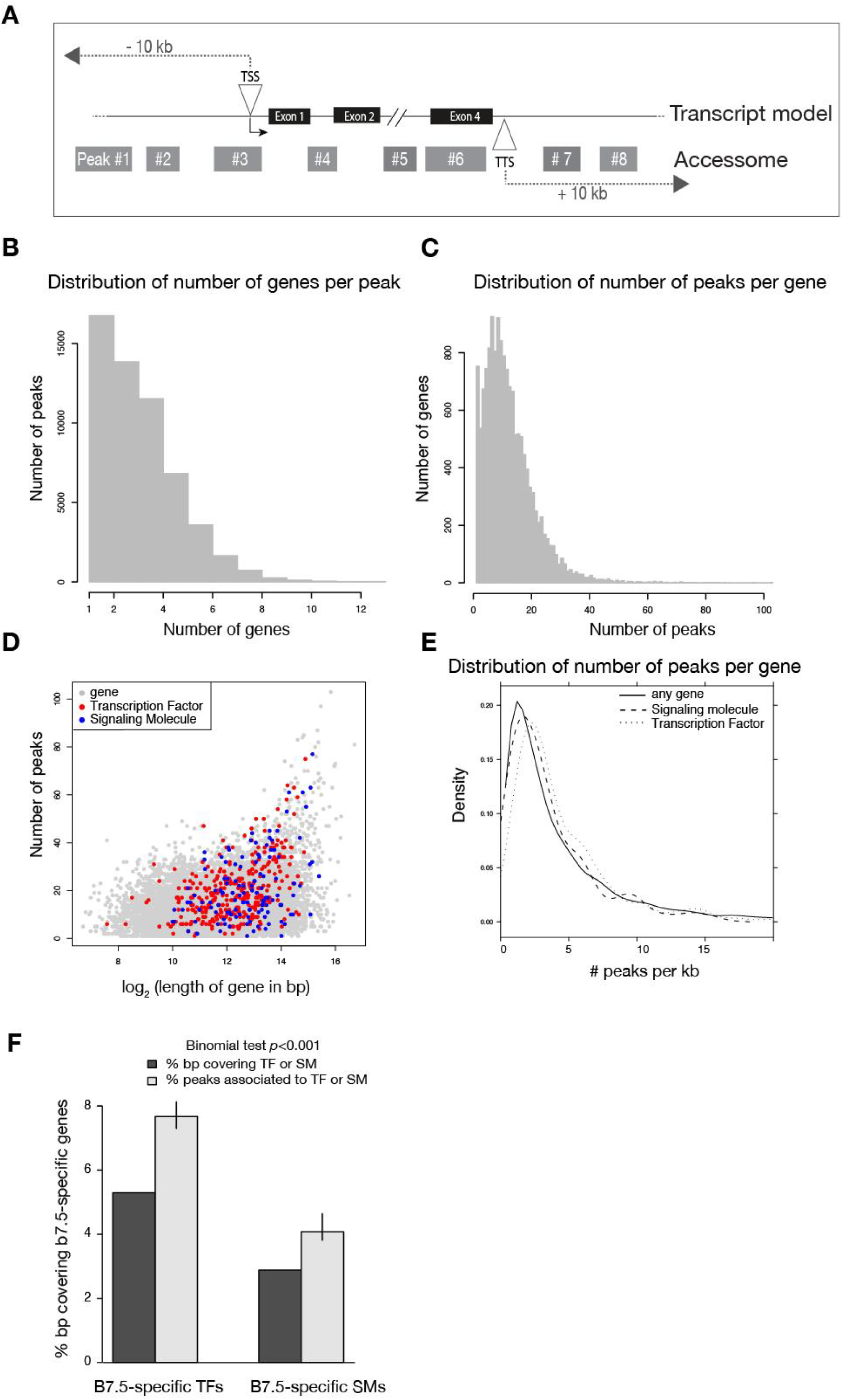
Annotation of the accessome. (**A**) Example diagram of accessible elements surrounding a gene locus. (**B**) Distribution of number of genes associated to each peak. (**C**) Distribution of number of peaks associated to each gene. (**D**) Gene size vs. number of associated peaks. Red and blue dots show all transcription factor (TF) and signaling molecule (SM) genes, respectively. (**E**) Density of peaks per kb of each gene. (**F**) Two-tailed binomial test for enrichment of accessible elements associated to TF or signaling molecule genes. Predicted probability of an element associating to a TF or signaling molecule is given by the proportion of bps in all gene bodies ±10kb covering TFs or signaling molecule gene bodies ±10kb. Error bars show the 99% confidence interval of the binomial test. Only genes that were differentially expressed in any scRNA-seq or bulkRNA-seq comparison (Wang et al., 2019) were considered.

**Figure 2—figure supplement 1.**
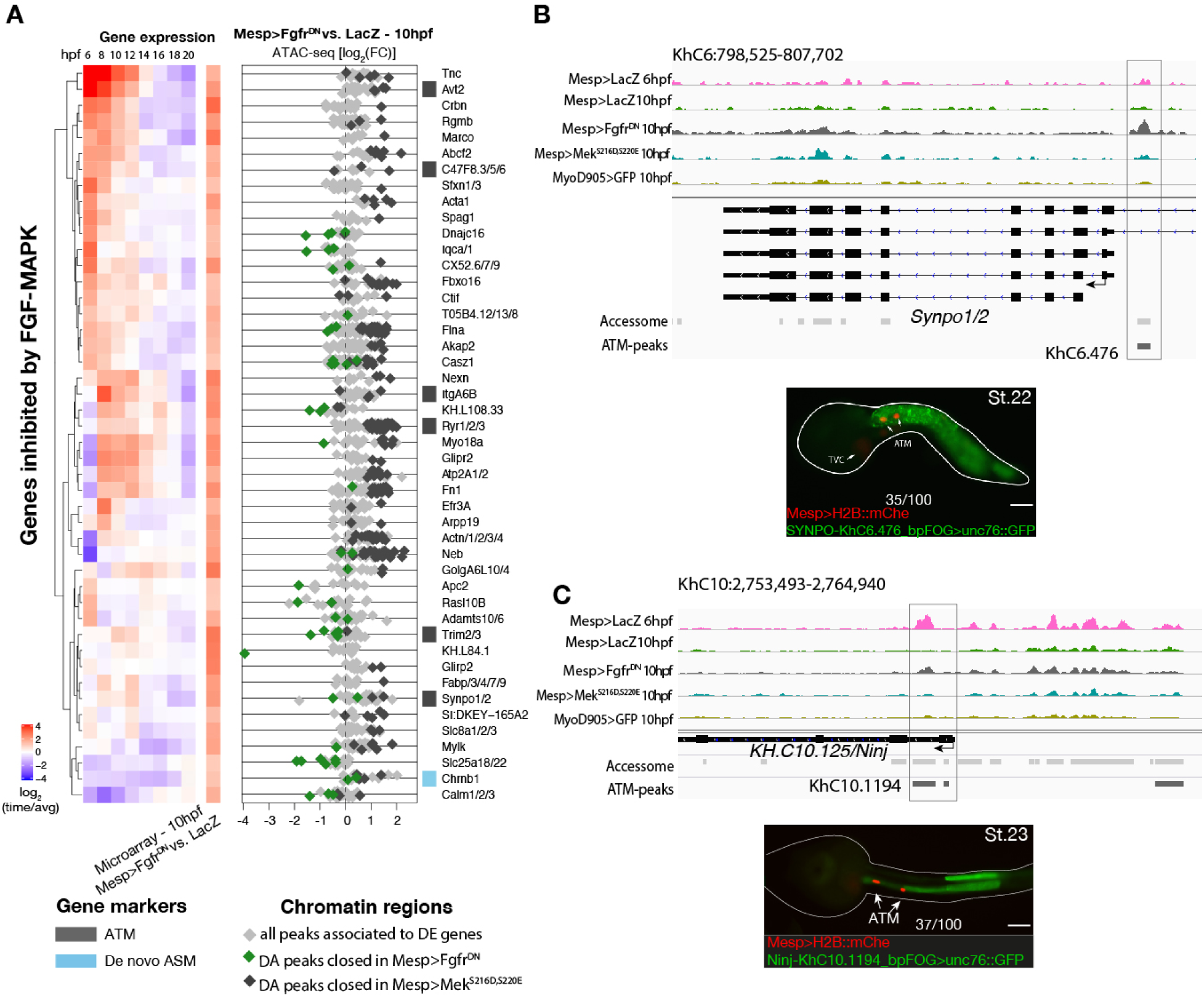
Inhibition of FGF signaling (Mesp>Fgfr^DN^ -10hpf) induces opening of ATM-specific elements. (**A**) Relation between expression and accessibility for DE genes in the top 99.25% quantile of fold change between expression in Fgfr^DN^ and control at 10hpf (log_2_FC > 1.31). (**B, C**) A 10 kb region on chromosome 6 and 10 including *Synpo1/2* (B) and *KH.C10.125* (C) displaying chromatin accessibility profiles from ATAC-seq (normalized by total sequencing depth). The newly identified enhancers are in boxed regions. ATAC-seq peaks were tested *in vivo* by reporter gene assay (see Table S4). Reporter gene assay in embryos at stage 23 (Hotta et al., 2007). GFP expression was detected specifically in ATM cells and not in TVCs. Nuclei of B7.5 lineage cells are labelled by *Mesp>H2B::mCherry* and *Mesp>hCD4:mCherry* (red). Numbers indicate observed/total of half-embryos scored. Scale bar= 30 μm.

**Figure 2—figure supplement 2.**
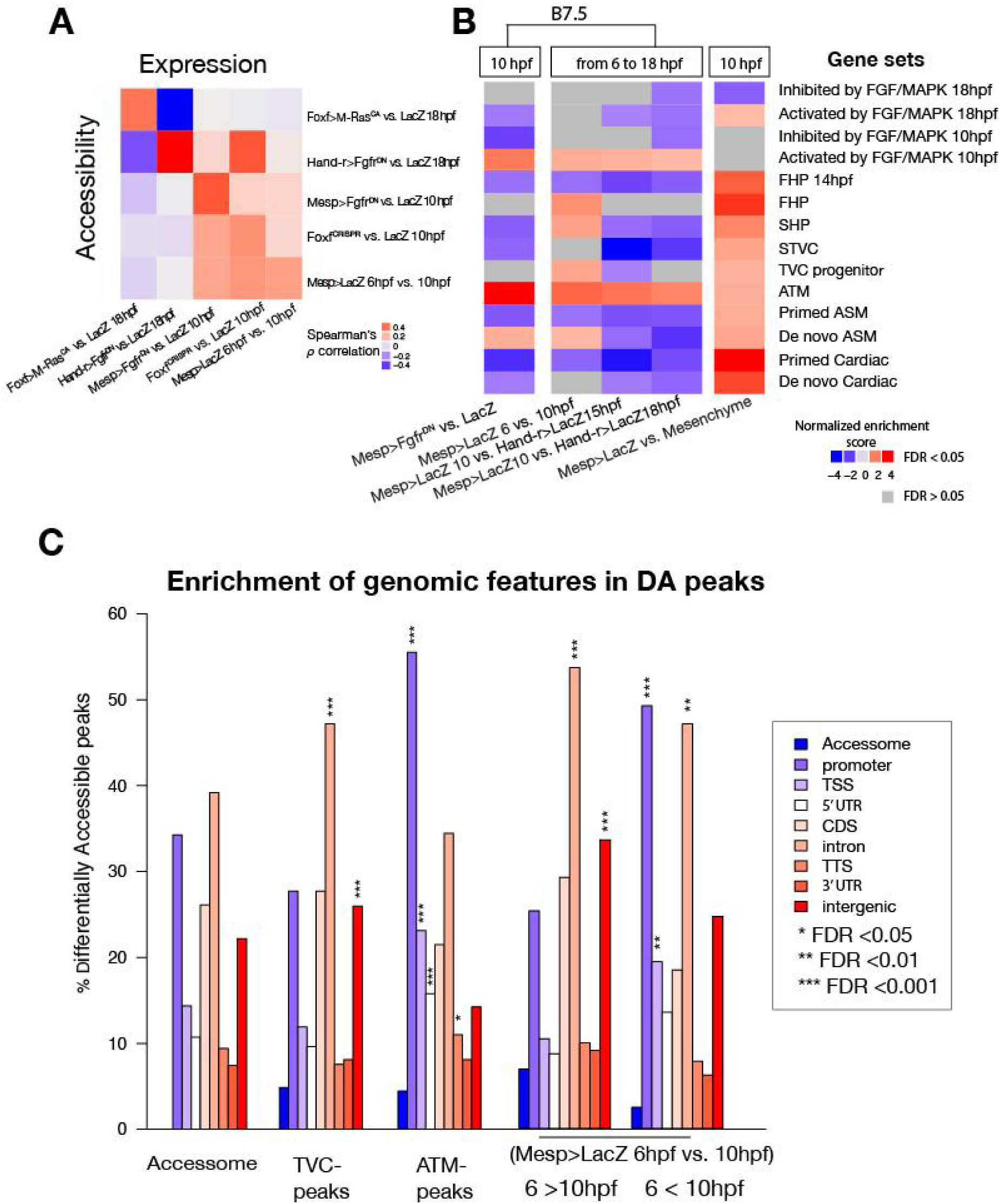
General characterization of differential accessibility. (**A**) Spearman correlation of expression of bulk RNA-seq or microarray (for B7.5 6 vs. 10 hpf and Mesp>Fgfr^DN^ vs. Mesp>LacZ - 10 hpf) pairwise comparisons with ATAC-seq pairwise comparisons. Correlation was calculated based on log_2_(FC) of differentially expressed genes associated to differentially accessible peaks for each comparison. Differentially expressed genes in Mesp>Fgfr^DN^ vs Mesp>LacZ at 10hpf derived from (Christiaen et al., 2008), Hand-r>Fgfr^DN^ vs Hand-r>LacZ and Foxf>M-Ras^CA^ vs Foxf>LacZ at 18hpf from (Wang et al., 2019), Foxf^CRISPR^ vs Control^CRISPR^ at 10hpf from the present study. (**B**) GSEA for ATAC-seq pairwise comparisons. A negative normalized enrichment score (NES) indicates elements annotated to that gene set are less accessible in the comparison. A positive NES indicates elements annotated to that gene set are more accessible in the comparison. (**C**) One-tailed hypergeometric test for enrichment of accessible elements overlapping a genomic feature. The leftmost column shows the expected distribution of accessible elements from the accessome.

**Figure 2—figure supplement 3.**
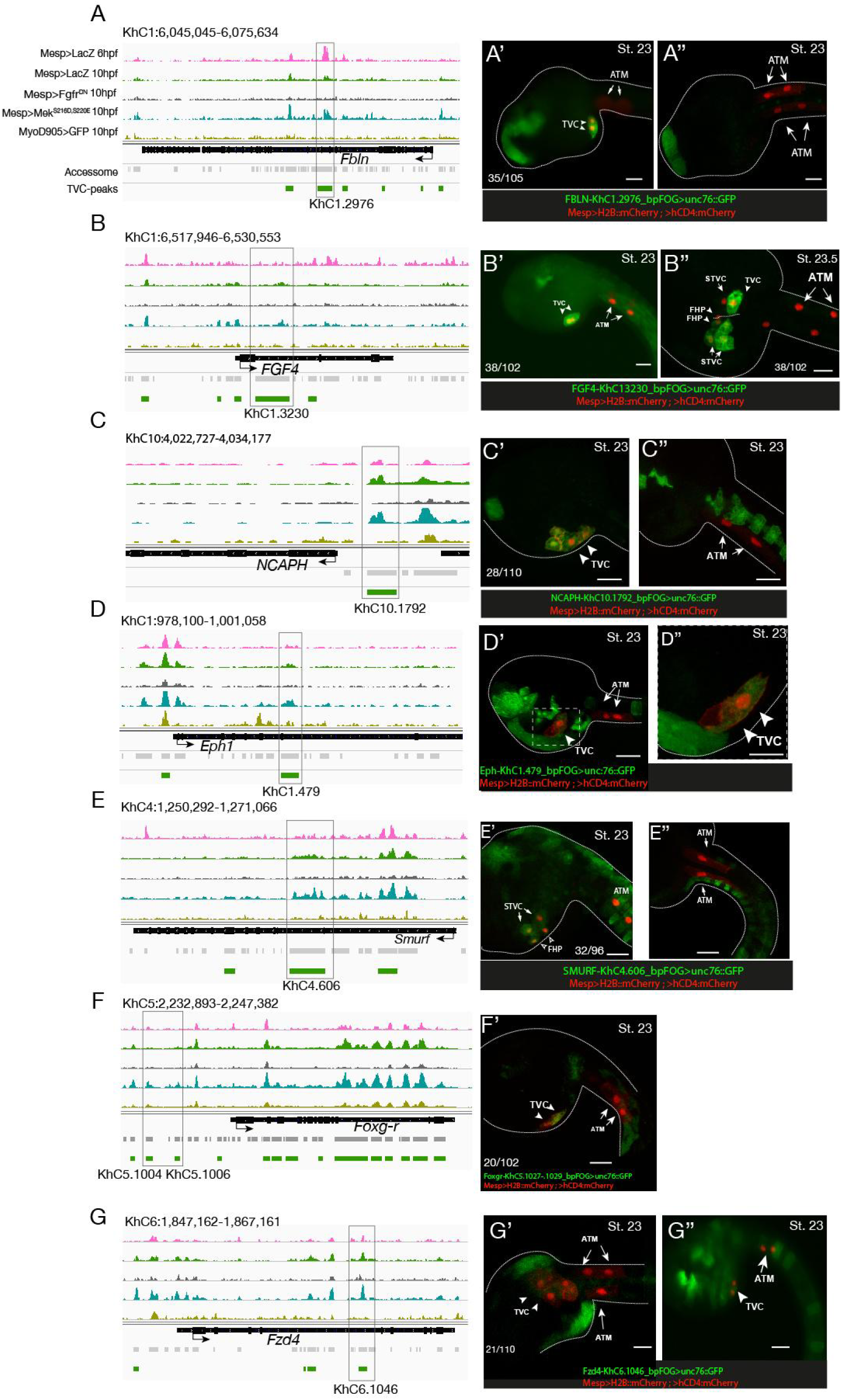
Candidate TVC-specific enhancers *in vivo* validation by reporter gene assay. (**A-G**) ATAC-seq peaks specifically accessible in TVC displaying chromatin accessibility profiles from ATAC-seq (normalized by total sequencing depth). Transcript model is indicated as black bar. The newly identified enhancers are in boxed regions. ATAC-seq peaks were tested *in vivo* by reporter gene assay (see Table 4). (A’-G’’) Reporter gene assay in embryos at stage 23 (Hotta et al., 2007). GFP expression was detected specifically in TVC cells and not in ATMs. Nuclei of B7.5 lineage cells are labelled by *Mesp>H2B::mCherry* and *Mesp>hCD4:mCherry* (red). Numbers indicate observed/total of half-embryos scored. Scale bar= 30 μm.

**Figure 2—figure supplement 4.**
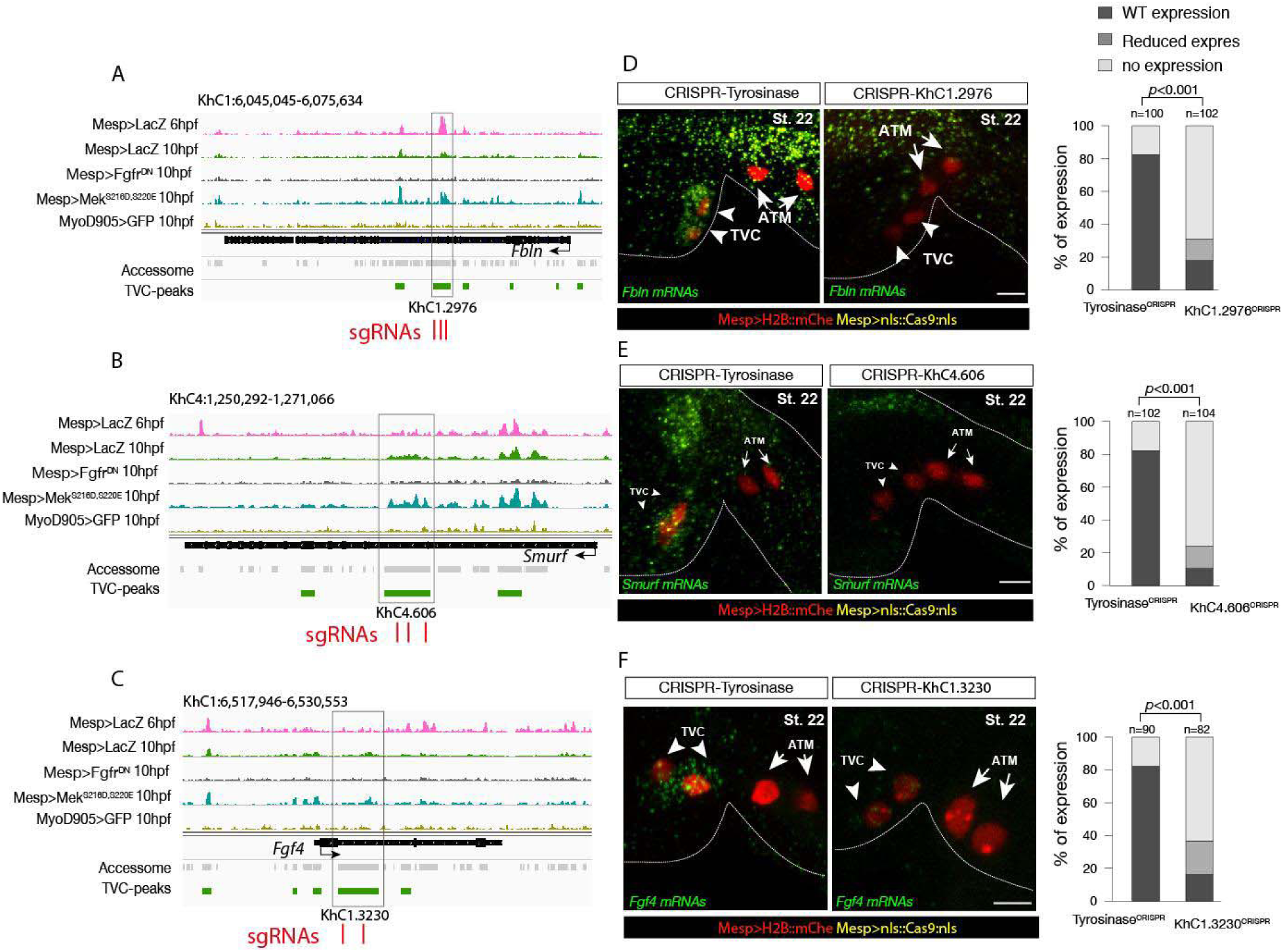
Candidate TVC-specific enhancers *in vivo* validation by CRISPR/Cas9. (**A-C**) TVC-specific enhancers in *Fbln*, *Smurf* and *Fgf4* loci targeted by two or three single guide RNA (sgRNAs) for CRISPR/Cas9-mediated deletions. Upper panel is showing chromatin accessibility profiles from ATAC-seq (normalized by total sequencing depth). Transcript model is indicated as black bar. (**D-F**) Endogenous expression of *Fbln* (D), *Smurf* (E) and *Fgf4* (F) visualized by *in situ* (green) in Tyrosinase^CRISPR^ (right panel) and upon CRISPR/Cas9-induced deletions of TVC-specific peaks (left panel). Nuclei of B7.5 lineage cells are labelled by *Mesp>nls::LacZ* and revealed with an anti beta-galactosidase antibody (red). Anterior to the left, stages indicated as “st.”. Experiment performed in μm. All the treatments were significant versus Tyrosinase (Fisher exact test, p < 0.001); “n” : total numbers of embryos.

**Figure 2—figure supplement 5.**
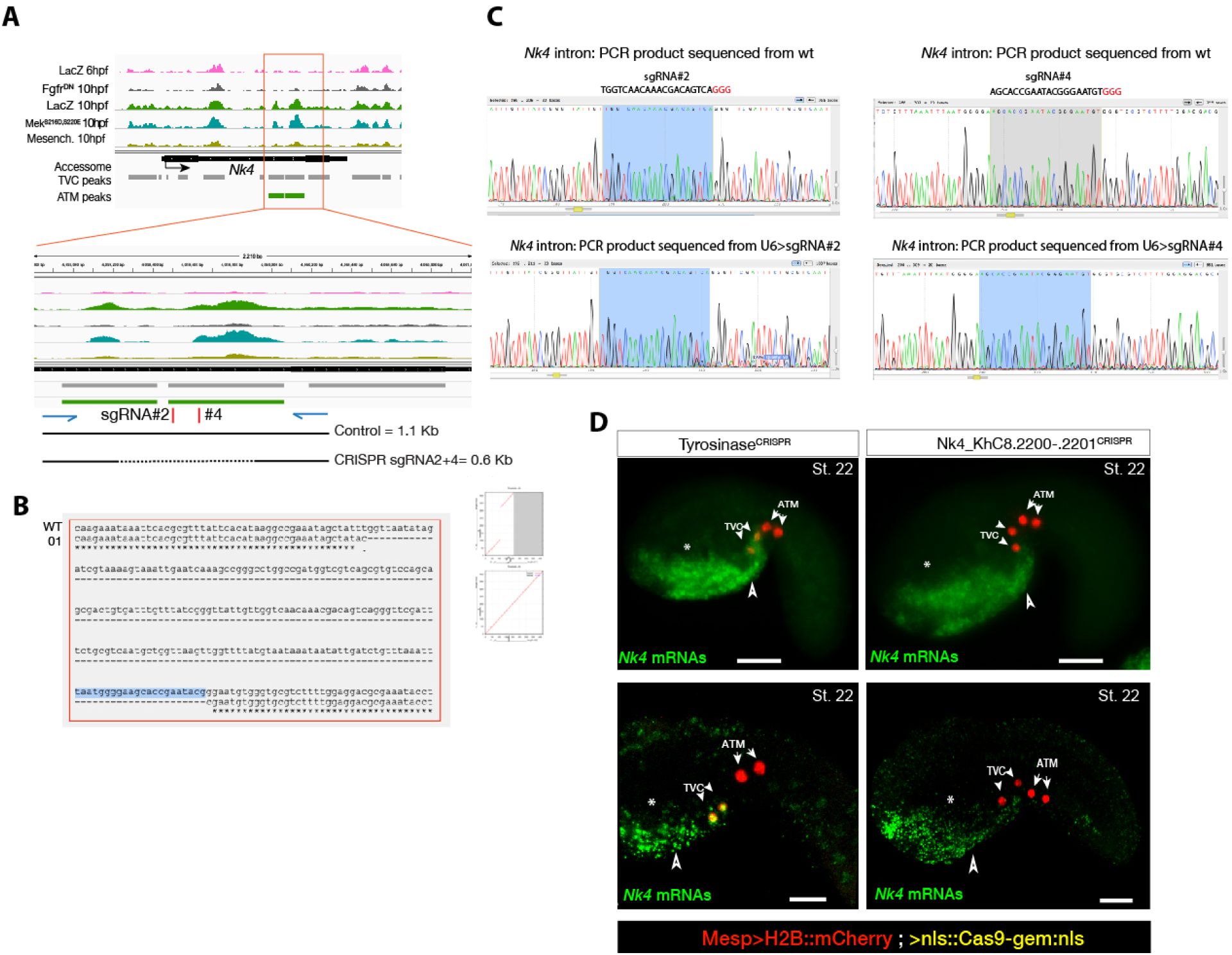
Peakshift validation of sgRNA efficiency. (**A**) IGV screenshot displaying chromatin accessibility profiles from ATAC-seq (normalized by total sequencing depth). Transcript model is indicated as black bar. The newly identified TVC-specific enhancer is in boxed regions; in the zoom region, blue arrows indicate primers used to amplify the region between the target sites. In wild-type embryos, the resulting PCR product is ~1.1 kilobase pairs. The two single guide RNAs used to target *Nk4/Nkx2-5* intronic element are in red. (**B**) Alignment of cloned PCR products amplified using the primers indicated in (A), from wild-type (wt) embryos, and from embryos electroporated with 25 µg *Ef1*α *>nls:Cas9-gem:nls* and 40 µg each of U6>sgRNA2 and U6>sgRNA4. Colonies 01 contains ~0.6 kilobase deletions between the approximate sites targeted by the two sgRNAs. (**C**) Quantification of indel-shifted electrophoresis chromatogram peaks (“Peakshift” assay (Gandhi et al., 2017)) revealed sgRNA mutagenesis efficacies. (**D**) *In situ* hybridization for *Nk4* (green) showing expression throughout the ventral head endoderm in embryos electroporated with Mesp>H2B::mCherry (red), Mesp>nls::Cas9-Gem::nls and U6>sgRNAs targeting Nk4 intron (Nk4_KhC8.2200-.2201^CRISPR^) or Tyrosinase (Tyrosinase^CRISPR^), used as control. In control embryos (panels on the left) *Nk4* expression is essentially wild-type with a strong expression in ventral head endoderm (asterix and arrowhead) and TVCs. In *Nk4_KhC8.2200-.2201^CRISPR^* embryos (panels on the right) *Nk4* expression is lost specifically in the B7.5 lineage and not in the other endogenous territories. Scale bars =30 µm.

**Figure 2—figure supplement 6.**
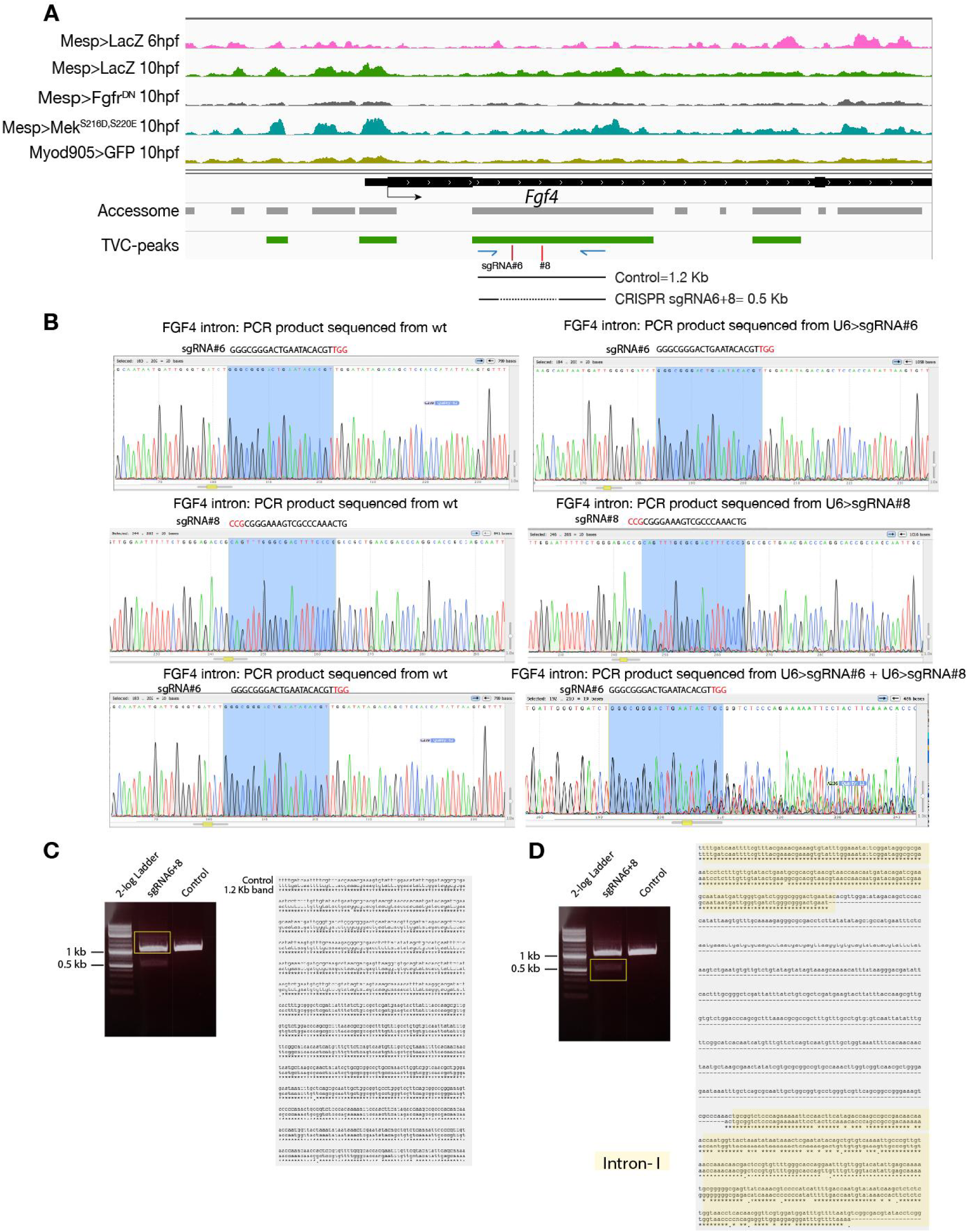
CRISPR validation on the TVC-specific *Fgf4* enhancer. (**A**) IGV screenshot displaying chromatin accessibility profiles from ATAC-seq (normalized by total sequencing depth). Transcript model is indicated as black bar. The newly identified TVC-specific enhancer is in boxed regions; in the zoom region, blue arrows indicate primers used to amplify the region between the target sites. In wild-type embryos, the resulting PCR product is ~1.2 kilobase pairs. The two single guide RNAs used to target *Fgf4* intronic element are in red. (**B**) Quantification of indel-shifted electrophoresis chromatogram peaks (“Peakshift” assay; (Gandhi et al., 2017)) revealed sgRNA mutagenesis efficacies. (**C-D**) 1% Agarose gel and alignment of cloned PCR products showing the result of *Fgf4* intron-I amplification with the oligos indicated in (A) from embryos electroporated with 25 µg *Ef*α*1>nls:Cas9-gem:nls* and 40 µg each of U6>sgRNA6 and U6>sgRNA8. (C) Alignment of cloned PCR products amplified using the primers indicated in (A), from the ~1.2 kilobase band that is similar to the control. (D) Alignment of cloned PCR products amplified using the primers indicated in (A), from the ~0.5 kilobase band that corresponds to the expected deletion between the approximate sites targeted by the two sgRNAs. The intronic element that is excluded from the sites targeted by the two sgRNAs remained intact (yellow box).

**Figure 3—figure supplement 1.**
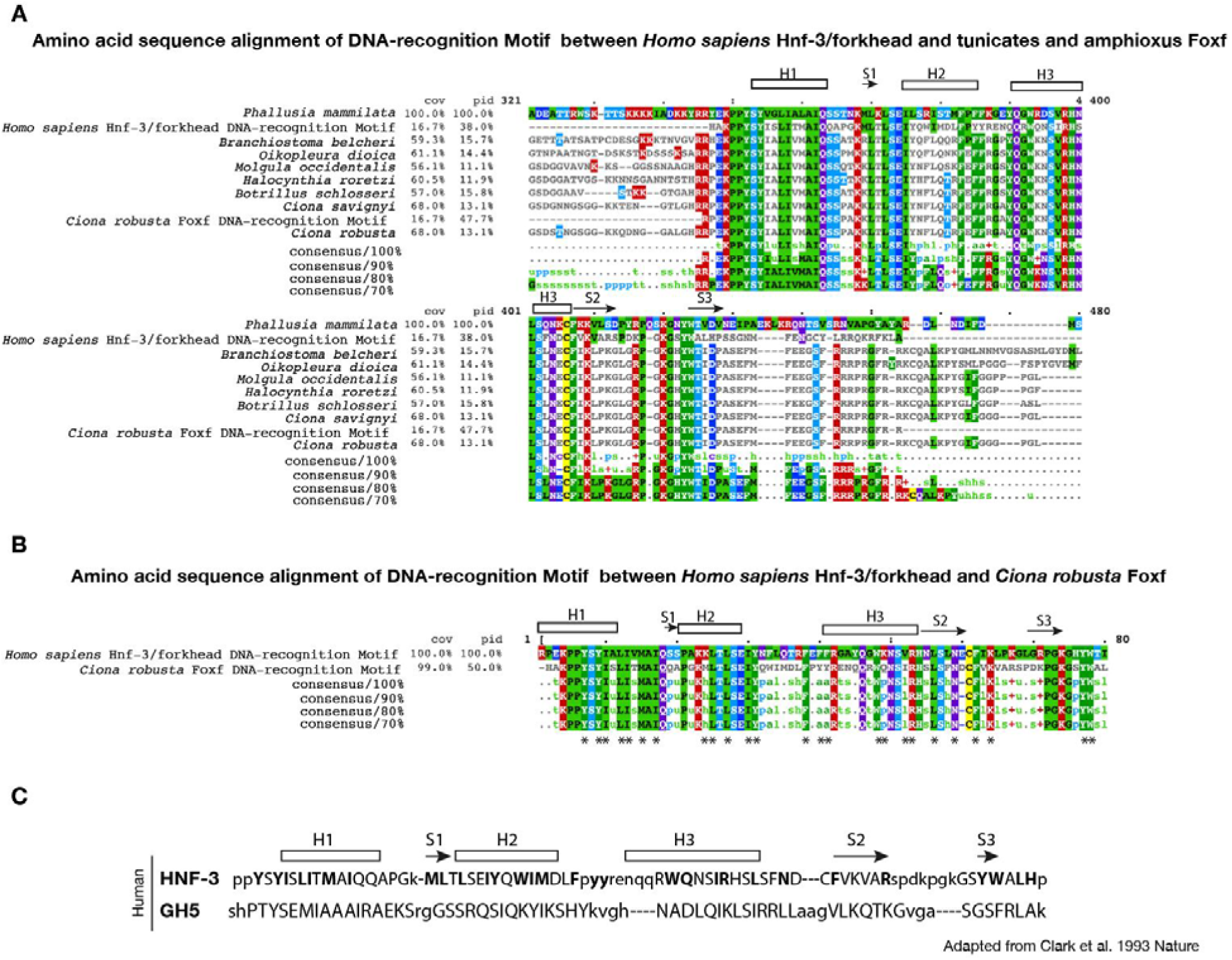
Conservation of DNA-recognition motifs of Foxf proteins. (**A**) Conservation of DNA-recognition motifs among *Homo sapiens* Hnf-3/forkhead and tunicates and amphioxus Foxf proteins. The protein sequences were retrieved from NCBI (www.ncbi.nlm.nih.gov), Ensembl (www.ensembl.org) and Aniseed (www.aniseed.cnrs.fr/aniseed) databases using Foxf of *Ciona robusta* as query sequence (see Supplementary File-1). Alignment of conserved amino-acid residues in Foxf orthologues was obtained using Clustal Omega (MView algorithm) (McWilliam et al., 2013). (**B**) Amino acid sequence alignment of DNA-recognition motif between *Homo sapiens* Hnf-3/forkhead (AAT74628.1) and *Ciona robusta* Foxf (NP_001071710.1). Asterisks indicate the structurally equivalenced residues between HFN3□ and GH5 as in (Clark et al., 1993). (**C**) Aminoacid sequence alignment in HFN3□ (107-223) and GH5 based on structural alignment and secondary structural elements indicated as rectangle for □-helices and arrows for β-strand (structurally equivalenced residues are indicated in upper case) adapted from (Clark et al., 1993).

**Figure 3—figure supplement 2.**
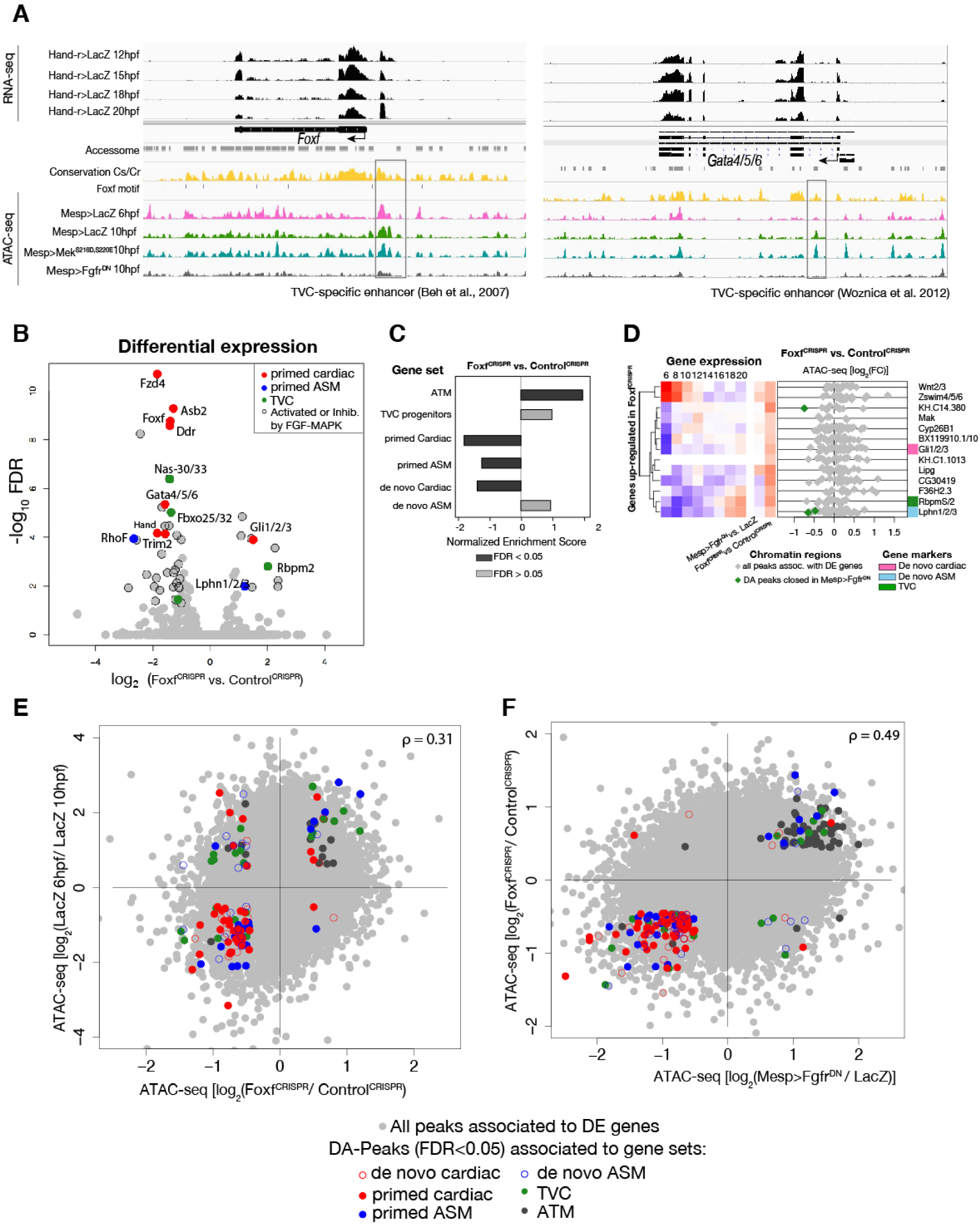
General characterization of Foxf ^CRISPR^. (**A**) ATAC-seq peaks displaying *Foxf* and *Gata4/5/6* loci with expression profiles from RNA-seq and chromatin accessibility profiles from ATAC-seq (normalized by total sequencing depth). Known TVC-specific enhancers of *Foxf* (Beh et al., 2007) and *Gata4/5/6* (Woznica et al., 2012) are in the boxed regions. (**B**) Differential gene expression determined by RNA-seq analysis of Foxf^CRISPR^ vs. Control^CRISPR^ at 10hpf. Colored dots represent genes that are differentially expressed (FDR<0.05) in Foxf^CRISPR^ vs. Control^CRISPR^ at 10hpf. Genes not differentially expressed are shown in gray. (**C**) GSEA showing normalized enrichment score (NES) of gene sets in peaks ranked by difference in accessibility between Foxf^CRISPR^ vs. Control^CRISPR^ at 10hpf. Dark gray bars indicate significant enrichment. Light gray bars are not significant (FDR < 0.05). (**D**) Change in accessibility between ATAC-seq samples. Colored dots represent peaks associated to genes that are differentially accessible (FDR<0.05 and |log2 FC| > 0.5). (**E**) Association between expression and accessibility of Foxf up-regulated genes and peaks which were both closed in Foxf^CRISPR^ and associated to TVC-specific peaks as in Figure 2C. Gene expression data for 6hpf and “FGF-MAPK perturbation 10hpf” (Christiaen et al., 2008), and from 8 to 20hpf (Razy-Krajka et al., 2014) were previously published.

**Figure 3—figure supplement 3.**
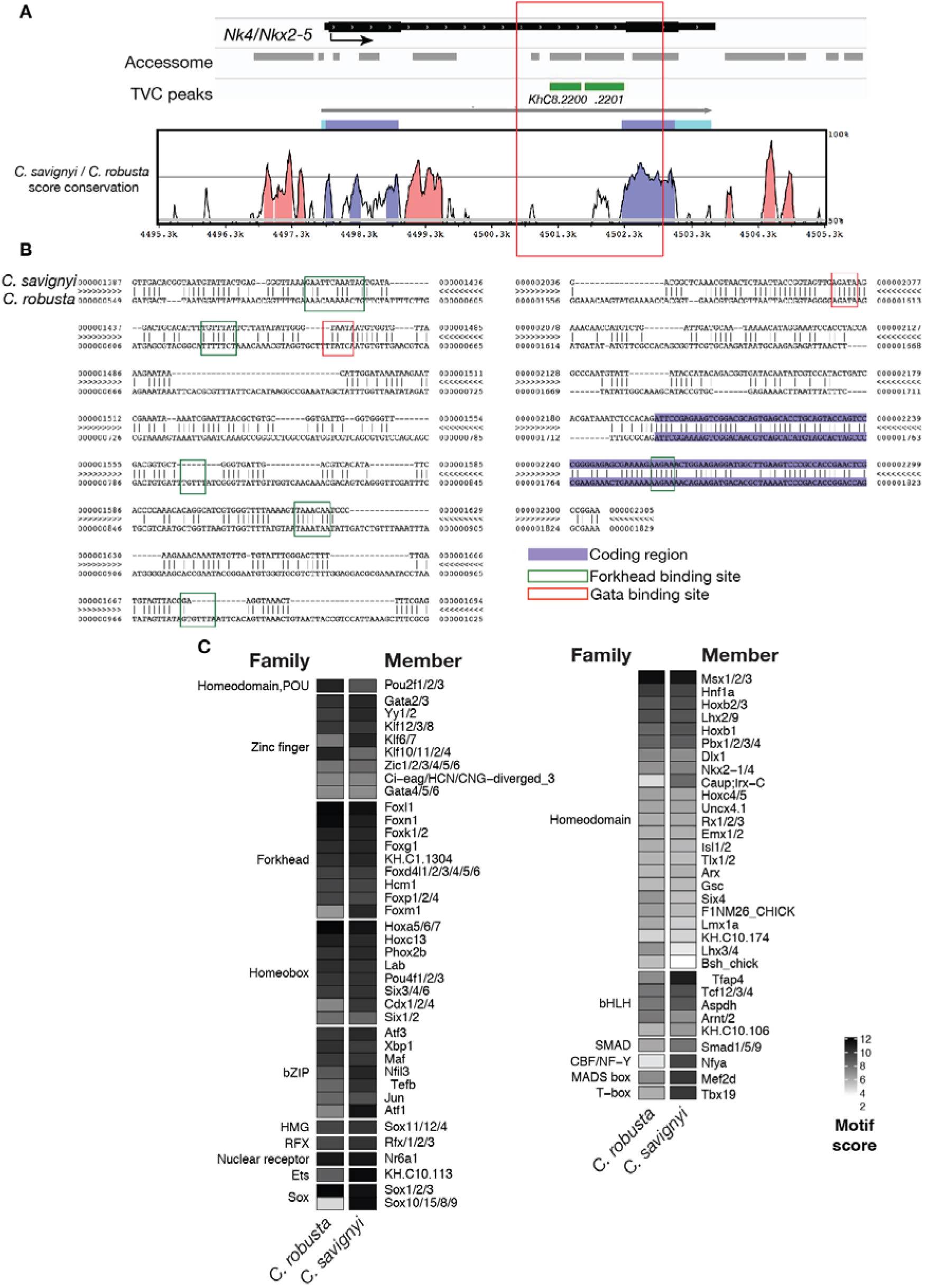
Conserved binding motifs in TVC-specific *Nk4/Nkx2-5* enhancer. (**A**) Conservation of TVC-accessible elements of *Nk4/Nk2-5* locus between *Ciona robusta* and *Ciona savignyi*. The transcript model is shown in black. Conservation score was calculated using mVISTA (http://genome.lbl.gov/vista/mvista/submit.shtml). Pink peaks indicate conserved non-coding sequences (>65% identity per 80 bp). (**B**) Alignment of DNA region corresponding to “*KhC8.2201*” peak between *Ciona savignyi* and *Ciona robusta* (bottom sequence). Only the core nucleotides are shown for putative Forkhead and Gata binding sites. (**C**) Match scores for TF binding motifs present in the boxed region (~1.2 Kb) including the I-intronic element of both *Ciona robusta* and *Ciona savignyi Nk4/Nk2-5* transcripts. Only the highest scoring match for each TF is shown.

**Figure 3—figure supplement 4.**
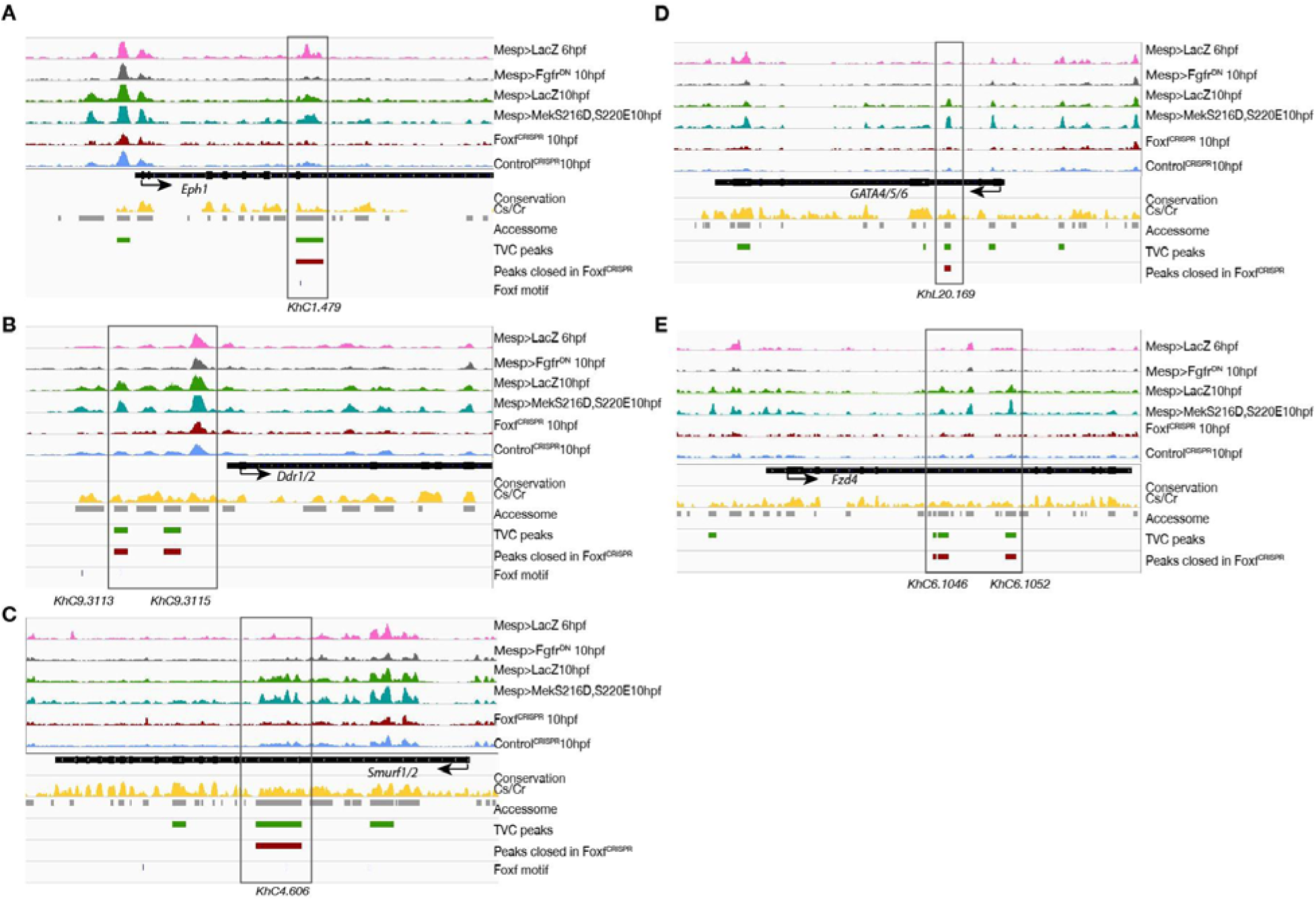
Foxf loss-of-function (Foxf^CRISPR^) caused closing of TVC-specific enhancers. **(A-E)** *Foxf* target gene loci chromatin accessibility profiles showing ATAC-seq (normalized by total sequencing depth) in the indicated conditions. Foxf core binding sites (GTAAACA) are displayed as blue lines. The transcript model is shown in black.

**Figure 3—figure supplement 5.**
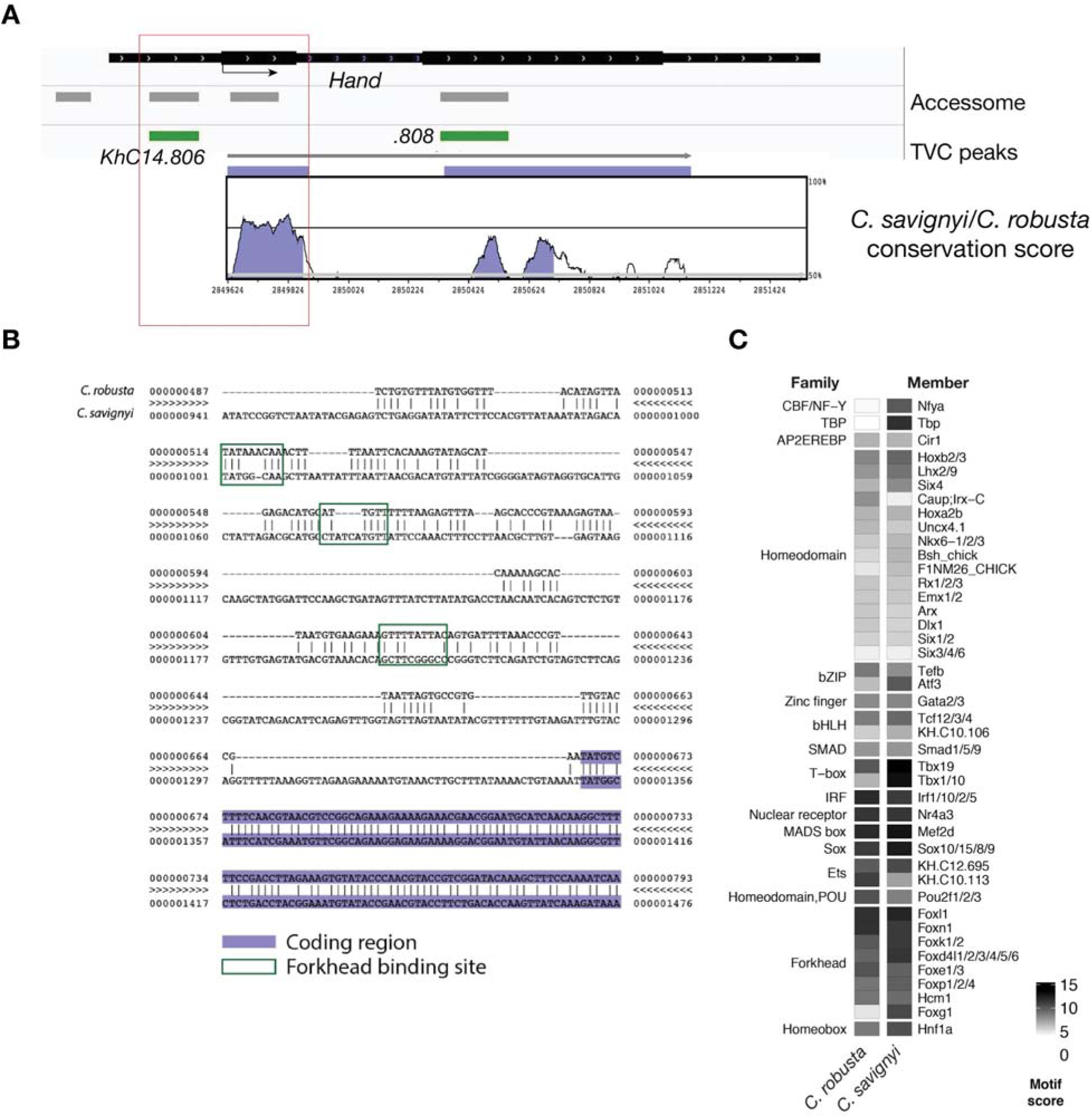
Conserved binding motifs in TVC-specific *Hand* enhancer. (**A**) Conservation of TVC-accessible elements of *Hand* locus between *Ciona robusta* and *Ciona savignyi*. The transcript model is shown in black. Conservation score was calculated using mVISTA (http://genome.lbl.gov/vista/mvista/submit.shtml). Violet peaks indicate conserved sequences (>65% identity per 80 bp). (**B**) Alignment of DNA region corresponding to “KhC14.806” peak between *Ciona robusta* and *Ciona savignyi* (bottom sequence). Putative Forkhead binding sites are evidenced in green box (only the core nucleotides are shown). (**C**) Match scores for TF binding motifs present in the boxed region (~0.8 Kb) upstream of the coding ATG in both *Ciona robusta* and *Ciona savignyi Hand* transcripts. Only the highest scoring match is shown for each TF.

**Figure 3—figure supplement 6.**
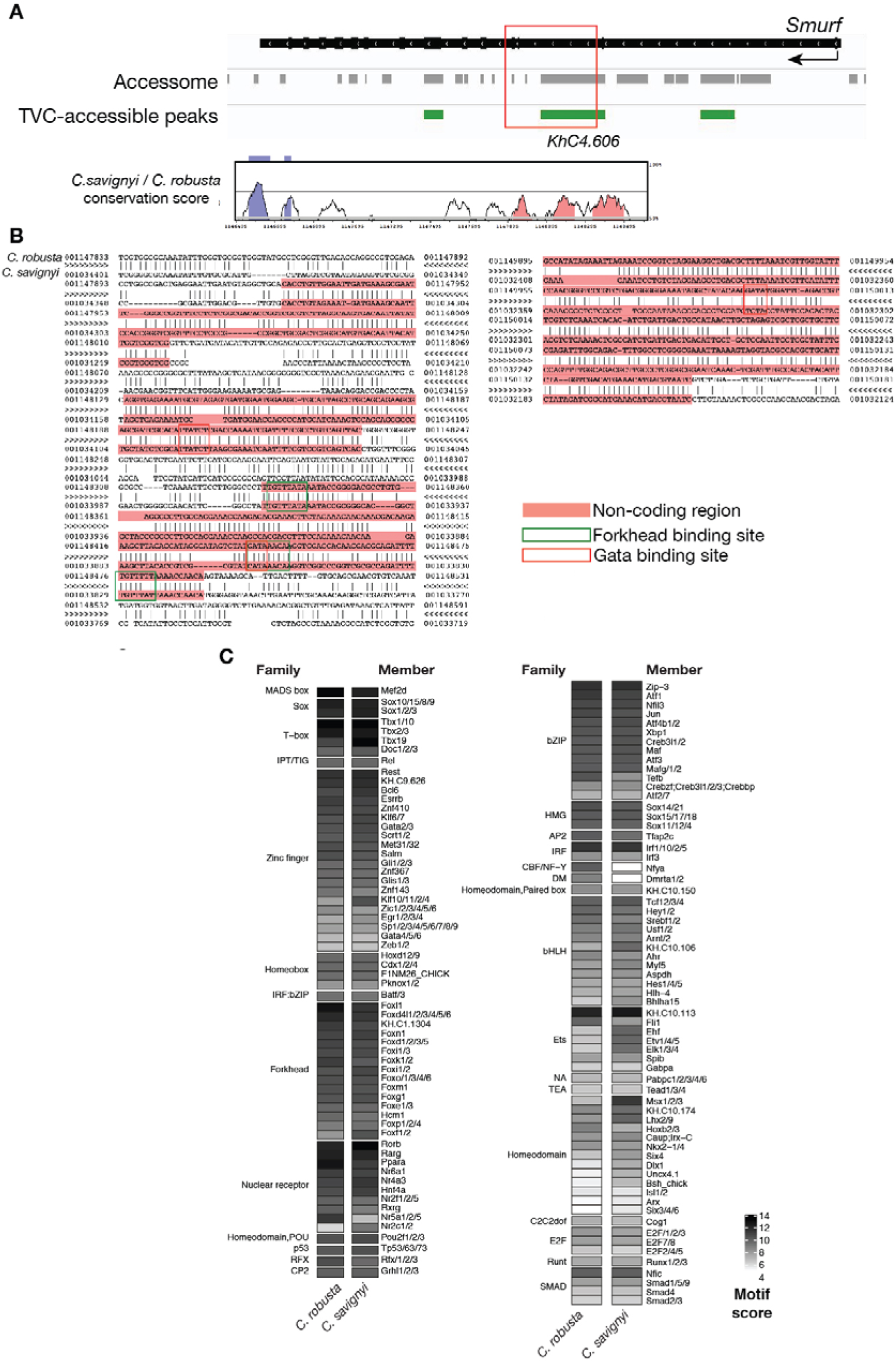
Conserved binding motifs in *Smurf* enhancer. (**A**) Conservation of TVC-accessible elements of *Smurf* locus between *Ciona robusta* and *Ciona savignyi*. The transcript model is shown in black. Conservation score was calculated using mVISTA (http://genome.lbl.gov/vista/mvista/submit.shtml). Pink peaks indicate conserved non-coding sequences (>65% identity per 80 bp). (**B**) Alignment of DNA region corresponding to “*KhC4.606*” peak between *Ciona robusta* and *Ciona savignyi* (bottom sequence). Only the core nucleotides are shown for putative Forkhead and GATA binding sites. (**C**) Match scores for TF binding motifs present in the boxed region including the II-intronic element and exon III-IV of both *Ciona robusta* and *Ciona savignyi Smurf* transcripts. Only the highest scoring match is shown for each TF.

**Figure 3—figure supplement 7.**
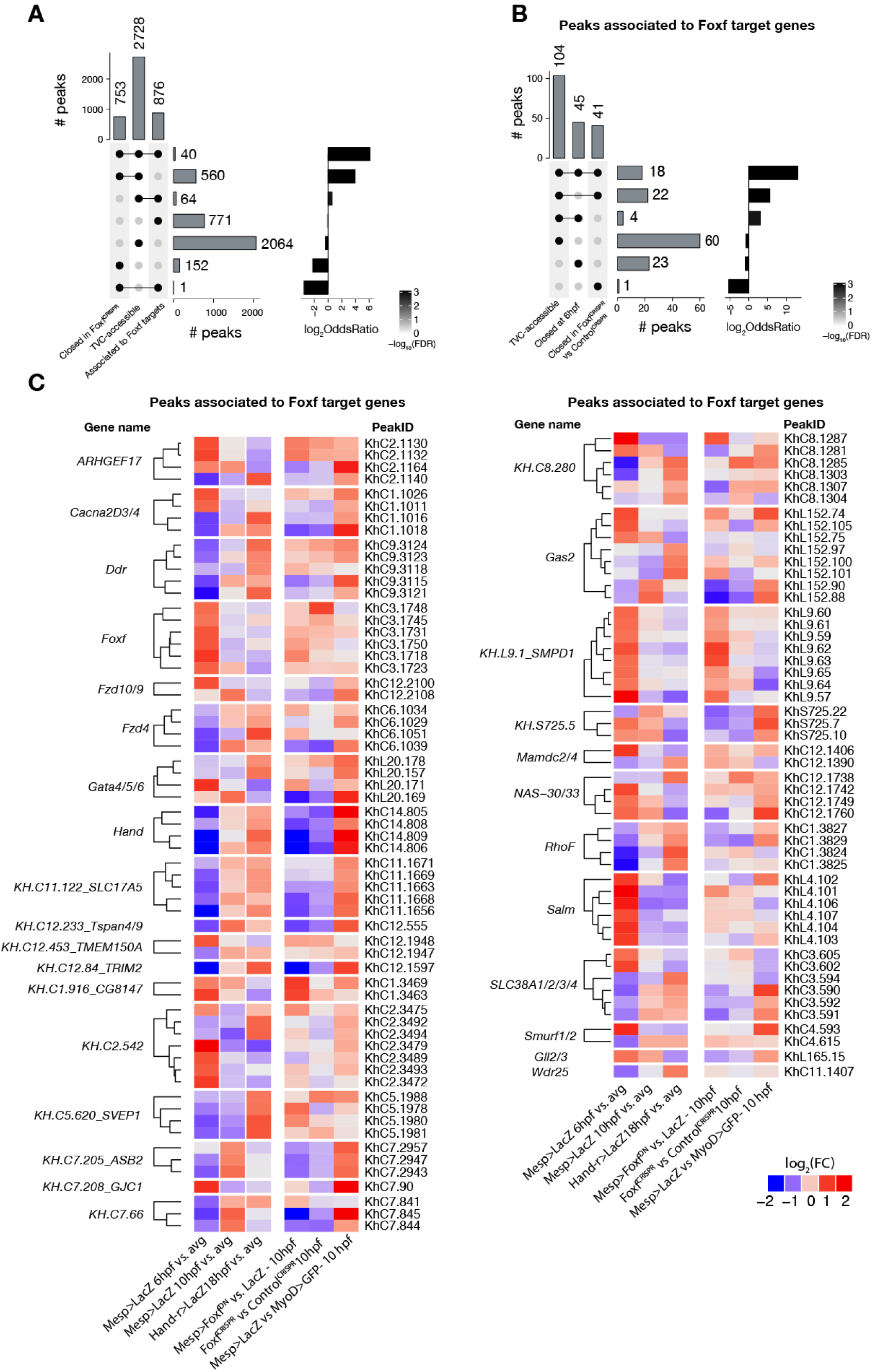
Accessible elements annotated to *Foxf* target genes. **(A)** Intersection of elements associated to *Foxf* target genes at 10 hpf (see Material and Methods) and differentially accessible elements. Peaks closed in Foxf^CRISPR^ are peaks more accessible in Control^CRISPR^ vs Foxf^CRISPR^ at 10hpf. A binomial test was performed on each intersection against the null hypothesis that the intersection’s constituent sets are independent (see Statistics under Materials & Methods). **(B)** Subset of (A) showing only elements associated to *Foxf* target genes. Peaks closed at 6hpf are peaks more accessible in Mesp>LacZ 10hpf vs Mesp>LacZ 6hpf. **(C)** ATAC-seq peaks (113 regions) associated to Foxf target genes showing differential accessibility (FDR<0.05, see Material and Methods) over time. The log_2_FC for each time point is versus the average (avg) of all control samples.

**Figure 4—figure supplement 1.**
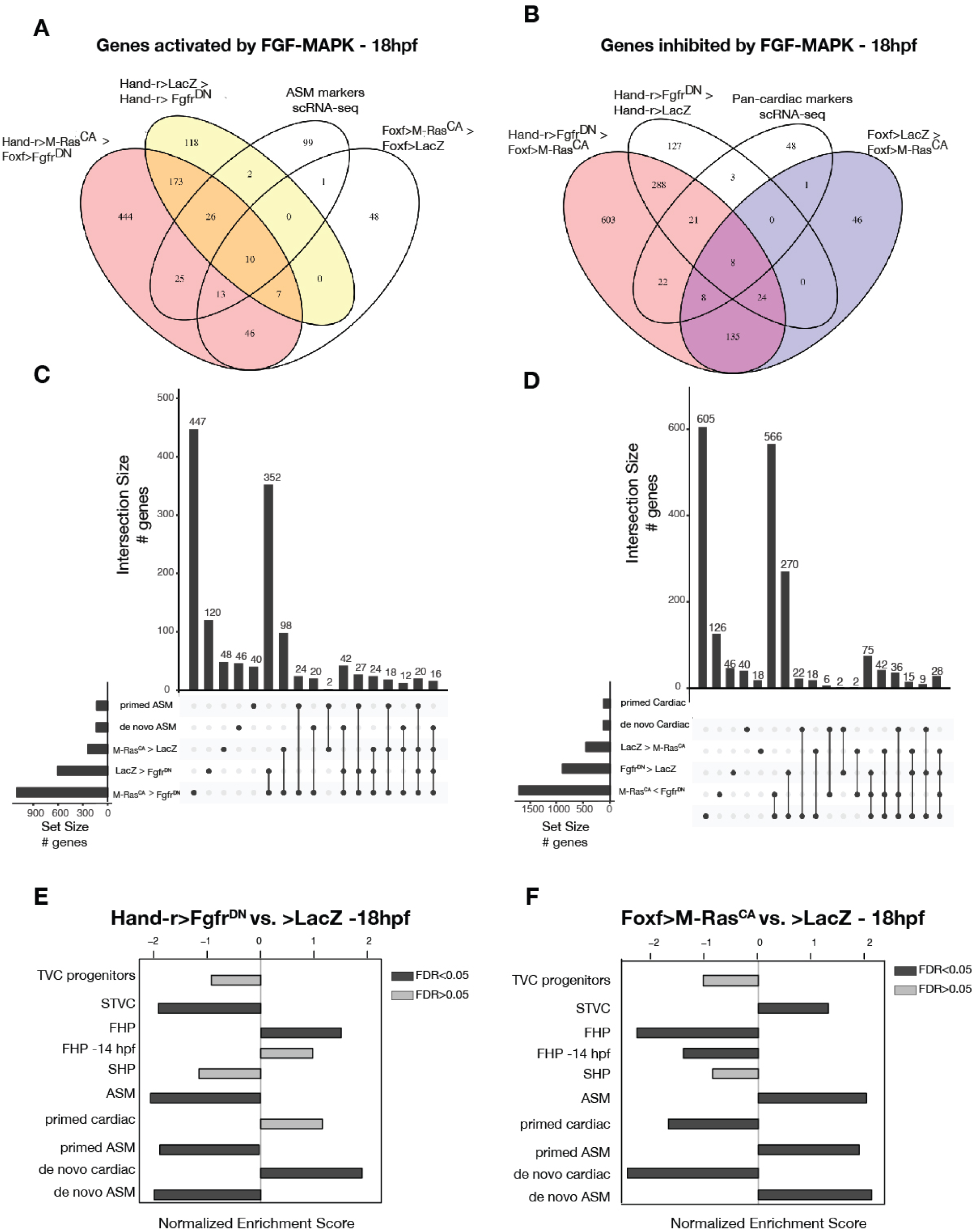
General characterization of FGF-MAPK perturbation. (**A-D**) Intersections of differentially expressed genes from bulk RNA-seq and scRNA-seq(Wang et al., 2019) comparisons in cardiopharyngeal restricted cells using Venn Diagram (A-B) and UpSet plots (C-D). FGF-MAPK activated genes at 18hpf are defined as the intersection of genes downregulated in Fgfr^DN^ vs. LacZ at 18 hpf and genes downregulated in Fgfr^DN^ vs. M-Ras^CA^ at 18 hpf (A). FGF-MAPK inhibited genes at 18hpf are defined as the intersection of genes downregulated in M-Ras^CA^ vs. LacZ at 18hpf and genes downregulated in M-Ras^CA^ vs. Fgfr^DN^ at 18hpf (FDR < 0.05 & |log2FC| > 1). (**E-F**) GSEA of significantly enriched gene sets shows opposite trends in Fgfr^DN^ vs. control and M-Ras^CA^ vs. control (LacZ). Dark gray bars indicate significant enrichment. Light gray bars are not significant (FDR < 0.05).

**Figure 4—figure supplement 2.**
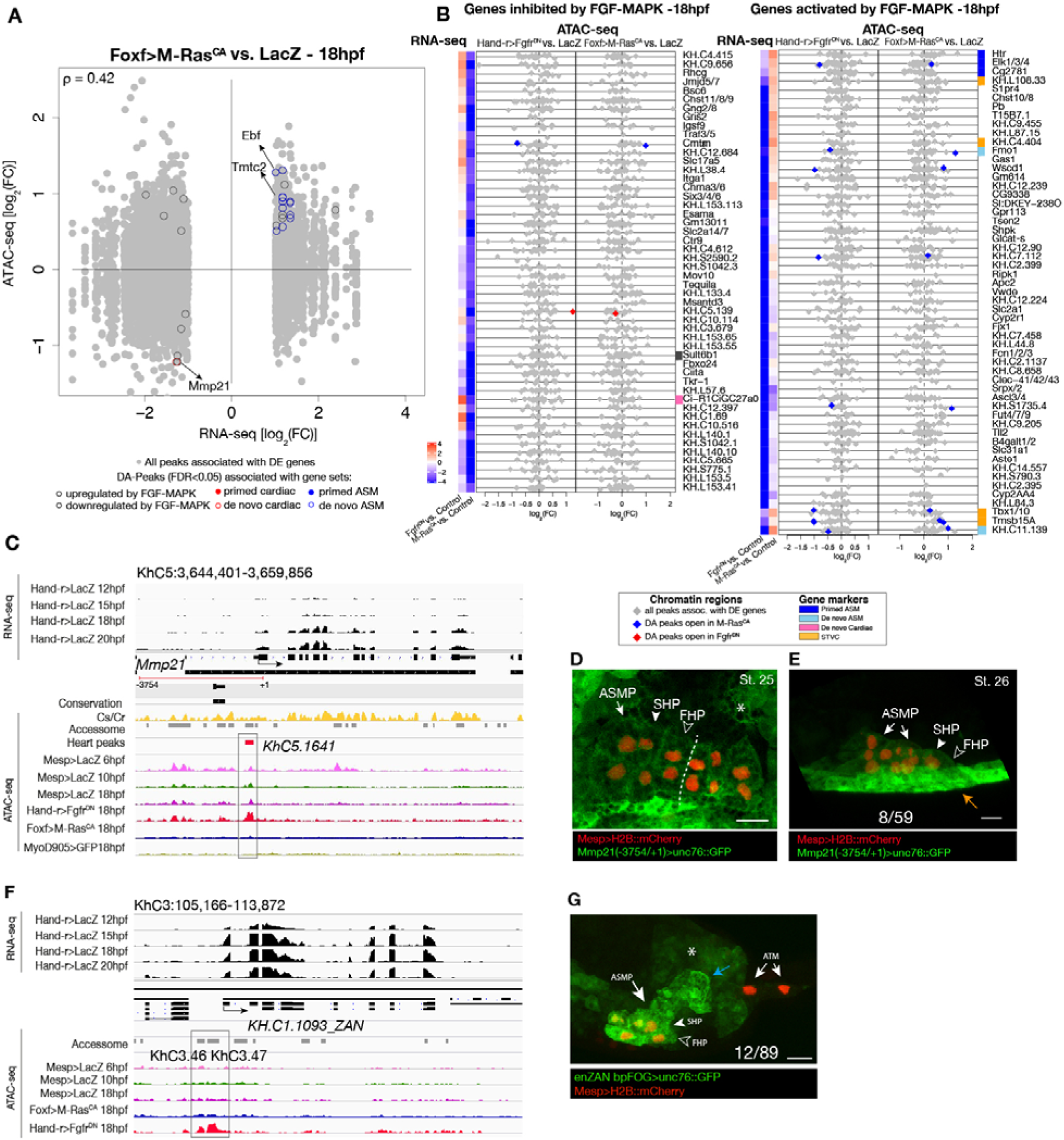
Differential accessibility in response to FGF/MAPK perturbation at 18hpf. (**A**) Differentially expressed (DE) genes vs differentially accessible (DA) peaks using bulk RNA-seq and ATAC-seq in Foxf>M-Ras^CA^ vs. LacZ at 18hpf. (**B**) Relationship between expression and accessibility of DE genes associated to DA peaks. The top axis shows the scale for log_2_FC of bulk RNA-seq (black dots). The bottom axis shows the log_2_FC scale for ATAC-seq (colored diamonds). Top 50 genes inhibited and activated by FGF-MAPK (based on log_2_FC) along with STVC genes. Heatmaps show gene expression over time. (**C**) A 15 kb region on chromosome 5 displaying expression profiles from RNA-seq and chromatin accessibility profiles from ATAC-seq (normalized by total sequencing depth). Transcript model is indicated as black bars, percentage of conservation score between *C. savignyi* and *C. robusta* obtained obtained from WASHU browser(Brozovic et al., 2018) (yellow peaks), accessome (light grey bars), TVC-(green bar) and ATM-specific peaks (dark gray bars). Peak accessible specifically in the heart (*KhC5.1641*) of *Mmp21* locus is in the boxed region. (**D-E)** Enhancer driven *in vivo* reporter expression (green) of a ~3kb region upstream the coding ATG and including the DA peak (KhC5.1641). (D) dorsal view, (E) lateral view; GFP is detected in the mesenchyme (asterisk) surrounding the cardiopharyngeal progenitors (D) and in the epidermis (orange arrow) (E). B7.5 cells are marked with Mesp>H2B::mCherry (red). Dotted line: ventral midline. Numbers indicate observed/total of half-embryos scored. Scale bar = 25 µm.

**Figure 4—figure supplement 3.**
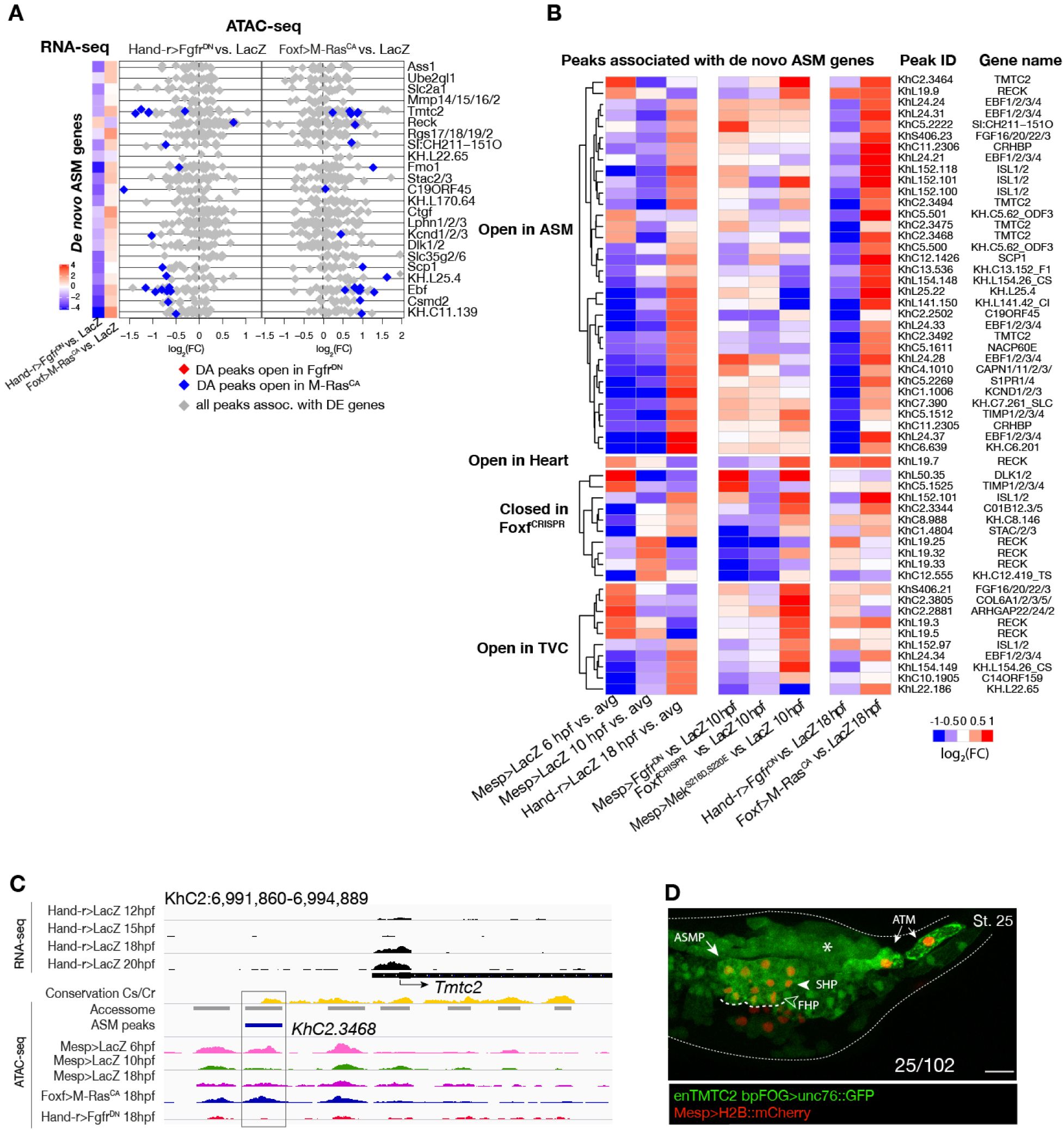
Accessibility of elements annotated to *de novo* ASM genes. (**A**) Relationship between accessibility and expression of *de novo* ASM genes in either condition indicated. (**B**) Time-dependent ATAC-seq peaks associated to *de novo* expressed ASM genes. The accessibility of these peaks is shown for 6, 10 and 18hpf vs. the average (avg.) accessibility in the controls and upon FGF-MAPK perturbation either at 10 and 18hpf. Peaks are clustered based on their accessibility as “Open in ASM” less accessible in Hand-r>Fgfr^DN^ vs. Foxf>M-Ras^CA^ or Hand-r>LacZ at 18hpf, “Open in Heart” less accessible in Foxf>M-Ras^CA^ vs Hand-r>Fgfr^DN^ or Hand-r>LacZ at 18hpf, “Closed in Fofx^CRISPR^” less accessible in Foxf^CRISPR^ vs. Control^CRISPR^, “Open in TVC” less accessible in Mesp>Fgfr^DN^ vs. Mesp>Mek^S216D,S220E^ or Mesp>LacZ at 10 hpf. Only peaks changing accessibility between 6hpf and 10hpf or 10hpf and 18hpf are shown.

**Figure 4—figure supplement 4.**
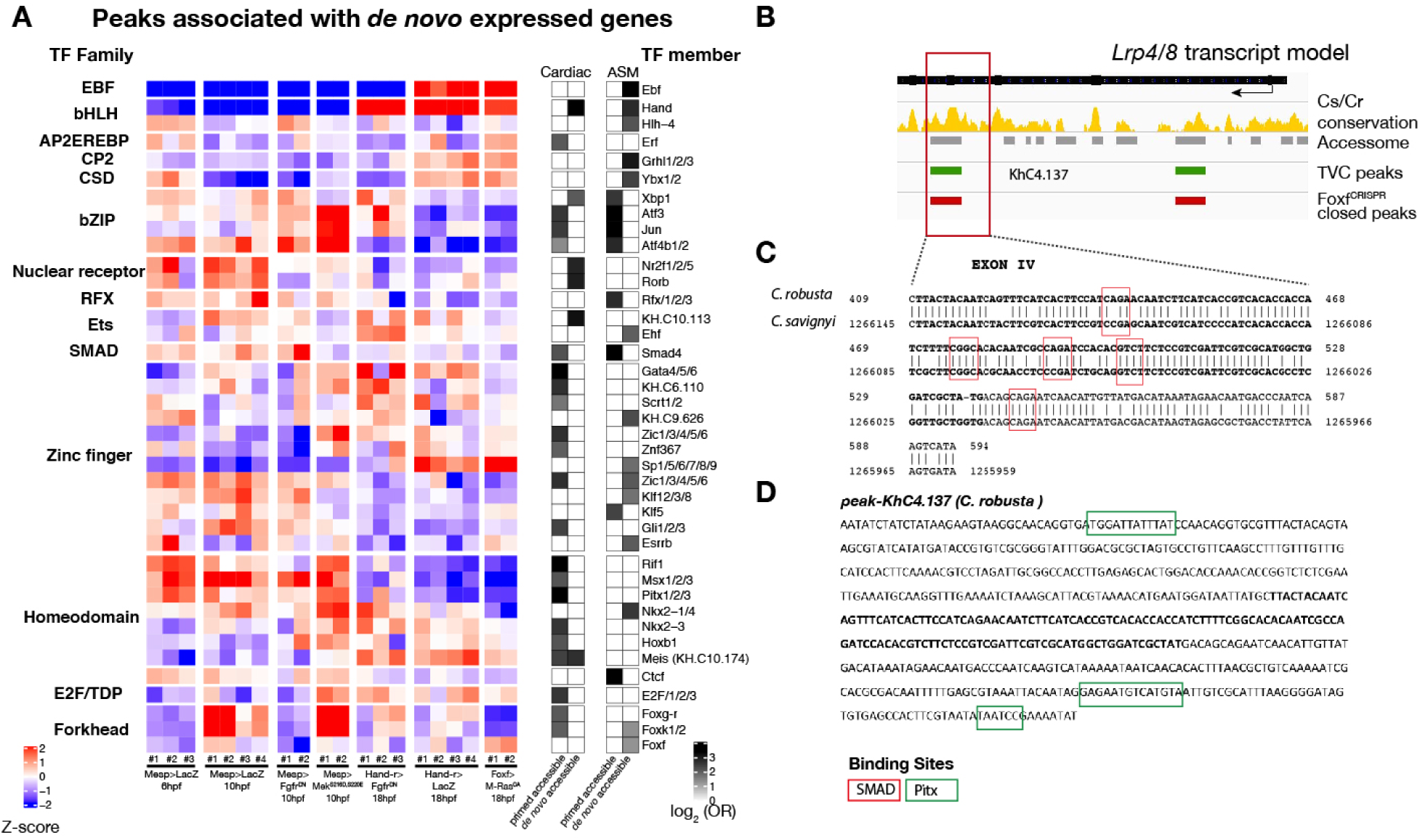
Accessibility of binding motifs over time for elements annotated to *de novo*-expressed genes. (**A**) TF binding motifs enriched in peaks associated to *de novo* cardiac and pharyngeal expressed genes parsed based on primed and *de novo* accessibility (see Material and Methods). log_2_ odds ratios (see Materials and Methods) are shown for motifs significantly enriched (one-tailed hypergeometric test, *p* < 0.05) in the indicated peak classes. Motif accessibility from chromVAR (Schep et al., 2017) is shown for all peaks associated to a *de novo*-expressed cardiac or ASM gene. Only the motif with the highest log_2_OR for each TF is shown. (**B**) Conservation of TVC-accessible peaks closed in Foxf^CRISPR^ in *Lrp4/8* locus between *C. savignyi* and *C. robusta*. The gene body is shown in black. Conservation score was obtained from WASHU browser (Brozovic et al., 2018). (**C**) Alignment of conserved region of “*KhC8.137*” peak between *C. robusta* and *C. savignyi*. (**D**) *C. robusta Hand* locus, exon IV sequence highlighted in bold and black. Only the core nucleotides are shown for candidate Pitx binding sites.

**Figure 5—figure supplement 1.**
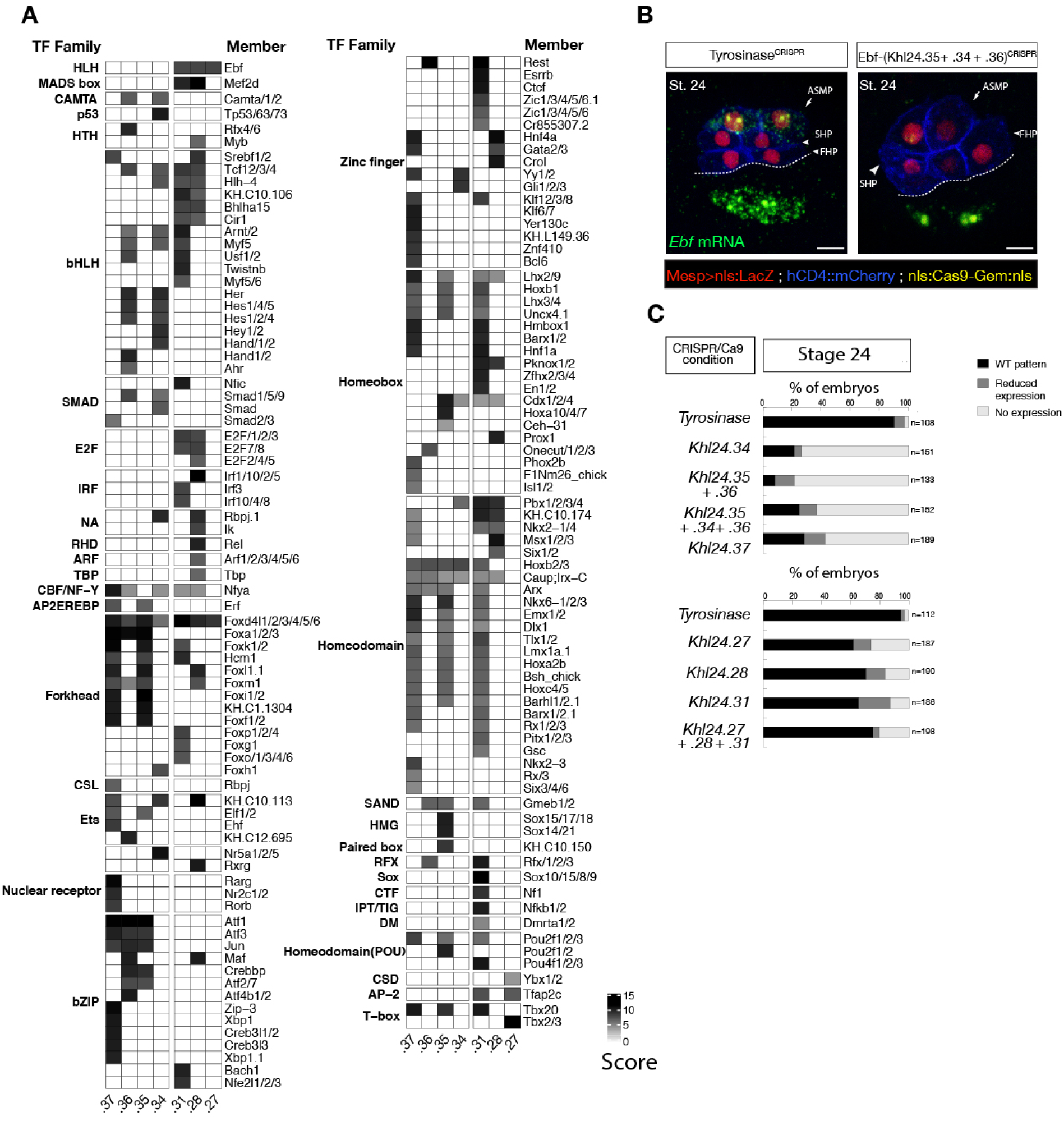
*Ebf* regulatory regions showing differentially accessibility over time contain distinct binding motifs. (**A**) Motif scores in each experimentally validated peak in the *Ebf* locus. Only the highest match score is shown for each motif. (**B**) Endogenous expression of Ebf visualized by in situ (green) in Tyrosinase^CRISPR^ and upon CRISPR/Cas9-induced deletions of TVC-specific peaks at stage 24 according to (Hotta et al., 2007). Nuclei of B7.5 lineage cells are labelled by Mesp>nls::LacZ and revealed with an anti beta-galactosidase antibody (red). Nuclei of B7.5 lineage cells are labelled by Mesp>nls::LacZ and revealed with an anti beta-galactosidase antibody (red). Scale bar = 10µm. (**C**) Proportions of larvae halves showing the indicated *Ebf* transcription patterns, in indicated experimental conditions. Experiment performed in biological replicates. All the treatments were significant versus Control (*Tyrosinase*) (Fisher exact test, *p* < 0.001).

**Figure 5—figure supplement 2.**
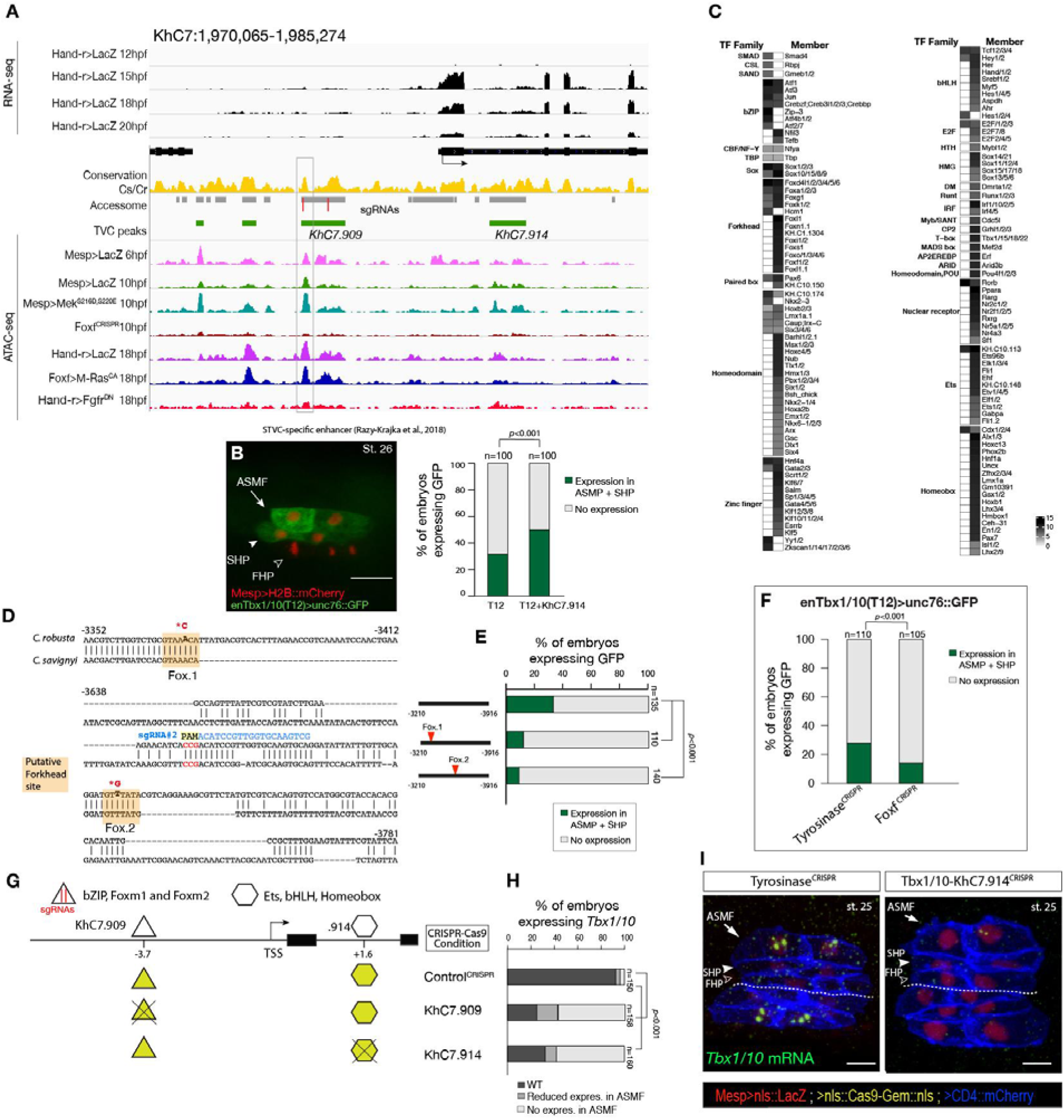
Combinations of *cis*-regulatory elements with distinct chromatin accessibility profiles are required for *Tbx1/10* transcription in pharyngeal-muscle precursors. (**A**) An 11 kb region on chromosome 7 displaying expression profiles of RNA-seq and chromatin accessibility profiles ATAC-seq normalized tag count in *Tbx1/10* locus. (**B**) STVC-specific enhancer (T12) (Razy-Krajka et al., 2018b) driven *in vivo* reporter expression (green) in ASMFs and SHPs at stage 26 (Hotta et al., 2007). Nuclei of B7.5 lineage cells are labelled by Mesp>H2B::mCherry (red). Scale bar, μm. T12 enhancer tested alone or fused to the intronic element (T12+*KhC7.914*). Statistical significance of the difference in reporter expression was tested using a Fisher exact test (*p* < 0.001) (**C**) Motif scores in each experimentally validated peak in the *Tbx1/10* locus. Only the highest match score is shown for each motif. **(D)** Sequence alignment of *Tbx1/10* enhancer (T12) between *Ciona robusta*/*Ciona savignyi*. Conserved blocks in the orange boxes with putative Forkhead binding sites. In blue is highlighted the single guide RNA (sgRNA#2) used to target CRISPR/Cas9 system, with the PAM domain in red; the point mutations induced in two conserved putative Forkhead binding sites (Fox1 and Fox2) are in bold and red after the asterisks. **(E)** Proportion of larvae expressing both GFP and mCherry in the STVC progeny when co-electroporated wild-type and mutant *Tbx1/10* reporters lacking the indicated putative Forkhead binding sites and Mesp > H2B::mCherry in comparison to the control. **(F)** Proportions of larvae halves showing GFP expressed in the ASMFs and SHPs in embryos electroporated with Mesp>Cas9 along with single guide RNAs targeting Tyrosinase (control^CRISPR^) as well as Foxf (Foxf^CRISPR^). (**G**) Schematic representation of regulatory elements in *Tbx1/10* locus as displayed in ATAC-seq profiles targeted for CRISPR/Cas9-mediated deletions. (**H**) Proportions of larvae halves showing the indicated *Tbx1/10* transcription patterns, in indicated experimental conditions (Fisher exact test, *p* < 0.001). (**I**) Endogenous expression of *TBX1/10* visualized by *in situ* (green) in Tyrosinase^CRISPR^ (left panel) and upon CRISPR/Cas9-induced deletion of TVC-specific peaks (right panel) at stage 25 according to (Hotta et al., 2007). Nuclei of B7.5 lineage cells are labelled by Mesp>nls::LacZ and revealed with an anti beta-galactosidase antibody (red). Nuclei of B7.5 lineage cells are labelled by Mesp>nls::LacZ and revealed with an anti beta-galactosidase antibody (red). Scale bar = 10µm. Experiment performed in biological replicates. Total numbers of individual halves scored per condition are shown in ‘n=’. All the treatments were significant versus Control (*Tyrosinase*) (Fisher exact test, *p* < 0.001).

**Figure 5—figure supplement 3.**
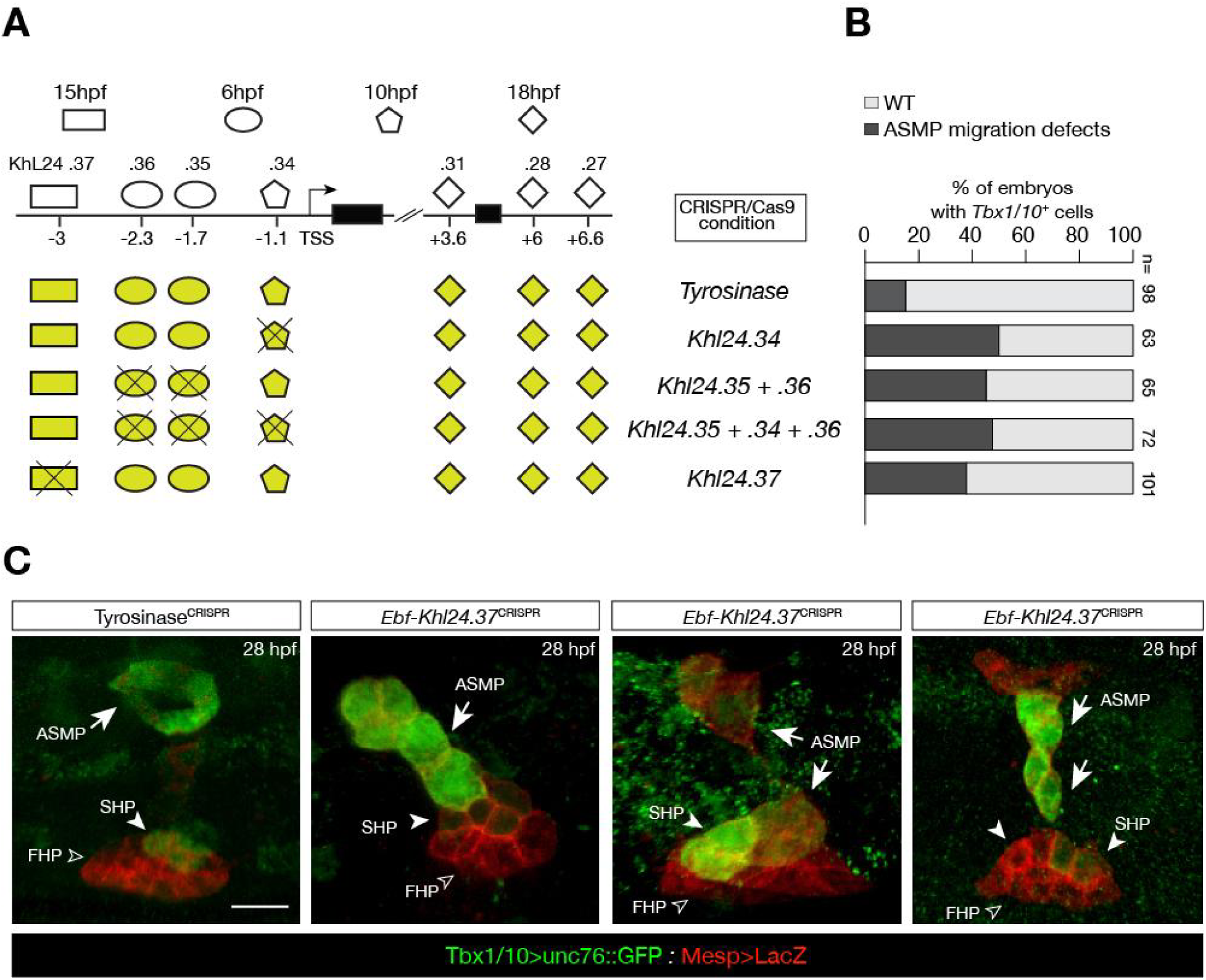
CRISPR/Cas9-mediated deletions on individual accessible elements upstream of *Ebf* caused phenotypic impact on pharyngeal muscle precursors morphogenesis. **(A)** Schematic representation of *Ebf cis*-regulatory elements targeted for CRISPR/Cas9-mediated deletions; the shapes of the distinct *cis*-regulatory elements are as in Figure 5C. **(B)** Proportions of larva halves showing GFP driven STVC-specific enhancer of *Tbx1/10* in indicated experimental conditions; all the treatments were significant versus Tyrosinase (Fisher exact test, p < 0.001; “n” is the total number of individual halves scored per condition.). **(C)** Example of an embryo at 28hpf showing GFP expression only in the ASM (solid arrowhead) and SHP (arrow) but not in the FHP (open arrowheads), where *Tbx1/10* enhancer is not active (Tyrosinase^CRISPR^, first panel on the left). Targeted deletions in *KhL24.37* peak induced ASMP cell migration defects. B7.5 lineage cells are labelled by Mesp>LacZ. Scale bar = 10 µm.

**Figure 5—figure supplement 4.**
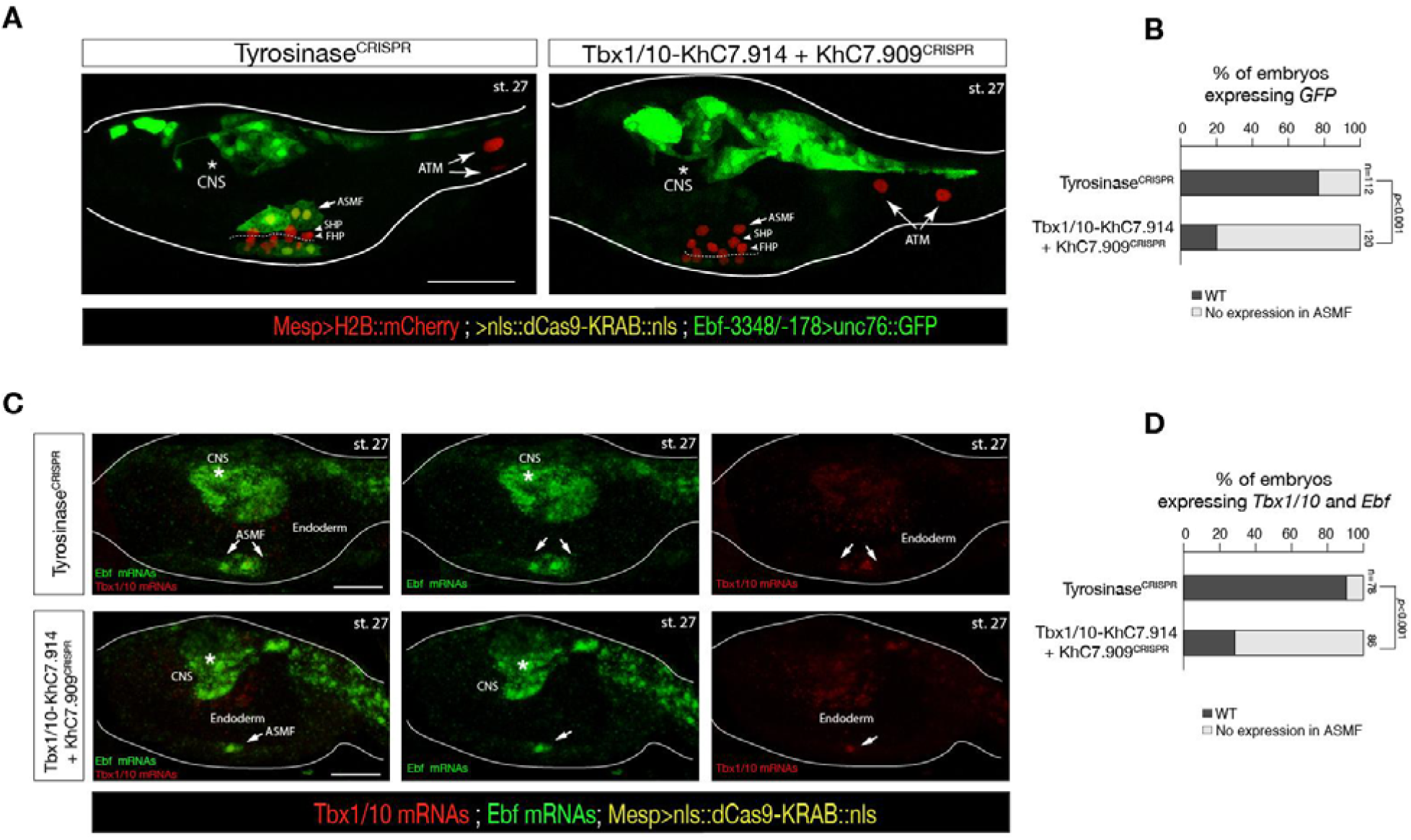
Intronic and distal enhancer accessibility in the *Tbx1/10* locus tested by dCas9-KRAB. **(A)** ASM-specific enhancer of *Ebf* (Ebf-3348/-178) (Wang et al., 2013) driven *in vivo* reporter expression (green) in embryos at stage 27 (Hotta et al., 2007) electroporated with Mesp>dCas9-KRAB along with single guide RNAs targeting *Tyrosinase* (control, left panel) as well as intronic (KhC7.914) and distal (KhC7.909) of *Tbx1/10* locus (right panel) as in Figure 5—figure supplement 2G. Nuclei of B7.5 lineage cells are labelled by Mesp>H2B::mCherry (red). White asterisk indicated central nervous system (CNS). Scale bar, 50 μm. **(B)** Proportions of larvae halves showing GFP expressed in the ASMFs in indicated experimental conditions (Fisher exact test, *p* < 0.001). **(C)** Double *in situ* hybridization of *Ebf* (green) and *Tbx1/10* (red) on embryos at stage 27 electroporated with Mesp>dCas9-KRAB along with single guide RNAs targeting Tyrosinase (control, left panel) as well as intronic (KhC7.914) and distal (KhC7.909) of Tbx1/10 locus. White asterisk indicated central nervous system (CNS). Scale bar, 20 μm. **(D)** Proportions of larvae halves expressing both *Tbx1/10* and *Ebf* in the ASMFs in indicated experimental conditions. Experiment performed in biological replicates. Total numbers of individual halves scored per condition are shown in ‘n=’. All the treatments were significant versus Control (*Tyrosinase*) (Fisher exact test, *p* < 0.001).

## Supplementary Tables

**Table S1.** Description of ATAC-seq dataset, with sample name, conditions, type of cells, number of mapped reads for all the biological replicates.

**Table S2.** Differentially FGF-MAPK-regulated genes (microarray) associated with differentially accessible peaks (ATAC-seq) in the pairwise comparison Mesp>Fgfr^DN^ vs. LacZ – 10hpf; genomic coordinates of each peak are reported.

**Table S3.** Known enhancers for TVC-specific genes characterized in previous studies; genomic coordinates of each peak are reported.

**Table S4.** Primer sequences used for cloning putative regulatory regions for functional enhancer assay.

**Table S5.** Primer sequences of single guide RNAs used to induce CRISPR/Cas9-mediated deletions in ATAC-seq peak as well as in Foxf (Foxf^CRISPR^) and to induce point mutations in Tbx1/10 STVC-specific enhancer; genomic coordinates and peak IDs are reported for each peak validated.

**Table S6.** ATM gene selected based on expression data deposited in ANISEED (https://www.aniseed.cnrs.fr).

**Table S7.** Differentially expressed genes from RNA-seq on FACS-purified cells following CRISPR/Cas9-induced loss of function of *Foxf*.

**Table S8.** Differentially accessible peaks from ATAC-seq on FACS-purified cells following CRISPR/Cas9-induced loss of function of Foxf (Foxf^CRISPR^ vs. Control^CRISPR^ at 10 hpf). Sheet1: closed in Foxf^CRISPR^ log_2_FC < −0.5 and FDR < 0.05); sheet2: open in Foxf^CRISPR^ log_2_FC > 0.5 and FDR < 0.05.

**Table S9.** Differentially accessible peaks from ATAC-seq, closed in the founder cells (LacZ 6hpf < LacZ 10hpf) and closed in Foxf^CRISPR^ vs. Control^CRISPR^ at 10 hpf.

**Table S10.** Differentially accessible peaks closed in Fgfr^DN^ at 10hpf and associated to de novo cardiac (sheet 1) and pharyngeal muscle (sheet 2) expressed genes.

**Table S11.** Differentially FGF-MAPK-regulated genes associated with differentially accessible peaks in MAPK activated −18hpf (sheet1) and MAPK inhibited −18hpf (sheet 2); genomic coordinates of each peak are reported.

**Table S12.** FGF-MAPK-regulated genes associated to ATAC-seq peaks that are non-differentially accessible in MAPK activated −18hpf and MAPK inhibited −18hpf.

**Table S13.** Accessible elements associated with de novo expressed heart and pharyngeal muscle markers into pre-accessible/primed or de novo accessible elements; genomic coordinates of each peak are reported.

## Supplementary Sequences

### FASTA sequences including the Foxf proteins (with their accession numbers) used for Figure 3—figure supplement 2

#### >NP_001071710.1 transcription factor protein [Ciona intestinalis]

MEVTGHHPGRVLSPVDVYQGRLNHTPQRVGHPQLSQQRSLPVLPHQYSLGNHHQPQLQHSHSIEQPPHHH QMPHPQPFQIQPNSHVLQCLQQVNKGEMPARPNSSEPALNAPLQPLHHSHIQRSPNGEAVPAMEELRNENA NNNNSAEGVMDGNTSAMSDGSNADLAVEHNLSIGAVSGAPETPPSTEEEGSDSTNGSGGKKQDNGGALG HRRPEKPPYSYIALIVMAIQSSPAKKLTLSEIYNFLQTRFEFFRGAYQGWKNSVRHNLSLNECFIKLPKGLGR PGKGHYWTIDPASEFMFEEGSFRRRPRGFRRKCQALKPYGIFGGGPGLIGHQGYGPPHEMFGAGGMPHGA MPPPSHHRPNLMGFDTAGMNAAAAASAHFFNGAAAAAVAANGITSPQTTSPTLPIPPKEPGTPHSPHNHSN PYSSNCAEVSTGTNTSVQVSPSMSAVANHPHYSPGAMFSWPGATSQSHGTYMRQNPPTPNGIPDPHTHAM MNGNVNGSVGQRMEYHHPFYATSRDSHLAYEAPTMKFKTECAMDPYSTNGMDRKPYPAMPTPIPVASGY SNGYYDTKSCAM

#### >Cisavi.CG.ENS81.R11.1287070-1294060.15296.p [Ciona savignyi]

MEVTGHHPGRVLSPVDVYQDRLNHTPQRVAHPQLSQQRSLPVQVHQYGIGSHHQPHLQHSHSEQPHLHH QMAHRQSFQIPPNSHVLQCLQQVNKDMPPRPNSSDPALSIPLQPLHHPHLQRSPNGDQVPALEELRHENAN NNNNSVDGGMDGNTSAMSDGSNADLSGEHSISIAPMSGAPETPPSTEEEGSDGNNGSGGKKTENGTLGHR RPEKPPYSYIALIVMAIQSSPAKKLTLSEIYNFLQTRFEFFRGAYQGWKNSVRHNLSLNECFIKLPKGLGRPG KGHYWTIDPASEFMFEEGSFRRRPRGFRRKCQALKPYGIFGGGPGLIGPQGYGHPHDMFGPSGMPHGAMPP PSHHRPNLMGFDPAGMNAAAAASAHFFNGAAAAAVAANGVTSPQTTSPTLPAVPKEPGTPHSPHNHPVTQ YPSNCAEVSSGTNSSIQVSPSMSAVANHHYSPGAMFSWPGTTSQSHGTYMRQNPGTTNGISDVHAHAMMN GNMNGSAGQRMEYPHPFYASPRESHLAYEAPAMKFKTECAMEPYPSNGMERKPYPAIPTPIPVPSGYSNGY YDTKSCAM

#### >XP_019628964.1 PREDICTED: forkhead box protein F1-A-like [Branchiostoma belcheri]

MTQEGTVENIQPTETATNTTLNGETTTATSATPCDESGKKKTNVGVRRHEKPPYSYIALIVMAIQSSATKRL TLSEIYQFLQQRFPFFRGPYQGWKNSVRHNLSLNECFIKLPKGLGRPGKGHYWTIDPASEFMFEEGSFRRRP RGFRRKCQALKPYGMLNNMVGSASMLGYDMLSGSQGTGTSPLQSLSPCSANSLNGMVNSTQGLMDTTGL MSYTSSMPGSLGGNLNQAPHGAAGVGGTGSAYMPSCSVGMSANDYGNGPNTVFTSQTTASLDNMSSGYT MSWAPAPTGHARYIKQQQDCGVGNTVSPTPAAVSPPMHAMTPVTQGETVQQQQQQHYLPQHSTQDVSD MAGNMSMRGLQSMADSSMCDRKPSYLLSSLQSSIHTGYYDTKCSM

#### >CBY09231.1 unnamed protein product [Oikopleura dioica]

MLDHESHKSLYGAHTDHIVGALNPAIVEKVSPSLSSLHEPLRTSSPVLEDYGVAQESISGHSTGINTSSNSSSP ENTTTDETSPAKDSTGTNPAATNGTDSKSTKDSSSKSARRPEKPPYSYIALIVMAIQSSPMKKLTLSEIYQFL QNKFEFFRGSYQGWKNSVRHNLSLNECFIKLPKGLGRPGKGHYWTIDPASEFMFEEGSFRRRPRGFRYRKC QALKPYSMLPGGGFSPYGVEMFPTGVPSSHGLASPASYNPAAQFGNFGLPQHGLPNALSKSPGLDGSTLSS TGIDTSSYADFTSQSSFFTQYPQYGFANMAGYGYPATSMTSTSMASSLPTPFPLSSSSPVNSTTSTLHRESTSP LNNPAHANQLEFDSSMKFKTEYGVTHTAQVAAGGKSPSIASAAGLSYPGALGAYSQYPQYSTPSSVLPDQ KPCIM

#### >XP_019628964.1 PREDICTED: forkhead box protein F1-A-like [Branchiostoma belcheri]

MTQEGTVENIQPTETATNTTLNGETTTATSATPCDESGKKKTNVGVRRHEKPPYSYIALIVMAIQSSATKRL TLSEIYQFLQQRFPFFRGPYQGWKNSVRHNLSLNECFIKLPKGLGRPGKGHYWTIDPASEFMFEEGSFRRRP RGFRRKCQALKPYGMLNNMVGSASMLGYDMLSGSQGTGTSPLQSLSPCSANSLNGMVNSTQGLMDTTGL MSYTSSMPGSLGGNLNQAPHGAAGVGGTGSAYMPSCSVGMSANDYGNGPNTVFTSQTTASLDNMSSGYT MSWAPAPTGHARYIKQQQDCGVGNTVSPTPAAVSPPMHAMTPVTQGETVQQQQQQHYLPQHSTQDVSD MAGNMSMRGLQSMADSSMCDRKPSYLLSSLQSSIHTGYYDTKCSM

#### >Boschl.CG.Botznik2013.chrUn.g03048.01.p [Botryllus schlosseri]

MLPQRQQSSPSSDTQPLPETPTDDTNINNNSNKIHDATVDESHNQESAGTGASVNGSETSTSTEEDGSDGGA AVSTKKGTGAHRRPEKPPYSYIALIVMAIQSSATKKLTLSEIYHFLQTRFEFFRGSYQGWKNSVRHNLSLNE CFIKLPKGLGRPGKGHYWTIDPASEFMFEEGSFRRRPRGFRRKCQALKPYGLFGGPASLVSHQGYPSHDMF GHTGPLHHGSMPPPRHQPGLLGFDPQGMNFLNGAAAAAAAANGMANPSSPNHPANSKVPGTPQTPPAVN PHYPTNTHCPNSAIDARTTQISPSSAAGMAVHNHPYSASMFNWSSASSHVAGGYMRHGPGNPVSPADHQV PIMNGMPPPGSRMDYHSLYAQARENHLPYDGKWSFVIEHVD

#### >Harore.CG.MTP2014.S48.g08804.01.p [Halocynthia roretzi]

MESGSCRNASVGVMSPNASESRLLPNSHPLSHTGPPPMHGQHNSPHSMPTIAHHRLSLNGPHLSVAPHPVIH GNAVESFHQAAMGGHMPHQQISPVAQQQHPNHHMLHISHSHDPLSSLHSLHYQQQQHQQLQRSQSHHMS TSLPHHNQPLDVGSSRPNSNEGLPHMSPHQRHQQLNNHDHSSRDSGMISHQGSPGENNRQVNSVQSVADN SANGNDDSNVNNSSNNLNNVAMSETAVEQDPEAENGVNSSFNAESETSTSTEEDGSDGGATVGSKKNNSG ANNTSTHRRPEKPPYSYIALIVMAIQSSTTKKLTLSEIYHFLQTRFEFFRGSYQGWKNSVRHNLSLNECFIKL PKGLGRPGKGHYWTIDPASEFMFEEGSFRRRPRGFRRKCQALKPYSIFGGPPGFMAPQGYPSHDMFPPSGS MPHGHMPPPRHQANLMGFDPSGVNFLNGAAAVVNGMASPTSPKIPGTPQSPNPSSNNLYPSPTSSSILSCAN SLIDTRTTQISTSASGMTMPNHHYSASMFNWPGAAAHAAGGYIRHGPLTTGSPSDNQPPHIMNGVPGGRM DYHSFYASAARDNYIPYEGKLCLNI*

#### >Moocci.CG.ELv1_2.S456097.g14882.01.p [Molgula occidentalis]

MIEMDGNLHCSSSLAIMSPTTHEMRGSLSQPLHHSIPSVQNQSAPSLQNQQHINPLQHQDQIPGQGGPMDA YRGRALSDNTHQGLHISHSHDALGLQQQMQYQHQHNALYRSQSMIPQPGVHLPPSVDSVVAPPRPGSNEIP HNISMERKTSQSSPISESRHPLSDTSLDDSNANNNSNNRTLSEDPEISADLNNTAASMTTGSEASTSTEEDGS DGGVAVNKKSGGSSNAAGHRRPEKPPYSYIALIVMAIQSSQTKKLTLSEIYNFLQSKFEFFRGSYQGWKNS VRHNLSLNECFIKLPKGLGRPGKGHYWTIDPASEFMFEEGSFRRRPRGFRRKCQALKPYGIFGGPPPGLMGP QSYPHPDMFPQVSGLPHGHMPPPRHQPNLMGFDPSGMGFLNGGAAPIASNRSPTSPLTPKPPGTPHSPSNHT GRLYGSPSAVTGPSPNDSRASQILHSPASMTTANPHYPSAMFSWPSPGPHPGTYIRHDATSTAASITETQNHF MNAAGSRIEHHPFYASADRANIAYEGKDGL

#### >phmamm.CG.MTP2014.S134.g03940.01.p [Phallusia mammillata]

MEMYLPHPAQMHSMKNSSQFLTADWDTMTSVGMKNWNISHTQVFPPIPTNPLSYNPQEPGEVGAFGVESNSSPNSDSRADEATTRWSKTTSKKKKIADKKYRRYEKPPYSYVGLIALAIQSSTNKMLKLSEILSRISTMFPFFKGEYQGWRDSVRHNLSQNKCFKKVLSDPYRPQSKGNYWTVDVNEIPAEKLKRQNTSVSRNVAPGYAYARDLNDIFDMSTGKLKVSNAVGSQHGFNTEAMDSNFAEHSSTSFNCISYDTTIACNDWNQHPAFTAVPTQSNTCKVAERKKDLTGTKQINMNEVSDISDGNENSSSAESCVPTDSSDDAGELGSKPSRSRSRRNKRKRALHLHHKRARPSCENQSPSPPSDYNRTKVQGNTKKADQPVPPNQKPFGDNNLAPKGEKSLGAVRSHLWSSIDQTVGPEGPPNITWTSSTPLNDVRPFFEPPSSQFDAHSSSQWSGDVTSIQTEPSATCDKTSQLFADQTDFLTEQASILPGRQGGDFSQPTVNQTLRPEFPHVAQTAQQHTFHGSSNLFAPPLNLPLDPGGSNFTHTSPHYLPAVEPIAYNLDKRYPSCEDPGLLRPGYTWPHSTGPFSDWVCNNFIPPSL*

#### >NP_001442.2 forkhead box protein F1 [Homo sapiens]

MSSAPEKQQPPHGGGGGGGGGGGAAMDPASSGPSKAKKTNAGIRRPEKPPYSYIALIVMAIQSSPTKRLTLSEIYQFLQSRFPFFRGSYQGWKNSVRHNLSLNECFIKLPKGLGRPGKGHYWTIDPASEFMFEEGSFRRRPRGFRRKCQALKPMYSMMNGLGFNHLPDTYGFQGSAGGLSCPPNSLALEGGLGMMNGHLPGNVDGMALPSHSVPHLPSNGGHSYMGGCGGAAAGEYPHHDSSVPASPLLPTGAGGVMEPHAVYSGSAAAWPPSASAALNSGASYIKQQPLSPCNPAANPLSGSLSTHSLEQPYLHQNSHNAPAELQGIPRYHSQSPSMCDRKEFVFSFNAMA SSSMHSAGGGSYYHQQVTYQDIKPCVM

